# Gut microbiota gate host exposure to cholinesterase inhibitors from dietary Solanums

**DOI:** 10.1101/2024.03.20.584512

**Authors:** Catherine S. Liou, Tijs Louwies, Mikhail Iakiviak, Jakub Rajniak, Pallavi P. Murugkar, Steven K. Higginbottom, Allison Weakley, Xiandong Meng, Phyu Htet, Ashley V. Cabrera, Marissa Jasper, Alex Dimas, EA Schulman, Michael A. Fischbach, Purna C. Kashyap, Elizabeth S. Sattely

## Abstract

Dietary plants are molecularly rich but the fates of these compounds post-ingestion and their implications for human health are largely unknown. Here, we systematically characterized the major chemical contributions of widely consumed Solanum species (nightshades) to the human metabolome. Using untargeted metabolomics, we found that a series of steroidal alkaloids resulting from glycoalkaloids tomatine, solanine, and chaconine are dominant diet-derived compounds in systemic circulation following ingestion of tomato and potato. By comparing serum and tissue metabolomes of colonized and microbiome-depleted mice, we determined that the gut microbiota modifies these compounds extensively, altering their absorption and gating host exposure. By screening the metabolic products in human urine and stool samples, we established that steroidal glycoalkaloid metabolism varies inter-individually in a population. Furthermore, using a collection of representative human commensal type strains, we found that a limited set of strains is responsible for steroidal glycoalkaloid metabolism, with the chemical output of a community determined by its strain-level composition. These findings enabled the rational design of complex synthetic microbial communities that controlled host exposure to steroidal alkaloid metabolites *in vivo*. Importantly, microbial metabolism of Solanum metabolites alters their acetylcholinesterase inhibition *in vitro* and gut motility *in vivo*. Our study provides insights into the molecular mechanisms of a diet-microbiome interactions and the effects of dietary metabolites on host physiology.

## INTRODUCTION

Plants contribute incredible molecular diversity to diet, providing not only carbohydrates, proteins, fats, and vitamins but also specialized small molecules. These secondary metabolites, produced by plants for defense, environmental stress tolerance, and nutrient acquisition, are estimated to be ingested in significant quantities of at least 50 milligrams per day (Murphy et al., 2012; Tennant et al., 2014) and have been associated with numerous pharmacological effects (Butelli et al., 2008; Fahey et al., 1997; Martin et al., 2013). The immense chemical complexity of plants outmatches our understanding of the major molecular inputs from diet, which obscures connections between dietary plant consumption and human health. Further exacerbating this challenge is metabolite transformation after ingestion which vastly expands the array of diet-derived metabolites in the human metabolome.

In addition to host digestive processes, dietary compounds can be transformed by commensal bacteria that reside in the gut. Metabolism by the gut microbiota has an outsized impact on access to nutrients from diet (Turnbaugh et al., 2006) and we are only beginning to explore the specific molecular contributions by bacterial metabolism to the host metabolome (Marcobal et al., 2013). Biotransformation by the gut microbiota can alter compound bioactivity and bioavailability; furthermore, microbiota-specific metabolism can vary inter-individually, resulting in divergent molecular exposures and health outcomes from a common dietary input (Fuhrman et al., 2008; Koeth et al., 2013). Elucidation of bioactive compounds from diet, their transformation by gut microbiota, and their ultimate impacts on human health will not only expand our ability to take advantage of the benefits of dietary plants but also provide valuable insight into the molecular interactions that exist at the interface of the plant, bacterial, and animal kingdoms.

Nightshades from the family Solanaceae, including potatoes, tomatoes, and eggplants, are of particular interest among dietary plants. Solanums are among the most abundantly consumed plants in the United States (Economic Research Service (ERS), U.S. Department of Agriculture (USDA), n.d.) and are rich in specialized compounds, including carotenoids, steroidal glycoalkaloids, and flavonoids, that have been studied in many pharmacological contexts (Chaudhary et al., 2018; Schieber and Saldaña, 2009). Diets rich in tomatoes and potatoes have been associated with different physiological outcomes (Giovannucci, 1999); tomato ingestion, for example, has been shown to reduce serum LDL oxidation (Agarwal and Rao, 1998) and decrease the incidence of cardiovascular disease in a cohort of women (Sesso et al., 2003). However, the specific compounds that drive these pharmacological effects are still under active investigation.

In this study, we use dietary Solanums as a model to explore the major molecular inputs from a commonly consumed food, the role of gut microbiota in gating host exposure to these inputs, and the impacts of commensal-specific metabolism on compound bioavailability and bioactivity. We performed untargeted metabolomics in animal models and identified steroidal alkaloids as the major diet-derived compounds in host circulation following Solanum ingestion. We mapped the metabolic fates of dietary steroidal glycoalkaloids and reconstituted the production of these metabolites *in vitro* and *ex vivo* using a combination of gut commensal strains and human liver enzymes. These results reveal a paradigm where specific bacterial strains are responsible for key steps in compound metabolism, yielding metabolite and bioactivity profiles that are unique to individual community compositions. We further show that unmetabolized steroidal glycoalkaloids from potato increase gut motility *in vivo,* an effect that is not produced by steroidal alkaloid aglycones, the metabolized form of this molecule. Biotransformation by the gut microbiota is a layer of metabolism that can profoundly impact interactions between diet and the host and must be characterized to fully understand the physiological effects of a dietary molecule.

## RESULTS

### Major molecular inputs from dietary Solanums

We focused on Solanums, including potatoes and tomatoes, as abundantly consumed plants rich in specialized compounds. To explore the major molecular inputs derived from ingestion of these plants, we performed untargeted metabolomics for unbiased identification of diet-derived compounds encountered in systemic circulation. We used liquid chromatography coupled to mass spectrometry (LC-MS) to profile serum from Swiss Webster mice provided with *ad libitum* access to ripe Microtom tomatoes in addition to a standard diet for 15 days (**Fig 1A**). For detection of a broad range of molecule classes in serum, analyses were performed using both hydrophilic interaction liquid chromatography (HILIC) for the retention of more polar compounds and reverse phase chromatography for the retention of nonpolar compounds. To identify tomato-associated compounds in systemic circulation, mass features were filtered for those that significantly accumulated to at least three-fold greater abundance in samples from mice receiving tomatoes than in mice that did not receive tomatoes (**Fig S1A**). Many of these molecules were absent in the tomato supplement and were therefore likely derived either from the metabolism of tomato compounds or endogenous mouse compounds produced in response to tomato consumption (**Fig 1A**). We identified one of these metabolite to be trimethylamine N-oxide (TMAO), known to be produced by intestinal bacteria from trimethylamine containing precursors like choline, betaine, and carnitine (Simó and García-Cañas, 2020). Serum TMAO levels were increased 2.5-fold by tomato supplementation (**Fig S1B**), an unexpected result considering the association of circulating TMAO with increased risk for cardiovascular disease (Zhu et al., 2016). This finding highlights the strength of an untargeted approach in identifying unexpected compounds resulting from a dietary intervention.

**Figure 1.**
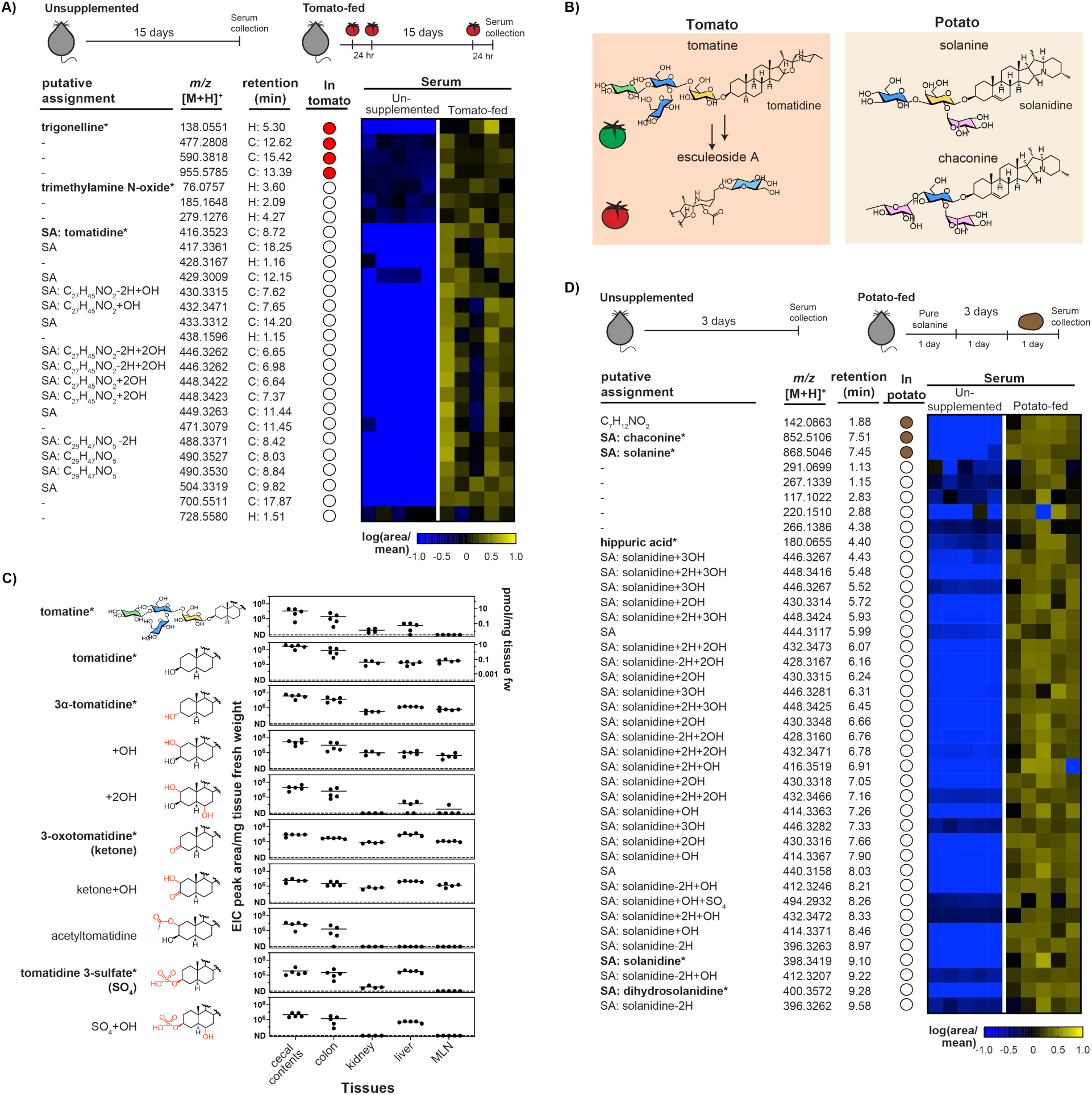
Major molecular inputs from dietary Solanums. A) Major mass features in mouse serum resulting from tomato intervention (schematic above) identified using untargeted LC-MS metabolomics in positive ionization mode with reversed-phase (C: C18) and normal phase (H: HILIC) chromatography. Abundances for differently accumulating mass features in each individual sample (n=5 per treatment) are shown in the heat map. Values shown are extracted ion chromatogram (EIC) peak areas normalized to the mean peak area across individuals. Mass features also observed in the tomato supplement are marked with a filled circle. Putative compound assignments are provided, with SA denoting putative steroidal alkaloid (SA) compounds. Assignments with an asterisk (*) have been confirmed by retention time and MS/MS matches to a chemical standard. B) The major steroidal glycoalkaloids (SGAs) present in tomato and potato. Glycone monomers are colored by identity (blue: glucose, yellow: galactose, green: xylose, pink: rhamnose) C) Accumulation of SGA-derived metabolites in mouse tissue following dietary intervention with purified tomatine (15 μM) in drinking water. Values shown are the mean EIC peak areas normalized to tissue fresh weight with individual replicates overlaid (n=5). Corresponding molar quantities for compounds with commercial standards are indicated on the right axes. Structures that have been characterized by comparison to a chemical standard or reproduced *in vitro* using enzymatic reactions are marked with an asterisk (*). Other structures are hypothesized based on *m/z* and MS/MS spectra; the positions of hydroxylations is not known. For molecules with multiple detected isomers (+OH and +2OH aglycones), the sum of peak areas is shown. ND = not detectable.

When subjected to tandem mass spectrometry (MS/MS) for structural characterization, fourteen of the 28 tomato-associated compounds found in serum yielded mass fragmentation patterns characteristic of steroidal alkaloids (SAs) (**Fig S1C**). Steroidal alkaloids are nitrogen containing compounds derived from cholesterol and accumulate highly in Solanums as glycosides known as steroidal glycoalkaloids (SGAs). Dietary Solanums accumulate species-specific SGAs with structural variations in both the steroidal aglycone and the glycone (**Fig 1B**). Tomato fruits produce the SGA tomatine which is converted *in planta* to esculeoside A during fruit ripening. We found that the major tomato-associated compounds in serum included tomatidine and esculeogenin A, the aglycones derived from tomatine and esculeoside A, respectively, as well as hydroxylated versions of these aglycones (**Fig 1A**). In contrast to the abundance of SGAs and the absence of detectable SA aglycones in tomato, we observed primarily aglycones accumulating in the serum. These results suggest that SGAs undergo significant metabolism following consumption.

### Tomato steroidal glycoalkaloids are deglycosylated and further modified after ingestion

To track the metabolic fates of SGAs independently of the other dietary components in tomatoes, we next provided Swiss Webster mice with pure tomatine in drinking water. Reasoning that metabolites may accumulate more highly in certain tissues than in systemic circulation, we used untargeted metabolomics to characterize the major tomatine metabolites in tissues and therefore have the potential to interact with tissue-specific physiology. We surveyed a panel of tissues for steroidal alkaloid content, including two tissues along the digestive tract (colonic tissue and luminal contents from the cecum), another two that are first-pass tissues for nutrient absorption (mesenteric lymph nodes, MLNs, for molecules transported by the lymphatic system and liver for molecules transported by the portal vein), and kidney, based on previous studies that have tracked tissue specific accumulation of SGAs from potato (Norred et al., 1976).

In addition to the hydroxylated SA metabolites previously observed in systemic circulation, we were able to detect several additional aglycone metabolites in the panel of tissues (**Fig 1C**). We characterized select compounds by retention time and MS/MS spectrum match to synthesized standards, confirming the structures of tomatidine oxidized at the C3 alcohol to yield the ketone 3-oxotomatidine (**Fig S2A**) and sulfated to form tomatidine 3-sulfate (**Fig S2B**). Other observed aglycone metabolites were putatively assigned through MS/MS analysis to be the 3αOH epimer and acetylated analogues of tomatidine, as well as hydroxylated versions of these modified tomatidine compounds (**Supplemental Table 1**). These assignments are supported by recent studies reporting steroidal alkaloid metabolites in pig, mouse, and human plasma following dietary intervention with tomato (Cichon et al., 2017; Do et al., 2024; Sholola et al., 2025). The elucidation of these metabolites reveals that, beyond deglycosylation, extensive structural modifications are enacted on SGAs following ingestion.

Because only glycoside and not aglycone SAs were present in tomato, we wanted to determine how the diversity of aglycones in serum and tissues were formed. Tomatine and many of the SA aglycone metabolites accumulated primarily in the cecal contents and the colon (**Fig 1C**). Select aglycones, however, including 3-oxotomatidine, hydroxylated tomatidine, and tomatidine sulfate were also abundant in the liver; accordingly, we hypothesized that these compounds could result from hepatic metabolism. To test this hypothesis, we incubated human liver protein extracts with tomatine and tomatidine and analyzed the products of metabolism using LC-MS. While human liver enzymes did not modify the glycoside tomatine *in vitro*, we observed conversion of the aglycone tomatidine to both tomatidine sulfate (**Fig S2C**) and hydroxytomatidine (**Fig S2D**). We did not detect oxidation of tomatidine to the ketone 3-oxotomatidine by the human liver enzymes. However, we could observe the reverse reaction; when 3-oxotomatidine was supplied as a substrate, the human liver enzymes non-selectively reduced the ketone to 3βOH tomatidine and its 3αOH epimer (**Fig S2E**). This conversion suggests that the putative 3αOH tomatidine epimer observed in mouse tissue following tomatine dietary intervention may be produced from the reduction of 3-oxotomatidine by hepatic metabolism. Despite our recapitulation of select SA metabolites from tomatidine, key metabolic transformations responsible for the SAs found in serum, including SGA deglycosylation and any reactions acting directly on the glycosides, were not promoted by human liver enzymes *in vitro*. These data suggest that hepatic metabolism contributes to aglycone modification, generating sulfated and hydroxylated aglycone analogues, but has a limited role in directly metabolizing dietary SGAs.

### Potato steroidal glycoalkaloids from potato undergo similar metabolism as tomato SGAs

To explore the molecular inputs from other dietary Solanums and track their metabolic fates, we provided Swiss Webster mice with a single dose of the potato SGA solanine, followed by diet supplemented with 10% potato peel. Using a similar approach as for the tomato dietary supplement, we performed untargeted metabolomics to identify the major diet-derived metabolites that accumulated in systemic circulation (**Fig S3A**). We found that dietary intervention with potatoes significantly increased serum levels of several compounds (**Fig 1D**), including hippuric acid (**Fig S3B**), potentially derived from phenolic compounds abundant in potato (Krupp et al., 2012). The majority of the compounds that accumulated in serum with potato consumption, however, had MS/MS spectra characteristic of solanidine-based steroidal alkaloids (**Fig S3C**). These metabolites included the major SGAs solanine and chaconine present in potato (**Fig 1B**), as well as the corresponding aglycone solanidine. Reminiscent of the metabolic fates of tomatine, we observed compounds derived from the aglycone solanidine, structurally modified by hydroxylation, sulfation, and oxidation. We additionally observed hydrogenation of the Δ5,6 double bond of solanidine to form 5α and 5β isomers of dihydrosolanidine, confirmed by MS/MS spectrum and retention time matching to a synthesized standard (**Fig S4A**). Dihydrosolanidine appears to undergo similar downstream modifications as the unsaturated solanidine, with hydroxylated and sulfated analogues of dihydrosolanidine detected in serum.

To investigate the origins of these serum metabolites, we incubated solanine, chaconine, and solanidine with human liver proteins *in vitro*. As was the case with the tomato SGA, human liver proteins did not directly act on the glycosides solanine and chaconine. We recapitulated conversion of the aglycone solanidine to sulfated (**Fig S4B**) and hydroxylated (**Fig S4C**) solanidine using the liver enzymes. However, other modified aglycone metabolites, including keto-and saturated analogues of solanidine, were not produced. These data show that, like tomato SGAs, potato SGAs undergo extensive modifications including deglycosylation, many of which occur independently of hepatic metabolism.

Taken together, we found that steroidal alkaloid aglycones represent a substantial portion of the detectable diet-derived metabolites in systemic circulation following ingestion of tomatoes and potatoes. These data support other studies that have noted steroidal alkaloids and their metabolites to be small molecule biomarkers characteristic of tomato consumption (Cichon et al., 2017; Do et al., 2024; Dzakovich et al., 2024; Hövelmann et al., 2020; Sholola et al., 2025). We further characterized the SA metabolites observed *in vivo* to uncover modifications that occur to SGAs following ingestion, including oxidations and hydrogenations. While some modification of SA aglycones could be attributed to hepatic metabolism, the source of SGA metabolism and deglycosylation remained unclear.

### Gut microbiota gate the formation of steroidal alkaloid aglycones *in vivo*

As we could not detect aglycone formation from SGAs by human metabolic enzymes *in vitro*, we next wanted to explore the mechanisms through which SGAs are deglycosylated *in vivo*. Considering the abundance of SA aglycones in the digestive tract (**Fig 1C**) and the established capabilities of gut commensals in polysaccharide hydrolysis (Cockburn and Koropatkin, 2016), we hypothesized that gut microbiota could play a role in SGA metabolism. Fecal pellets from conventional mice and mice treated with broad spectrum antibiotics (Rakoff-Nahoum et al., 2004) were incubated with exogenously added tomatine. We observed formation of the aglycone tomatidine in the feces of conventional mice, but not in feces from mice receiving antibiotics (**Fig S5A**), indicating that native murine commensal bacteria are capable of tomatine deglycosylation.

In addition to gut microbiota, host digestive processes including acid hydrolysis in the stomach and intestinal glycosidases may also contribute to SGA deglycosylation. To assess the contributions of gut microbiota towards SGA deglycosylation *in vivo*, we provided C57BL/6J mice with diet supplemented with 10% tomato concurrently with antibiotics treatment (**Fig 2A**). The aglycone tomatidine was absent from the serum (**Fig 2B**) and tissues (**Fig S5D**) in mice receiving antibiotics. Other modified aglycone metabolites, including 3-oxotomatidine (**Fig 2C**) and tomatidine sulfate (**Fig S5D**), were similarly depleted by antibiotics treatment. These data show that hydrolysis of the lycotetraose glycone in tomatine is performed primarily by gut microbiota and not by host digestion. Furthermore, microbiome-mediated metabolism gates formation of the diverse modified aglycones that accumulate across host tissues.

**Figure 2.**
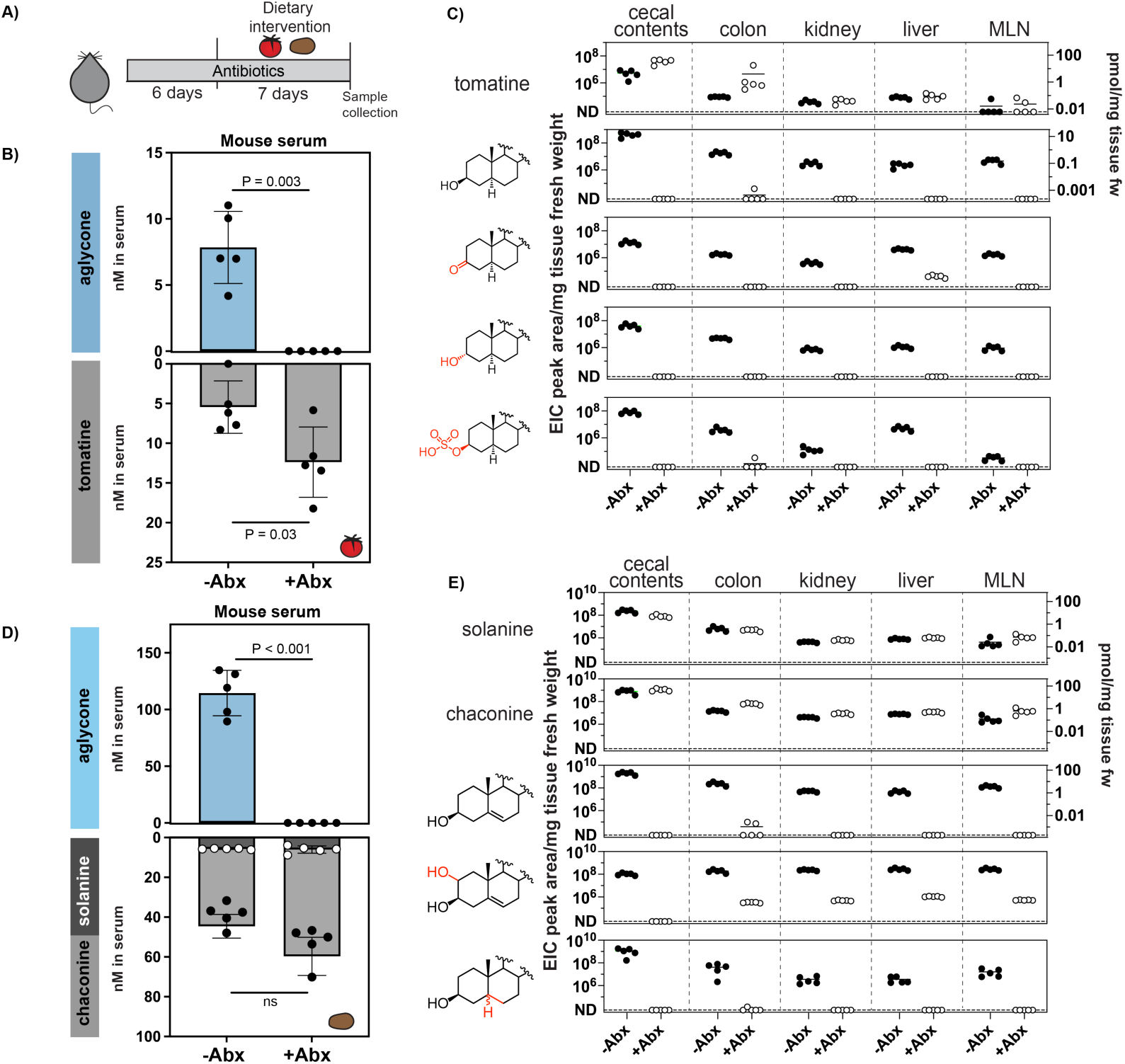
Gut microbiota gate host exposure to steroidal alkaloid aglycones. A) Schematic of concurrent antibiotic treatment and Solanum dietary intervention in C57BL/6J mice. Animals were provided AIN93G diet supplemented with 10% w/w lyophilized tomato or 5% w/w lyophilized potato peels for a week. Tomato-supplemented diet provided about 42 μmol/kg diet tomatine and 16.5 μmol/kg diet esculeoside A. Potato-supplemented diet provided about 132 μmol/kg diet chaconine and 53 μmol/kg diet solanine. B) Tomatine and tomatidine (aglycone) concentrations in serum (nM) following a 10% tomato dietary intervention in C57BL/6J mice receiving water (-Abx) or antibiotics (+Abx). Values shown are the mean±SD for each group, with individual replicates overlaid (n=5). P values were determined using a two-tailed t test. ns = not significant (P>0.05). C) Accumulation of a subset of tomatine-derived metabolites in tissue following dietary intervention with 10% tomato diet in C57BL/6J mice receiving water (filled circles) or antibiotics solution (unfilled circles). Values shown are mean EIC peak areas normalized to tissue fresh weight (fw) with individual replicates overlaid (n=5). Molar quantities for compounds with available commercial standards are indicated on the right axes. For molecules with multiple detected isomers (+OH and +2OH aglycones), the sum of peak areas is shown. ND = not detectable. D) Chaconine, solanine, and solanidine (aglycone) concentrations in serum (nM) following a 5% potato dietary intervention in C57BL/6J mice receiving water or antibiotics. Values shown are the mean±SD for each treatment group, with individual replicates overlaid (n=5). P values determined using a two-tailed t test. ns = not significant (P>0.05). E) Accumulation of a selection of glycoalkaloid-derived metabolites in tissues following dietary intervention with 5% potato diet in C57BL/6J mice receiving water (filled circles) or antibiotics solution (unfilled circles). Values shown are mean EIC peak areas normalized to tissue fresh weight with individual replicates overlaid (n=5). Molar quantities for compounds with commercial standards are indicated on the right axes where applicable. For molecules with multiple detected isomers (+OH and +2OH aglycones), the sum of peak areas is shown. ND = not detectable.

Esculeoside A, the major tomato SGA produced from tomatine during fruit ripening, was present alongside tomatine in the tomato dietary intervention. Though esculeoside A is structurally differentiated from tomatine by hydroxylation, acetylation, F ring inversion, and C27 glucosylation (Nohara et al., 2010) (**Fig 1B**), we observed similar microbiota-exclusive removal of the C3 conjugated tetraose from esculeoside A in the cecal contents of mice fed tomatoes (**Fig S5B**). Hydrolysis of the C27 glucose of esculeoside A, however, did not appear to be solely microbiota mediated as we were able to observe de-glucosylated esculeoside A in the cecal contents of antibiotics-treated mice. Though host digestion was able to hydrolyze the C27 glucose from esculeoside A, formation of the aglycone esculeogenin A through subsequent C3 lycotetraose hydrolysis was microbiome-dependent and completely absent in antibiotics-treated mice. These results indicate that the C3 lycotetraose is resistant to host metabolism but is readily liberated by microbial glycosidases. Notably, we did not detect any intermediates of tomatine or esculeoside A with partially metabolized C3 glycones, suggesting that the microbiota hydrolyze the SGA glycone as a single tetraose unit as opposed to sequential hydrolysis of sugar monomers and dimers.

Because the potato SGAs are structurally distinct to tomato SGAs, we next explored whether the gut microbiota is similarly necessary for solanine and chaconine deglycosylation. C57BL/6J mice concurrently receiving antibiotics and a diet containing potato peels were deficient in the aglycone solanidine, both in systemic circulation (**Fig 2D**) and tissues that highly accumulated SGAs (**Fig 2E**). These animals were also deficient in aglycones modified by saturation, hydroxylation, and sulfation (**Fig S5E**). While partially deglycosylated intermediates of tomatine were not observed *in vivo*, we detected intermediates of solanine and chaconine with one or two monomers hydrolyzed from the glycone in cecal contents of mice after potato dietary intervention (**Fig S5C**). These partially deglycosylated potato SGAs were only present in mice that did not receive antibiotics, indicating that potato SGA glycones show similar resistance to host digestion as the tomatine glycone.

Taken together, we found that SGAs are resistant to host digestion *in vivo*, requiring metabolism by the gut microbiota. Fomation of the modified aglycones detected across host tissue are preceded and gated by microbial deglycosylation, indicating that microbial metabolism determines host exposure to diverse SGA metabolites. Moreover, these results reveal that antibiotic alteration of the gut microbiota can significantly alter specific processing of drug-like molecules in the diet and, as a consequence, host exposure to the derived metabolites.

### Inter-individual variation in SGA metabolism between human donors

A key implication of SGAs being metabolized exclusively by gut microbiota is potential variation in metabolism between individuals, with unique bacterial communities generating distinct metabolite profiles from a common molecular input from diet. After establishing the necessity of gut microbiota in metabolizing SGAs *in vivo*, we next explored how these compounds might be metabolized differently between individuals. Having previous characterized the major SA metabolites produced by native mouse microbiota, we began by identifying the major compounds produced from SGAs by human gut microbial communities. Stool samples (Carter et al., 2025) from thirty human donors were incubated *ex vivo* with the major SGAs from tomato, potato, and eggplant (**Fig S7A**) and assayed for product formation by LC-MS. MS fragmentation was used to identify mass features containing characteristic steroidal substructures (**Fig S6A**). Using this untargeted approach, we identified not only previously characterized compounds like sulfated and oxidized aglycones, but also those had not been previously observed in our analyses of mouse samples. One such metabolite was tomatidine and solasodine conjugated to a putative phosphoethanolamine moiety (**Fig S6B**), which we characterized by LC-MS/MS and hydrogen-deuterium exchange LC-MS to lend support to our proposed structural assignment (**Fig S6C**).

Comparing glycoalkaloid conversion by thirty human donor samples, we found that SGA metabolism was not universal, with some samples extensively deglycosylating SGAs and others inactive (**Fig 3A**). Leveraging the shared structural features of the eggplant and potato SGAs, we noted that SGA metabolism appeared to correlate with identity of the glycone. Solanine and the eggplant SGA solasonine share the same solatriose glycone and were metabolized by a similar subset of samples. In the same manner, another subset of donors was able to metabolize both chaconine and the eggplant SGA solamargine, which share a chacotriose sugar.

**Figure 3.**
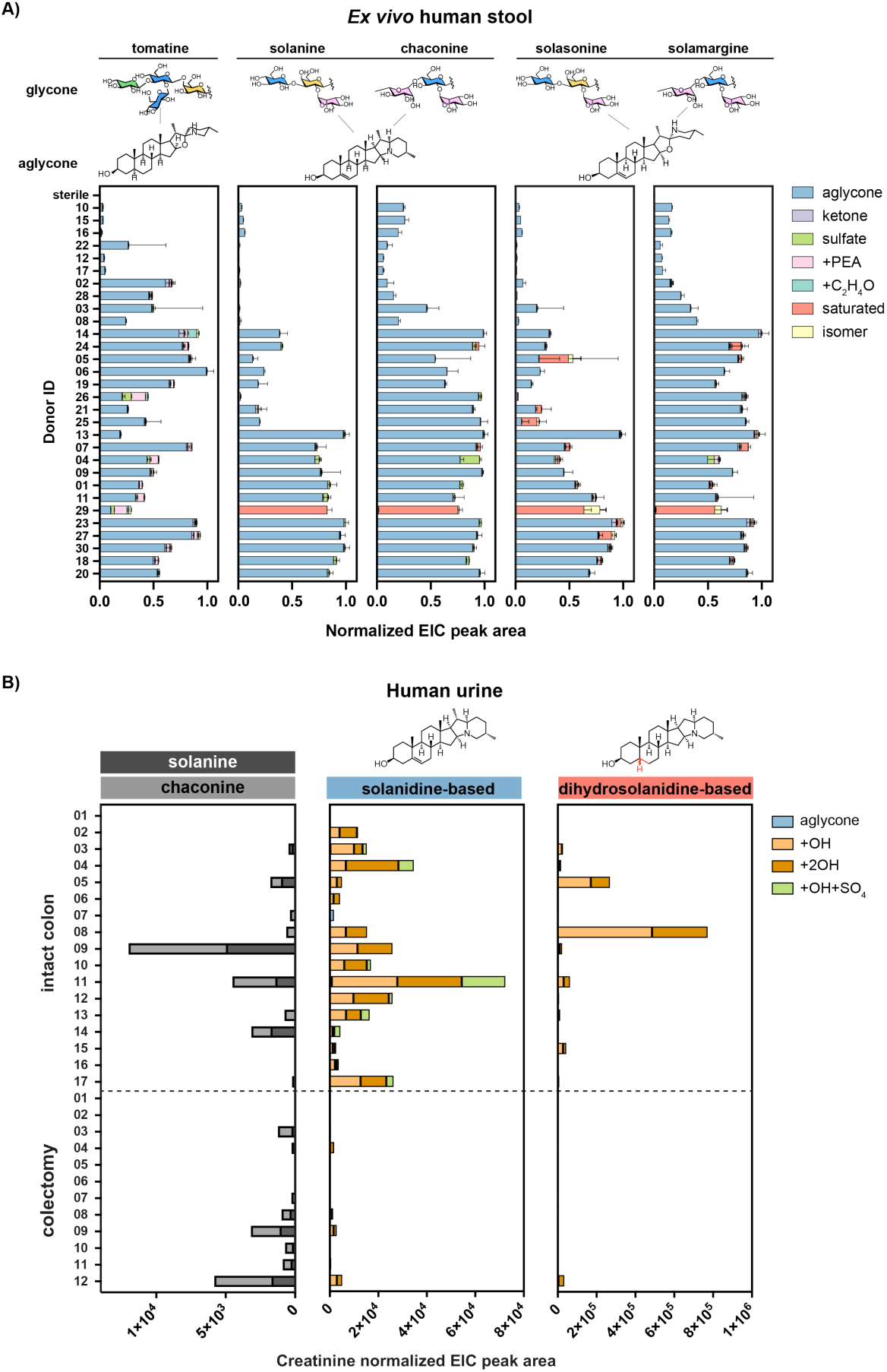
Inter-individual variation in steroidal glycoalkaloid metabolism between human donors. A) *Ex vivo* metabolism of tomato, potato, and eggplant SGAs by human stool samples from 30 donors. Stool samples were incubated with SGA substrate (25 μM) in SAAC minimal medium for 48 hours. EIC peak areas of the product aglycones are normalized to the maximum sum of product peak areas registered across all donors. Values shown are the mean±SD of three replicate incubations per donor and substrate combination. Colors indicate different aglycone products (ketone: oxidation at C3OH, sulfate: sulfation at C3OH, PEA: phosphoethanolamine, saturated: Δ5,6 hydrogenation, isomer: 3αOH epimer). B) Glycoalkaloid (solanine and chaconine) and aglycones (solanidine- and saturated solanidine-derived) detected in urine samples from human colectomy patients and donors with intact colons. Compound abundance is shown as EIC peak area normalized to the creatinine content in each sample. Values shown are for individual donor samples. Colors indicate different aglycone products observed.

In addition to differences in extent of SGA deglycosylation, we also observed inter-individual variation in the types of aglycones produced. Saturated analogues of solanidine and solasodine were predominant in only one out of the thirty samples surveyed, suggesting that the hydrogenation of the Δ5,6 double bond in these steroidal alkaloids is potentially a rare modification. Modifications that appeared more ubiquitous across the donors included phosphoethanolamine conjugation and sulfation.

To investigate inter-individual variation in aglycone modification independently of SGA deglycosylation, we directly incubated the stool samples with the aglycones tomatidine, solanidine, and solasodine. We noted that some samples that were inactive on glycosylated substrates were nonetheless active when supplied with the corresponding aglycone. For example, sample 26 did not significantly metabolize either solanine or solasonine but still produced abundant sulfonated aglycone when incubated directly with solanidine and solasodine (**Fig S7B**), respectively. These results suggest that SGA deglycosylation and aglycone modification are independent metabolic steps and may be performed by different bacterial strains in the community.

To further explore the relevance of this metabolism to humans *in vivo*, we analyzed urine samples collected from colectomy patients lacking an intact colon and therefore deficient in colonic microbiota (Mair et al., 2018). Even without a specified dietary intervention, we were able to detect potato SGAs solanine and chaconine co-occurring in the majority of urine samples, indicative of recent potato consumption (**Fig 3B**). Similarly to antibiotics treated mice, urine from colectomy patients were largely depleted in all SA aglycones compared to samples from donors with intact colons, showing that gut microbial metabolism is necessary for SA aglycone formation in humans. Furthermore, the presence of these molecules in human urine indicate that readily detectable quantities of steroidal alkaloids accumulate in systemic circulation in the absence of a specific dietary intervention, highlighting the abundance of these Solanum molecules in commonly consumed foods.

We observed two major classes of aglycones derived from potato SGAs in human urine, based either on solanidine or its Δ5,6 saturated analogue dihydrosolanidine (**Fig 3B**). While solanidine-based aglycones were detectable in most of the donors with intact colons, dihydrosolanidine-based aglycones were abundant only in a few samples, a pattern which corroborates the rarity of Δ5,6 saturation performed by human stool samples *ex vivo* (**Fig 3A**). In contrast to the products of *ex vivo* metabolism by human stool, the vast majority of the SA metabolites detected in human urine were hydroxylated aglycones. With *in vitro* evidence that hepatic enzymes perform this metabolism (**Fig S4C**), we propose that gut microbiota-generated aglycones are absorbed and undergo hydroxylation in the liver *in vivo*.

Taken together, we observe significant variation in both the extent of conversion and product profiles deriving from SGA metabolism between different human samples. This variation indicates that individuals may be exposed to different compounds from a common dietary Solanum input, depending on compositions of individual gut microbial communities.

### Microbial community composition determines SGA metabolic fate

Having noted significant inter-individual variation in SGA processing in human stool and urine samples, we next aimed to investigate how microbiota metabolism of dietary compounds is distributed among specific human commensal strains. We individually cultured 116 human commensal type strains selected to span diverse phyla and represent those that are abundant and prevalent in a Western microbiome (Cheng et al., 2022) (**Fig S8, Supplemental Table 2**). These type strains were cultured in rich medium supplemented with potato, tomato, and eggplant SGAs and assayed for compound metabolism using LC-MS (**Fig 4A**). We found that SGA deglycosylation was limited to a subset of the strains surveyed. For example, tomatine deglycosylation was detected in seven out of the 116 type strains, including *Roseburia intestinalis* L1-82, *Bacteroides uniformis* ATCC 8492, and strains from the genus *Parabacteroides*. Some strains were found to act broadly on the panel of SGAs tested; *Bacteroides* sp. 3_1_19, for example, was capable of metabolizing all five SGAs provided. Other strains yet were found to perform substrate-specific deglycosylation, such as *Bacteroides uniformis* ATCC 8492, which acted only on tomatine, and *Ruminococcus gnavus* ATCC 29149, which acted only on the SGAs with a chacotriose glycone, chaconine and solamargine. These data show that SGA deglycosylation is a substrate-specific transformation, likely involving strain-specific sets of polysaccharide metabolizing enzymes for select metabolism of dietary glycans.

**Figure 4.**
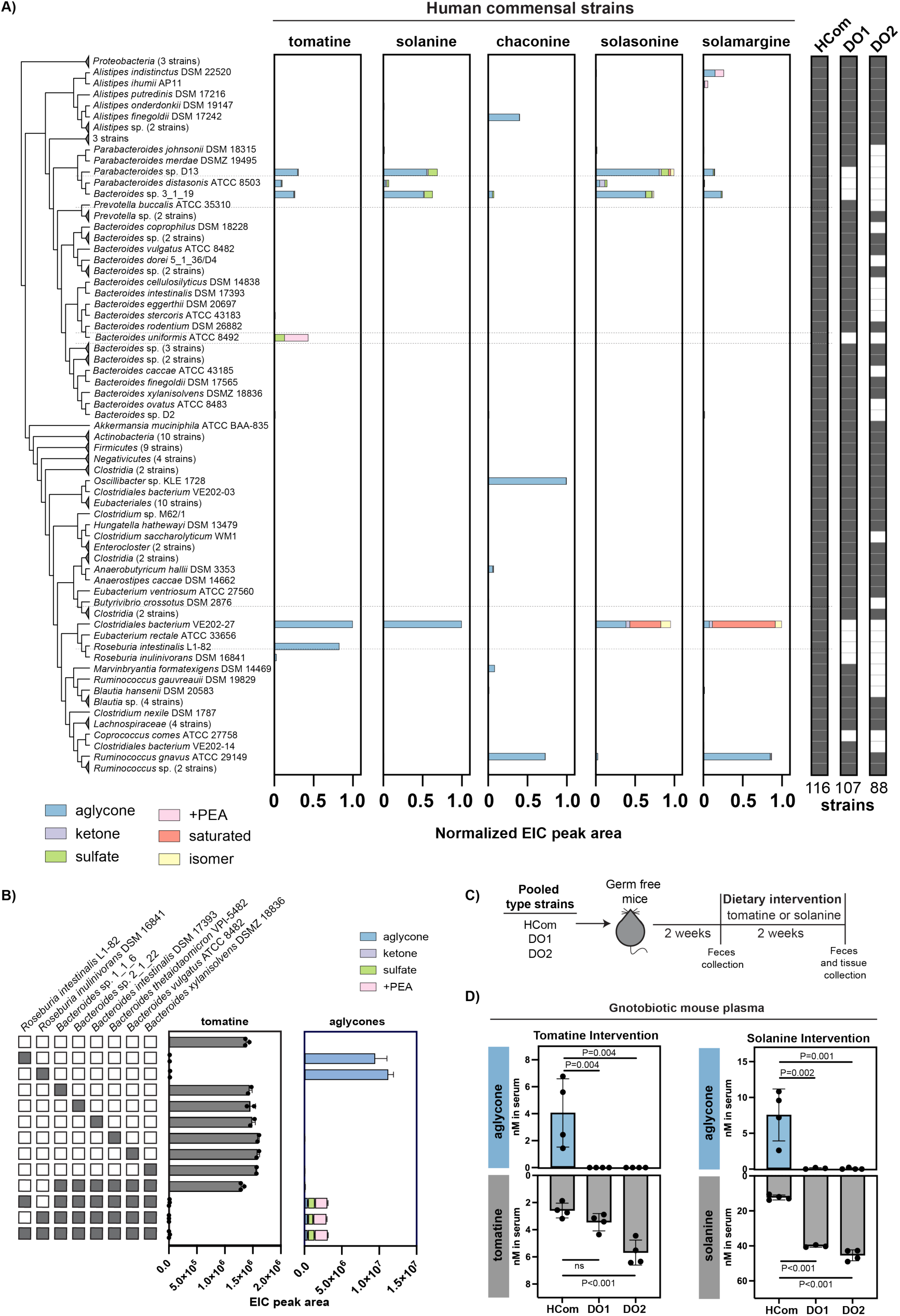
Microbial community composition determines SGA metabolic fate. A) Metabolism of tomato, potato, and eggplant SGAs by human commensal type strains after 48 hours of growth in Mega medium supplemented with SGA substrate (25 μM). A phylogenetic tree of a subset of surveyed strains is shown on the left. EIC peak areas of the product aglycones are normalized to the maximum sum of product peak areas registered across all strains. Colors indicate different aglycone products as described in **Fig 3A**. Phylogenetically related, non-metabolizing strains were collapsed into a single node for clarity, indicated with a triangle. Strain composition in three synthetic human type strain communities is shown on the right, including HCom containing all 116 type strains and two dropout communities (DO1 and DO2) containing subsets of HCom strains. B) Type strains were combined to demonstrate cooperative metabolism of SGAs to highly modified aglycones in a bacterial community. Cultures individually grown in Mega medium were mixed and resuspended to high density in medium supplemented with 20 μM tomatine. Grey boxes indicate the strains included in each mixture. Type strains included two *Roseburia* sp. as tomatine deglycosylating strains that do not further modify the liberated aglycone and six Bacteroides sp. as tomatine-inactive strains that modify the liberated aglycone. EIC peak areas of tomatine, tomatidine, and modified aglycones were measured by LC-MS after incubation. Values shown are the mean±SD of replicate incubations (n=3), with individual replicates overlaid. Colors indicate different aglycone products. C) Schematic of type strain community engraftment and purified SGA dietary intervention (40 mmol/kg tomatine or solanine in diet) in gnotobiotic C57BL/6 mice. D) SGA and aglycone concentrations in plasma (nM) following dietary supplementation with tomatine or solanine (40 mmol/kg diet) in gnotobiotic C57BL/6J mice colonized with type strain communities. Values shown are the mean±SD for each group, with individual replicates overlaid (n=3-4). P values determined using a two-tailed t test. ns = not significant (P>0.05).

To directly assay for aglycone modification, we also cultured the commensal strains in media supplemented with the aglycones tomatidine, solanidine, and solasodine (**Fig S8**). Bacteroidetes were found to be broadly active on these compounds to generate the same sulfated and phosphoethanolamine-conjugated metabolites previously characterized in human stool. A few of the strains capable of SGA deglycosylation were also found to modify the resulting aglycone; *Bacteroides uniformis* ATCC 8492 produced sulfated and phosphoethanolamine-conjugated compounds following incubation with both tomatine and tomatidine. However, this behavior was not universal across strains, with different subsets of strains active on the aglycone and the corresponding glycoside.

We hypothesized that SGAs may be metabolized cooperatively in a community context, reliant on a subset of strains to liberate the aglycone before further modification by other members of the community. We explored this possibility by incubating cocultures of select strains with tomatine in rich medium (**Fig 4B**). Monocultures of *Roseburia intestinalis* LI-82 and *Roseburia inulinivorans* DSM 16841 produced unmodified tomatidine aglycone from an SGA input. However, culturing these strains in the presence of non-deglycosylating *Bacteroides* strains capable of aglycone modification enabled production of sulfonated and phosphoethanolamine-conjugated tomatidine. These results demonstrate that, despite their relative hydrophobicity, steroidal alkaloid aglycone intermediates can be exchanged between bacterial strains in a community. Furthermore, in this preliminary study with an eight member bacterial community, we show that the metabolic fate of SGAs can be manipulated by the inclusion or exclusion of certain strains from a microbial community *in vitro*.

To extend these results to more complex communities and explore their relevance *in vivo*, we generated three communities comprised of subsets of the type strains surveyed for SGA metabolism (**Fig 4A**). The first community, denoted HCom (Cheng et al., 2022; Wang et al., 2023), included all 116 type strains and represented a model Western microbiome capable of SGA metabolism. Community DO1 included 107 strains and was designed to drop out strains found to perform significant tomatine and solanine deglycosylation in monoculture. Community DO2 included 88 strains and was designed to additionally drop out strains that showed any detectable levels of solanine and tomatine deglycosylation, however slight, or were phylogenetically related to strongly metabolizing strains. Type strains were individually grown, pooled, and then inoculated into germ-free C57BL/6 female mice by oral gavage (**Fig 4C**). After two weeks of colonization, feces were collected and analyzed for relative strain abundance by metagenomic sequencing. We were able to detect successful engraftment of the majority of the strains from the pooled bacterial inoculum (**Fig S9A**) by their representation in feces (**Fig S9B**), with the exception of a limited subset of strains that reproducibly dropped below our signal to noise threshold. Importantly, the relative abundances of the strains excluded from DO1 and DO2 were significantly lower or undetectable in the dropout communities compared to HCom, indicating the successful dropout of metabolizing strains from colonized communities *in vivo* (**Fig S9B**). One notable exception was *Clostridiales bacterium* VE202-27 which appeared in similar abundance across all fecal samples, likely due to its incomplete exclusion from the dropout community pooled inoculum (**Fig S9A**).

Animals colonized with these three communities were provided either AIN93G diet, or diet supplemented with tomatine or solanine for two weeks. Relative strain abundances in feces collected after dietary intervention did not show significant differences in community composition as a result of SGA supplementation (**Fig S9C**). Following SGA supplementation, plasma and tissues were collected and analyzed by LC-MS for steroidal alkaloid content. To contextualize the metabolite output of the synthetic community, we started by identifying the SA product profile of tomatine (**Fig S10A**) and solanine (**Fig S10B**) in HCom-colonized mice and compared these with the SA products in conventional mice fed tomatoes (**Fig 2**, **Fig S5**). The SA products in these gnotobiotic animals recapitulated those characterized in conventional mice and human samples, showing that the synthetic HCom community can be used as a model SGA metabolizing community *in vivo*. Interestingly, HCom-colonized mice produced more of certain modified aglycones compared to conventional mice, including PEA-conjugated tomatidine and the ketone 3-oxotomatidine, highlighting the importance of community composition in determining host exposure to different SA products.

To further investigate the impact of community composition on host SA exposure, we quantified SA metabolites in HCom and dropout community colonized mice. The detected aglycones tomatidine and solanidine were only detected in the plasma of mice colonized with the complete HCom community and were absent from the plasma of mice colonized with either dropout community (**Fig 4D**). Furthermore, we observed significantly higher levels of unmetabolized SGA in plasma of mice colonized with dropout communities, indicating that gut community composition controls exposure to not only steroidal alkaloid aglycones but also glycosides. Profiling tissues that accumulate higher levels of SA metabolites than plasma, we noted that while SGA deglycosylation was around 10 to 100-fold lower in DO1 and DO2 colonized mice, there were still detectable quantities of aglycones in cecal contents, colon, kidney, and liver (**Fig S10C, Fig S10D**); we hypothesize that this residual deglycosylating activity in the dropout communities can be attributed to the incomplete exclusion of metabolizing strains in the inoculum (**Fig S9A**). Taken together, these results show that strain level metabolism of SGAs can be determined and programmed to direct the metabolic output of a designed community. Importantly, these communities approximate metabolism observed by human microbiota.

### SGA metabolism by gut microbiota dictates compound bioavailability and bioactivity

Having established that SGAs are major dietary inputs from Solanums and that gut microbiota dictate the metabolic fates of SGAs, we next explored the consequences of this metabolism on host physiology. As hydrolysis of the glycone by gut microbiota significantly impacts physiochemical properties like solubility and molecular size, we began by investigating the impact of SGA deglycosylation on compound bioavailability. To gauge the amount of compound absorbed into systemic circulation, we estimated the total concentration of absorbed SGA-derived compounds as the sum of SGAs and all aglycones in mouse plasma after a potato dietary intervention (**Fig 2A**, **Fig S11A**). Gut microbiome-mediated SGA deglycosylation significantly increased the concentration of SGA-derived metabolites in systemic circulation compared to the concentration in mice receiving antibiotics, suggesting that SA aglycones have increased bioavailability and absorption from the gut lumen relative to glycosides. For more direct interrogation of this hypothesis, we used a Caco-2 human colonic cell monolayer as a model for compound transport across the intestinal enterocyte barrier (Hubatsch et al., 2007). Potato SAs solanine, chaconine, and solanidine were added to the apical side of the monolayer representing the intestinal lumen. Following incubation, compound transport across the monolayer was determined using LC-MS. While the SGAs solanine and chaconine primarily remained on the apical side, the majority of the aglycone solanidine was detected on the basolateral side, indicating that solanidine is indeed better transported across the gut epithelium than SGAs (**Fig S11B**). These data show that metabolism of SGAs by the gut microbiota convert SGAs to a better absorbed form and dictate the circulating concentrations of steroidal alkaloid compounds in the host.

We next explored the impact of SGA metabolism on compound biological activity with a focus on potato SAs as these have been best investigated in prior work. SAs have been shown to have numerous effects on host physiology, including anti-inflammation (Kuo et al., 2017) and chemoprevention through cytotoxicity in cancer cell lines (Friedman et al., 2009; Kúdelová et al., 2013; Shieh et al., 2011). We began by exploring the impact of deglycosylation on SGA cytotoxicity *in vitro*. We treated the human colorectal adenocarcinoma line Caco-2 with potato SGAs solanine and chaconine and observed concentration-dependent loss of cell viability. In contrast, the aglycone solanidine had a negligible impact on cell viability (**Fig S11C**). A similar association between glycosylation and *in vitro* toxicity was observed for the tomato SGA tomatine and the aglycone tomatidine (**Fig 11D**) (Friedman et al., 2009). These data suggest that SGA deglycosylation by gut microbiota inactivates the cytotoxic potential of these compounds.

Uniquely among the dietary SGAs, the potato SGAs solanine and chaconine are also established cholinergic toxins in humans, capable of inhibiting the cholinesterases that terminate acetylcholine (ACh) neurotransmission by hydrolysis to inactive choline (Nigg et al., 1996; Roddick, 1989). This activity has led to SGAs being implicated in poisonings caused by the ingestion of improperly stored potatoes (McMillan and Thompson, J.C., 1979). Considering the established interactions between potato SGAs and these physiologically essential cholinesterases, we wanted to explore the effects of gut microbiota-mediated SGA metabolism on ACh signaling. Recombinant human acetylcholinesterase (AChE) and butyrylcholinesterase (BChE) were treated with solanine, chaconine, and solanidine. Enzyme activity in the presence of these compounds was monitored by tracking hydrolysis of substrate mimics acetylthiocholine and butyrylthiocholine, respectively, using Ellman’s assay (Ellman et al., 1961). The SGAs solanine and chaconine inhibited AChE activity relative to a vehicle treated control in a dose-dependent manner while the aglycone solanidine did not significantly impact AChE activity at concentrations up to 100 μM (**Fig 5A**). Though AChE activity was not impacted by solanidine at the concentrations tested, solanidine did inhibit BChE activity to similar extents to the SGAs (**Fig S11E**). These data show that deglycosylation of potato SGAs solanine and chaconine to the aglycone solanidine alleviates the AChE inhibitory but not the BChE inhibitory activity of these compounds.

**Figure 5.**
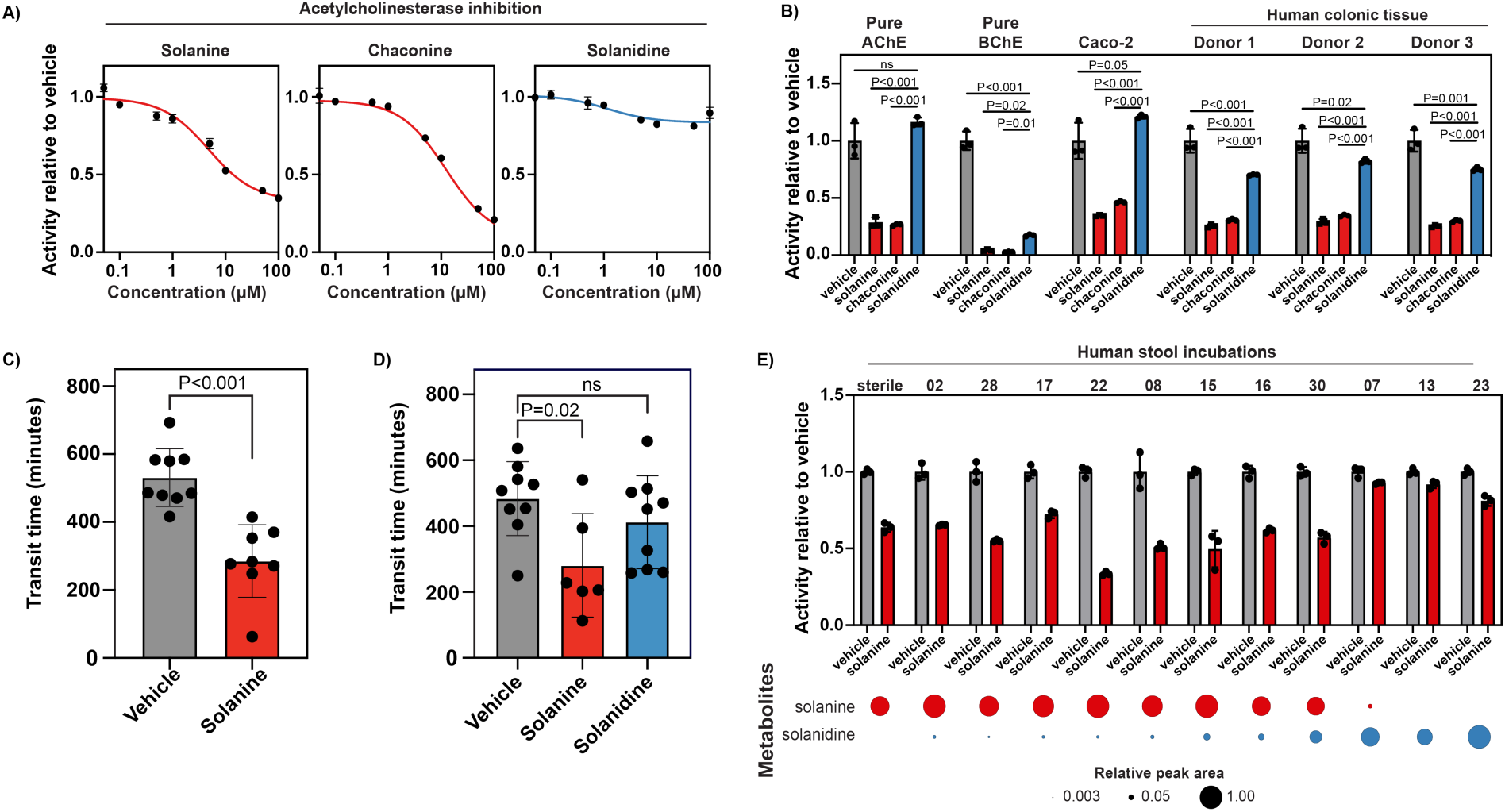
Solanine affects host acetylcholinesterase activity and whole gut transit in a glycosylation dependent manner. A) *In vitro* inhibition of recombinant human AChE activity by potato steroidal alkaloids. Data shown are normalized to AChE treated with vehicle only (DMSO for solanine and chaconine, ethanol for solanidine) with a total vehicle concentration of 1% for all treatments. Values shown are the mean±SD of three replicates. 5 ng AChE was used in each reaction. B) *In vitro* inhibition of cholinesterase activity by 50 μM potato SAs in human protein homogenates from normal colonic tissue (n=3 donors) and Caco-2 colonic epithelial cells. Activity was normalized to the activity of the vehicle-treated sample. Values shown are the mean±SD of three replicates. 5 ng each of recombinant human AChE and BChE were used for pure protein reactions, while 35 μg total protein was used for each tissue or cell homogenate. P values were determined using a two-tailed t test. ns = not significant (P>0.05). C) Whole gut transit time in male Swiss Webster mice receiving 50 mg/kg body weight solanine administered by oral gavage in a vehicle of 6% Carmine Red and 0.5% methylcellulose. Transit time is calculated as the time elapsed between oral gavage and the first appearance of a red fecal pellet. Values shown are the mean±SD for each group, with individual replicates overlaid (n=8 or 9). P values were determined using a two-tailed t test. ns = not significant (P>0.05). D) Whole gut transit time in male Swiss Webster mice receiving 50 mg/kg body weight solanine or an equivalent molar dose of 23 mg/kg body weight solanidine. Values shown are the mean±SD for each group, with individual replicates overlaid (n=6 or 9). P values were determined using Dunnett’s multiple comparison test. ns = not significant (P>0.05). E) Inhibition of recombinant human AChE activity by spent media from human stool incubated with 20 μM solanine in SAAC minimal medium. Activity was normalized to the activity of AChE treated with spent media from the same donor sample incubated with vehicle only. Values shown are the mean±SD of three replicates. The relative abundances of SGA and aglycone following incubation with solanine are shown below for each stool sample. Values shown are EIC peak areas normalized to the maximum peak area across all samples.

Beyond its role in the central nervous system, ACh is a key neurotransmitter in the enteric nervous system (Goyal and Hirano, 1996). Because AChE and BChE can both contribute to ACh hydrolysis in the gastrointestinal tract, we aimed to investigate the effect of SGA metabolites on a physiological mix of cholinesterases. We chose to focus on the colon as the primary tissue of interest, as SGAs are present in high concentrations in the gastrointestinal tract and are extensively metabolized by colonic microbiota. In mice receiving 5% potato peel in diet with antibiotic treatment to model a microbiome incapable of SGA metabolism, the sum of SGAs measured in colonic tissue was 2.98 ± 0.38 pmol/mg fresh weight tissue (**Fig S5E**), concentrations which corresponded to about 40% inhibition of AChE activity *in vitro* (**Fig 5A**). Protein homogenates were prepared from Caco-2 cells and human colonic tissue samples. We incubated these samples with equal concentrations of solanine, chaconine, and solanidine, and assayed for acetylthiocholine hydrolysis. SGAs strongly inhibited ACh hydrolysis in both Caco-2 and colon tissue homogenates (**Fig 5B**). In contrast, the aglycone solanidine only slightly inhibited ACh hydrolysis in tissue homogenates and did not inhibit activity at all in Caco-2 lysates. The difference between the slight effect of solanidine in colonic tissue and the lack of an effect in Caco-2 may be explained by a higher expression of solanidine-sensitive BChE in colonic tissue compared to Caco-2 (Plageman et al., 2002). Indeed, treatment of these samples with the BChE selective inhibitor iso-OMPA to isolate the contribution of BChE towards ACh hydrolysis indicated that about 40% of ACh hydrolysis can be attributed to BChE in colonic tissues compared to no detectable BChE activity in Caco-2 (**Fig S11F**). The modest inhibition of overall ACh hydrolysis by solanidine in colonic tissue was rescued by concurrent treatment with iso-OMPA, indicating that the mechanism for this activity is through BChE inhibition (**Fig S11G**). Taken together, these data show that potato SGAs inhibit overall ACh hydrolysis in relevant host tissues and deglycosylation of these compounds alleviates this inhibition.

The enteric nervous system controls motility, ion transport, and blood flow in the gastrointestinal tract and is mediated by neurotransmitters, including serotonin and ACh (Furness, 2012). Accordingly, a common pharmacological effect of AChE inhibitors is increased gastrointestinal motility, with drugs like neostigmine used to alleviate post-operative ileus (Kreis et al., 2001). We hypothesized that SGAs, as AChE inhibitory compounds that accumulate highly in the gastrointestinal tract, might similarly impact gut motility. To investigate the effects of SGAs on gut motility *in vivo*, Swiss Webster mice were administered the SGA solanine by oral gavage alongside Carmine Red as a non-absorbed tracer. A dose of 50 mg/kg body weight (bw) solanine was selected to be comparable to the daily SGA intake achieved through *ad libitum* consumption of diet supplemented with 5% potato peels (about 32 mg/kg bw total SGAs per day, **Fig 2A**). Whole gut transit was determined as the time required for red dye to appear in a fecal pellet following oral gavage, measured using an automated set up (Kacmaz et al., 2021). Male mice receiving solanine showed significantly decreased whole gut transit time relative to mice receiving vehicle, indicating increased gut motility (**Fig 5C**). Interestingly, female Swiss Webster mice did not exhibit a similar effect when treated with the same dose of solanine (**Fig S11H**). Though gut transit time was decreased by solanine treatment in male mice, fecal water content (**Fig S11I**) and cumulative fecal pellets (**Fig S11J**) did not significantly differ between solanine and vehicle treatments. These gut transit changes modulated by SGAs, combined with *ex vivo* data demonstrating the ability of SGAs to inhibit AChE, suggest that unprocessed dietary SGAs can modulate gut motility through an AChE mechanism.

In contrast to SGAs, SA aglycones did not inhibit AChE *in vitro* and *ex vivo*; to determine if these activities also alter *in vivo* function, we next aimed to measure the effect of deglycosylated SA metabolites on gut motility. Male Swiss Webster mice were administered vehicle or equimolar doses of solanine (50 mg/kg bw) or solanidine (23 mg/kg bw) by oral gavage alongside Carmine Red. While solanine treated mice showed decreased gut transit time, mice receiving the aglycone solanidine did not have gut transit times that were statistically significantly different from vehicle-treated mice (**Fig 5D**). These results indicate that, unlike the glycoside solanine, the aglycone solanidine does not have a significant impact on gut motility *in vivo*. Given that the aglycone is a direct product of gut microbiota metabolism, these results taken together suggest a critical role for the microbiota in mediating the effects of these dietary molecules on the host.

Having noted significant inter-individual differences in microbiome-mediated SGA conversion (**Fig 3**), we wanted to determine the effect of this variation on AChE inhibition. We used spent media from individual human stool samples incubated with solanine to modulate AChE activity. These donor samples were selected to show a range of solanine metabolism (**Fig 3A**), including samples inactive on solanine and samples that completely deglycosylated solanine. Solanine conversion by different donor samples resulted in different extents of AChE inhibition (**Fig 5E**). Media from metabolically inactive stool samples inhibited AChE while stool samples with extensive aglycone production showed only slight AChE inhibition. These data show that the inter-individual differences that exist in microbiota-mediated SGA metabolism can translate to varying levels of host AChE inhibition *ex vivo*. Taken together, our mapping of the metabolic fates of dietary SGAs revealed the crucial role of gut microbiota in controlling host exposure to and host interactions with these compounds.

## DISCUSSION

Despite the prevalence and abundance of specialized metabolites in diet, there is limited understanding of the metabolic fates of these molecules following consumption. This gap is exemplified by our work with dietary Solanums. Recent studies have noted the accumulation of steroidal alkaloid aglycones in human, mouse, and pig circulation following tomato consumption (Cichon et al., 2017; Do et al., 2024; Dzakovich et al., 2024; Hövelmann et al., 2020; Sholola et al., 2025). However, like most specialized metabolites in diet, the pathways through which these aglycones are formed from a glycosylated dietary input and *in vivo* consequence of this metabolism are largely unknown. Through antibiotics treatment in animal models, *ex vivo* incubations with human liver and stool, and metabolomics of human urine, we have established the mechanisms through which SGAs are metabolized *in vivo*. From these studies, we determined that the gut microbiome initiates metabolism and dictates exposure to Solanum molecules. Considering the impact of these SGAs on gut transit and neurotransmission, we have uncovered a layer of inter-individually varying metabolism that determines the effect of dietary compounds on host physiology. Taken together, our study provides a framework towards molecular resolution of the diet-derived components that impact human health.

To demonstrate the role of gut community composition in determining SGA metabolism *in vivo*, we rationally designed complex communities of ∼100 human commensal type strains to achieve different metabolic outputs from tomatine and solanine. This study builds on previous work establishing that similarly large and complex synthetic communities can be used to recapitulate and explore host-microbiome interactions (Cheng et al., 2022; Nagashima et al., 2023; Wang et al., 2023). These studies investigated HCom metabolism of host-derived molecules like amino acids (Cheng et al., 2022) and bile acids (Wang et al., 2023). There are only a few examples to date that use complex synthetic communities to transform defined dietary molecules in a predictable manner, including sugars *in vitro* (Clark et al., 2021) and N-acylethanolamines (Cheng et al., 2024), fiber (Kovatcheva-Datchary et al., 2019), and choline (Romano et al., 2015) *in vivo*. Our work uses HCom and its dropout communities to program the metabolism of specialized compounds that are abundant in diet. Importantly, HCom recapitulates the diversity of Solanum specialized metabolites observed in human samples. In our study, SGA metabolism was dramatically depleted by the exclusion of nine strains in community DO1, establishing a closely matched knock out community that will enable mechanistic interrogation of the interactions between diet, microbiota, and host physiology. Using our profiling of individual type strains, we envision that other dropout communities can be designed to understand the relative contributions of strains towards SGA metabolism, insights into the transfer of intermediates between strains, and formation of modified SA aglycones.

Beyond abundant steroidal glycoalkaloids from Solanums, other plant saponins have been shown to undergo gut microbiome-mediated deglycosylation. Ginsenosides from *Panax ginseng* were found to be deglycosylated to the protopanaxadiol aglycone, mediated by *Bacteroides*, *Bifidobacterium*, *Fusobacterium*, and *Prevotella* strains (Kim, 2018). In contrast to our observations with SGA metabolism, however, ginsenosides do not appear to be solely dependent on the gut microbiome for metabolism and are also hydrolyzed by host digestive processes (Han et al., 1982). Beyond saponins, other dietary plant glycosides are also known to be metabolized by the gut microbiota (Liou et al., 2020). Notable examples include flavonoids, known to undergo deglycosylation and extensive metabolism to form phloroglucinol, catechol, and equol, among other products (Schneider et al., 1999). The ability of broad specificity host glycosidases to metabolize flavonoids varies with the identity and linkage of the glycone moiety (Day et al., 1998). In contrast to many plant glycosides encountered in the diet and potentially resulting from the complexity of the glycone moieties, we find SGAs from Solanums to be unusually resistant to hydrolysis by host digestion. These results support previous observations that tomatine is stable to incubation in acidic conditions at 37°C (Friedman et al., 1998), and is reminiscent of nondigestible dietary fibers that are fermented by gut commensals (Zhao et al., 2018).

While Solanum pathogens (Kaup et al., 2005; Osbourn et al., 1995) clearly benefit from deactivation of these plant defense compounds (**Supplemental Discussion**), the motivation of gut commensals to deglycosylate SGAs is less apparent. One possible motive is access to the liberated sugars for competitive advantage in a community context; gut commensals are known to harvest sugars from plant glycosides for use as carbon sources (Rakoff-Nahoum et al., 2014). Regardless, identification of the strains involved in SGA deglycosylation enables further investigation into the ecological and genetic bases of this metabolism and understanding of its prevalence across human populations.

Though aglycones are formed from gut microbiota-mediated deglycosylation of dietary SGAs, we have found that they are transient intermediates both *in vivo* and in the context of a synthetic bacterial community. Our efforts to characterize the metabolic fates of SGAs have led to the identification of aglycones that are modified further by hydroxylation, sulfation, 3OH oxidation, Δ5,6 saturation, and phosphoethanolamine conjugation. There is rich biology associated with each of these transformations, including notable intersections with metabolism of steroids like cholesterol by the host and gut microbiota (**Supplemental Discussion**).

Crucially, metabolism of dietary SGAs by gut microbiota can transform the bioavailability and bioactivity of these compounds in the host. For example, the SA aglycones liberated by gut bacteria have been studied in a variety of physiological contexts (**Supplemental Discussion**). Notably, potato SGAs are known to act as AChE inhibitors (Roddick, 1989), a class of compound associated with drastic pharmacological and toxicological impacts (**Supplemental Discussion**). Systemic AChE inhibition, including irreversible enzyme inhibition by sarin and organophosphates, can be lethal due to the local accumulation of ACh and continued stimulation of vital nervous processes. However, reversible and tissue specific AChE inhibition is a clinical strategy for the treatment of Alzheimer’s disease (Ibach and Haen, 2004), glaucoma (Almasieh et al., 2010), and post-operative ileus (Andrew Luckey et al., 2003). We show in this study that the gut microbiota alleviates the AChE inhibitory properties of potato SGAs solanine and chaconine *in vitro* by deglycosylating the compounds to a non-inhibitory form (**Fig 5A**, **Fig 5D**). We further demonstrate that oral treatment with the potato SGA solanine, but not the corresponding aglycone solanidine, increases gastrointestinal motility *in vivo*. While the exact mechanism through which solanine affects gut transit *in vivo* remains to be investigated, we note that the phenotype correlates with molecule glycosylation similarly to AChE inhibition.

In the gastrointestinal tract where we found SGAs to significantly accumulate, acetylcholine is additionally a key regulator of multiple non-neuronal processes including chloride secretion (Yajima et al., 2011), epithelial cell proliferation (Yu et al., 2017), fluid secretion (Billipp et al., 2023), and incretin gut hormone release (Balks et al., 1997). Considering the role of ACh in these diverse physiological processes, we suspect that AChE inhibition by SGAs may impact intestinal physiology through a myriad of mechanisms beyond cholinergic toxicity. While there is still much to be investigated regarding the contributions of the gut microbiota to mitigating the symptoms of SGA exposure, we show that gut microbial metabolism can significantly deplete SGA levels in the colon *in vivo* following a solanine dietary intervention (**Fig S10D**). Furthermore, variation in the abilities of individual communities to deglycosylate potato SGAs (**Fig 4**) can lead to interindividual differences in host exposure to these bioactive SGAs. These results emphasize the importance of considering gut microbiota in future studies investigating the physiological impacts of SGAs.

Taken together, our study serves as a model for how the gut microbiota modulate host exposure and the biological effects of some of the most prevalent drug-like molecules from commonly consumed vegetables. Our data show how specialized dietary compounds can have measurable effects on host biology and how metabolism can be programmed at the level of the gut microbiome. This type of mechanistic analysis will likely contribute to our larger systems level understanding of how dietary interventions, coupled with gut microbiota metabolism, impact host health and prevent disease.

## ACKNOWLEDGEMENTS

This work was supported by USDA NIFA grant 2022-67017-36242 (to E.S.S.) and the ONO Pharma Foundation. P.C.K. was supported by NIH grant R01DK114007. We thank J.L. Sonnenburg (Stanford), T.W. Meyer, T.W. (Stanford), and the Stanford Tissue Bank for generously providing human samples for analysis. M. Carter (Stanford) for assistance accessing human stool samples; J.Z. Long and J.T. Kim (Stanford) for helpful discussions and preliminary experiments; A.G. Cheng (Stanford) and X. Feng (Stanford) for helpful discussions and sharing protocols; J. Arreola (Stanford) for assistance with strain growth; S. Jain (C.Z. Biohub) for assistance with phylogenetic analyses; D.L. Wengier (Stanford) for assistance with acquiring tomatoes; J.C.T. Liu (Stanford) for assistance with structural characterization and comments on this manuscript; J.E. Blum and S.P. Niehs for comments on this manuscript.

## CONTRIBUTIONS

C.S.L., T.L., P.C.K, and E.S.S designed experiments and analyzed data. C.S.L., T.L, M.I., J.R., P.P.M, S.K.H., A.W, X.M, A.V.C., E.A.S, M.J., P.H., and A.D. performed experiments. C.S.L. wrote the paper with edits from E.S.S.. M.A.F. assisted with microbiome communities.

## METHODS

### Chemicals

Steroidal alkaloid compounds were acquired from commercial sources: tomatine (Tokyo Chemical Industry, MedChemExpress), tomatidine (Cayman Chemical Company), solanine (Sigma Aldrich), chaconine (Sigma Aldrich), solanidine (Carbosynth), solamargine (MedChemExpress), solasonine (MedChemExpress), solasodine (Santa Cruz Biotechnology). All steroidal alkaloid compounds were prepared as 10 mM stocks in vehicle. Steroidal glycoalkaloids were dissolved in dimethylsulfoxide (DMSO). Tomatidine and solasodine were dissolved in methanol and solanidine was dissolved in ethanol. Other chemical compounds were acquired as follows: trimethylamine N-oxide (Cayman Chemical Company), trigonelline chloride (Cayman Chemical Company), hippuric acid (Sigma Aldrich), acetylthiocholine iodide (Cayman Chemical), butyrylthiocholine iodide (Tokyo Chemical Industry), and iso-OMPA (Sigma Aldrich).

### Mouse Studies

Animal studies were performed under protocols approved by the Stanford University Institutional Animal Care and Use Committee and Mayo Clinic Institutional Animal Care and Use Committee. For all studies, animals were allowed to acclimate for a week before any treatment.

#### *Ad libitum* fresh tomato feeding

Female Swiss Webster mice from 6-8 weeks of age (Taconic) were fed *ad libitum* with AIN93G diet (Dyets, Inc). Three days following the introduction of AIN93G diet, 40 g of red ripe Microtom tomatoes were provided daily alongside diet for each cage of five mice for 15 days, with uneaten tomatoes removed daily before replenishing with fresh tomatoes. The quantities of tomatine and esculeoside A in the provided tomatoes were determined by liquid chromatography-mass spectrometry (LC-MS) as described below to be about 720 μmol/kg fresh weight escueloside A and 74 μmol/kg fresh weight tomatine. Molar quantities of both compounds were estimated using a commercial tomatine standard. Mice were sacrificed by CO_2_ asphyxiation 24 hours following the final provision of tomatoes. Blood was collected by cardiac puncture into Microtainer SST tubes (BD) and serum harvested. All samples were frozen in liquid nitrogen and stored at −80°C until analysis by LC-MS.

#### Pure tomatine feeding

Female Swiss Webster mice from 6-8 weeks of age (Taconic) were fed *ad libitum* with AIN93G diet (Dyets, Inc). Three days following the introduction of AIN93G diet, mice were provided with 15 μM pure tomatine in drinking water. Drinking water was sterile filtered and refreshed every 2 to 4 days for 15 days, after which mice were sacrificed by CO_2_ asphyxiation and serum, cecal contents, colon, liver, kidneys, and mesenteric lymph nodes (MLNs) were harvested. Samples were frozen in liquid nitrogen and stored at −80°C until analysis by LC-MS.

#### Potato feeding

Female Swiss Webster mice from 6-8 weeks of age (Taconic) were fed *ad libitum* with AIN93G diet (Dyets, Inc). One week following the start of AIN93G diet, mice were dosed with 0.5 μmol solanine per mouse by oral gavage. Two days following this single dose, mice were provided with 10% w/w potato peels incorporated into AIN93G diet. Potato-supplemented diet was prepared by autoclaving peels from Yukon Gold potatoes at 121°C for 30 minutes, lyophilizing to dryness, and homogenizing using a blender to generate dry powder. Potato peel powder was mixed into powdered AIN93G diet (Dyets, Inc) at 10% w/w, moistened with water to pellet, and then lyophilized to dryness. The quantity of solanine and chaconine in the provided potato-supplemented diet was determined by LC-MS using commercial standards below to be about 400 μmol/kg diet chaconine and 110 μmol/kg diet solanine. Mice that did not receive potato diet instead received similarly pelleted AIN93G. After one day of potato-supplemented diet, mice were sacrificed by CO_2_ asphyxiation and serum was collected. Samples were frozen in liquid nitrogen and stored at −80°C until analysis by LC-MS.

#### Antibiotics treatment for *ex vivo* feces metabolism

Female Swiss Webster mice from 6-8 weeks of age (Taconic) were fed *ad libitum* with AIN93G diet (Dyets, Inc). Three days following the start of AIN93G diet, mice received five consecutive daily doses of 10 mg each ampicillin, vancomycin, metronidazole, and neomycin by oral gavage (Kuss et al., 2011). Each mouse received 0.25 mL of a combined stock of 40 g/L of each antibiotic prepared in phosphate buffered saline (PBS, pH 7.2). Mice not treated with antibiotics instead received 0.25 mL of PBS by oral gavage. Following five daily doses of antibiotics by oral gavage, antibiotics were provided *ad libitum* in drinking water for three days: ampicillin (1 g/L), vancomycin (0.5 g/L), neomycin (1 g/L), and metronidazole (1 g/L) (Croswell et al., 2009; Rakoff-Nahoum et al., 2004). Feces were collected into sterile tubes and processed as described below.

#### Concurrent antibiotics treatment and dietary intervention

Female C57BL/6J mice from 6-8 weeks of age (Jackson Laboratory) were fed *ad libitum* with AIN93G diet (Dyets, Inc). One week following the introduction of AIN93G diet, antibiotics were provided *ad libitum* in drinking water for the duration of the study: ampicillin (1 g/L), vancomycin (0.5 g/L), neomycin (1 g/L), and metronidazole (1 g/L). Antibiotics were refreshed every 3-4 days. Six days following the start of antibiotics treatment, mice were provided potato-supplemented diet (5% dry weight), prepared as described above. Diet supplemented with 5% (w/w) potato peels was analyzed using LC-MS and estimated to contain 132 μmol/kg diet chaconine and 53 μmol/kg diet solanine. Another set of mice was provided tomato-supplemented diet (10% dry weight) seventeen days following the start of antibiotics treatment. Sugar plum tomatoes were incorporated into AIN93G diet as described for potato supplemented diet above, but surface-sterilized with 70% ethanol and rinsed with water instead of autoclaved. Diet supplemented with 10% (w/w) tomato was analyzed using LC-MS and estimated to contain 42 μmol/kg diet tomatine and 16.5 μmol/kg diet esculeoside A using chemical standards for tomatine. Animals received potato supplemented diet for 8 days or tomato supplemented diet for 7 days, and were then sacrificed by CO_2_ asphyxiation. Serum, colon, livers, cecal contents, kidneys, and MLNs were collected and frozen in liquid nitrogen and stored at −80°C until analysis by LC-MS.

#### Gnotobiotic mouse study

Germ-free female C57BL/6 mice from 7-9 weeks of age were maintained in gnotobiotic isolators and provided gamma-irradiated AIN93G diet (Research Diets, Inc). Three days following the introduction of AIN93G diet, mice were inoculated by oral gavage with 200 μL of pooled bacterial glycerol stocks that were thawed to room temperature immediately prior. Inoculation was repeated the next day to ensure colonization of the pooled strains. Two weeks following inoculation, mice were continued on gamma-irradiated AIN93G or supplied with gamma-irradiated AIN93G supplemented with 40 mg/kg diet tomatine or solanine (Research Diets, Inc). Fresh fecal pellets were collected just prior to introduction of the dietary intervention and stored at −80°C until analysis by metagenomic sequencing. After two weeks of dietary intervention, animals were sacrificed by CO_2_ asphyxiation. Plasma, colons, livers, kidneys, and unpassed fecal pellets in colons were collected and stored at −80°C until analysis.

#### Whole gut transit assay

Whole gut transit was measured using an automated set up constructed as previously described (Kacmaz et al., 2021). Cylindrical plastic jars with lids were modified by replacing their bottoms with perforated aluminum sheets. These jars were placed on top of Teflon-coated funnels that led to 35 mm collection plates filled with 100% glycerol. A high-definition (HD) camera (TENVIS JPT3815 W-HD) was placed below each collection plate. The setup was surrounded by flexible LED strips to improve illumination when the cabinet doors were closed. VideoVelocity software was used to record time-lapse videos (1 frame every 10 s).

On the day of the experiment, male and female Swiss Webster mice from 6-8 weeks of age (Taconic) were fasted for one hour and then gavaged with 300 µL Carmine solution with solanine, solanidine, or no additional compound as a vehicle control. The solanine-containing Carmine red solution was prepared by dissolving solanine or solanidine in methylcellulose (0.5% w/v) (Sigma-Aldrich, USA) to 7.5 mM. Carmine red (6% w/v) (Sigma-Aldrich, USA) was added to the solanine-methylcellulose mixture. Mice were gavaged with 300 μL of 50 mg/kg bw of solanine-Carmine red, an equivalent molar dosage of 23 mg/kg bw of solanidine-Carmine red, or Carmine red vehicle. After gavage, mice were placed in the transit chambers and recordings were started (T_start_). The recordings were stopped after 12-13 hours, after which the time-lapse videos were analyzed and time-stamped for the occurrence of pellets. Transit time was calculated as the time elapsed between T_start_ and the appearance of the first red pellet (T_red_). For determination of fecal water content, fecal pellets were collected from each animal following the conclusion of the gut transit assay. Fecal pellets were weighed before and after drying. Fecal water content was calculated as 1-(dry mass/wet mass).

#### Sample preparation for LC-MS

Harvested tissues were lyophilized to dryness and then homogenized using a ball mill homogenizer (Retsch). Homogenized tissues were extracted with a fixed volume of 2:1:1 acetonitrile (MeCN):methanol(MeOH):water(H_2_O) per dry mass tissue, vortexed to mix, and then centrifuged at 18,000 *g* for 10 minutes. The resulting supernatant was diluted according to tissue type (for cecal content samples) and then filtered through a 0.45 μm PTFE filter for analysis by LC-MS.

Serum and plasma samples were thawed and proteins precipitated by the addition of three volumes of ice chilled 3:1 MeCN:MeOH. Samples were vortexed, stored at −80°C for at least two hours, and then centrifuged at 18,000 *g* for 10 minutes at 4°C. To concentrate the sample, supernatant was transferred to a new microcentrifuge tube, flash-frozen in liquid nitrogen, and lyophilized to dryness. Dried samples were resuspended in 2:1:1 MeCN:MeOH:H_2_O to the original volume and filtered through a 0.45 μm PTFE for analysis by LC-MS.

### Steroidal alkaloid metabolism by human liver enzymes

To assay for metabolism of steroidal alkaloids by human liver enzymes, 1 mg/mL of liver S9 proteins from pooled human donors (Sekisui XenoTech) was incubated in PBS pH 7.4, supplemented with 25 μM tomatine, tomatidine, solanine, chaconine, or solanidine. Samples were incubated at 37°C for 5 minutes. Metabolism was initiated by the addition of cofactors required for metabolic reactions and supplied as two separate mixtures: 1) 1 mM glutathione, 2 mM NADPH, and 0.5 mM acetyl-CoA, and 2) 200 μM 3’-phosphoadenosine-5’-phosphosulfate (PAPS), 500 μM uridine diphosphate glucuronic acid (UDPGA), 25 μg/L alamethicin, 2 mM magnesium chloride, and 2 mM NADPH. Each reaction was prepared in duplicate, with one set immediately quenched by the addition of three volumes of cold 3:1 MeCN:MeOH for a pre-incubation control. The other set of reactions was incubated at 37°C for 60-90 minutes and then quenched as before. Quenched samples were stored at −80°C overnight, thawed, and centrifuged at 18,000 *g* for 10 minutes. The resulting supernatant was collected and filtered through a 0.45 μm PTFE filter for analysis by LC-MS.

### Synthesis of standards for steroidal alkaloid metabolites

#### 3-oxotomatidine

Synthesis of 3-oxotomatidine was adapted from previous reports (Chagnon et al., 2014). Tomatidine hydrochloride was dissolved in dichloromethane (DCM). Three molar equivalents of Dess-Martin periodinane (DMP, Alfa Aesar) were added and the mixture was stirred at room temperature overnight. The reaction was quenched with one volume of 0.8 M sodium thiosulfate and saturated sodium bicarbonate, stirred at room temperature for 1 hour, and then extracted three times with one volume of DCM. The organic layers were combined and dried on anhydrous calcium sulfate. LC-MS analysis of the reaction products at this stage yielded an estimated tomatidine conversion of about 50%. To improve substrate conversion, three equivalents of DMP were again added to this mixture and the workup repeated as described above. LC-MS analysis of the crude reaction after a second round of reaction generally showed close to full conversion, with little detectable remaining unreacted tomatidine.

To isolate the 3-oxotomatidine product, the crude reaction mixture in DCM was dried down, resuspended in a small volume of N,N-dimethylformamide (DMF), and separated by reversed phase chromatography using a 100Å, 30 μm, 12 g Sfär C18 D Duo column (Biotage). Water with 0.1% (v/v) formic acid and acetonitrile with 0.1% (v/v) formic acid were used as mobile phase solvents with a flow rate of 12 mL/min. Chromatographic separation was achieved using the following linear gradient (with percentages indicating levels of acetonitrile with formic acid): 10%, 3 column volumes (CV); 10% to 20%, 3 CV; 20% to 50%, 25 CV; 50% to 95%, 5 CV; 95%, 10 CV. Fractions were assayed by LC-MS; fractions containing pure 3-oxotomatidine were combined and evaporated to dryness.

#### Sulfotomatidine

Sulfotomatidine synthesis was adapted from previous reports for the sulfation of dietary phenolics (Almeida et al., 2017). Tomatidine hydrochloride was dissolved in pyridine and combined with one molar equivalent of sulfur trioxide pyridine complex. The mixture was sparged with argon and stirred at 65°C for 24 hours. The reaction was quenched with one volume of water and the organic solvent evaporated. The crude reaction mixture was analyzed by LC-MS for retention time match and MS/MS match to the mass feature in biological samples.

#### Dihydrosolanidine

Dihydrosolanidine was synthesized from solanidine (Carbosynth) by hydrogenation mediated by a palladium on carbon catalyst. Solanidine was dissolved in ethanol and one mass equivalent of palladium on carbon was added with stirring. Hydrogen gas was bubbled through the mixture for 6 hours and the reaction was filtered to remove catalyst. The crude reaction mixture was analyzed by LC-MS for retention time match and MS/MS match to the sample mass feature.

### Human urine from colectomy patients

De-identified urine samples were obtained from a previously published study (Mair et al., 2018). The donors were 12 patients with total colectomies (6 male and 6 female) and 17 patients with intact colons with no known gastrointestinal disease and no antibiotic use for one month pre-collection (11 male and 6 female). Donors did not receive specified dietary regimens or supplements.

Urine samples were centrifuged at 18,000 *g* for 10 minutes and then analyzed by LC-MS as described below. Sample creatinine levels were measured using a colorimetric creatinine assay kit (Abcam) and normalization factors were calculated using the Cockcroft-Gault equation (Cockcroft and Gault, 1976). Ion counts for metabolites of interest were multiplied by the normalization factor to enable comparison in compound abundance between samples.

### Bacterial and fecal culturing

Fecal samples and human commensal strains were handled under anaerobic conditions using the GasPak anaerobic system (BD) or an anaerobic chamber (Coy) in an atmosphere of 10% CO_2_, 5% H_2_, and 85% N_2_. To maintain a reducing environment for fecal bacteria during sample collection and handling, fecal samples were collected and resuspended in pre-reduced transport medium (Cheng et al., 2022). Transport medium contained, per L: 0.2 g KCl, 0.1 g CaCl2, 0.1 g MgCl2, 0.2 g KH_2_PO_4_, 1.15 g Na_2_HPO_4_, 3 g NaCl, 0.1 g uric acid, 0.1 g glutathione. For growth on solid media where applicable, bacterial strains were grown on brain heart infusion agar supplemented with 5 μg/mL hemin and 0.5 μg/mL menadione (BHIS). For growth in liquid cultures, bacterial strains were grown in the following media: Mega medium (Wu et al., 2015) modified to exclude soluble starch and Tween 80, modified Columbia medium, yeast casitone fatty acid medium (YCFAM), yeast casitone fatty acids broth with carbohydrates (YCFAC, Anaerobe Systems), modified peptone yeast glucose medium (PYG, DSMZ Medium 104), brain heart infusion (BHI), modified Gifu anaerobic broth (mGAM, Hyserve), Wilkins-Chalgren broth (DSMZ Medium 339a), Ethanoligenes medium (DSMZ Medium 1057), tryptic soy broth (TSB) with 5% blood, Bifidobacterium medium, and chopped meat medium. More details on strain culturing can be found in their respective experimental sections below and in **Supplemental Table 2**.

### *Ex vivo* steroidal alkaloid metabolism by mouse feces

To assay for steroidal glycoalkaloid metabolism by murine fecal bacteria, freshly collected fecal pellets were suspended at 2% w/v in pre-reduced transport medium and then vortexed until homogenized. Larger particulates were allowed to settle, and the slurry was transferred to a new tube. An equal volume of pre-reduced transport medium containing 40 μM tomatine was added to the 2% w/v fecal slurry, resulting in a final incubation of 1% w/v feces with 20 μM tomatine. Samples were incubated anaerobically using the GasPak anaerobic system for 36 hours at 37°C. Following incubation, two volumes of 3:1 MeCN:MeOH were added to extract metabolites and quench metabolism. Samples were vortexed, centrifuged at 12,000 *g* for 10 minutes, and then supernatant was diluted and filtered through a 0.45 μm PTFE filter for analysis by LC-MS.

### *Ex vivo* human stool metabolism

De-identified stool samples from human donors were obtained from the RAMP study (Rejuvenating the Aging Microbiota with Prebiotics, clinical trial ID: NCT03690999) (Carter et al., 2025). This cohort was comprised of 89 individuals (49 female and 40 male) ranging from 60 years to 83.9 years. A subset of feces samples from thirty donors were randomly selected and assayed for SGA metabolism. Samples were stored at -20°C in donor’s home freezers until they were transferred to the research laboratory and stored at −80°C until use. Frozen samples were transferred into an anaerobic chamber, suspended at 10% w/v in pre-reduced transport medium with 20% glycerol, vortexed for 2 minutes to homogenize, and then allowed to settle. The stool slurry was aliquoted into cryotubes, flash-frozen in liquid nitrogen, and stored at −80°C until use. For use in metabolism assays, frozen stool resuspensions were thawed and diluted ten-fold to 1% w/v in pre-reduced transport medium. The 1% w/v stool slurry was combined in 96 well plate format with an equal volume of 2x standard amino acid complete minimal medium (SAAC), prepared as described by Dodd, *et al* (Dodd et al., 2017), supplemented with 40 μM of steroidal alkaloid substrate. Each reaction contained final concentrations of 0.5% w/v stool sample, 1x SAAC, 0.5x transport medium, and 20 μM steroidal alkaloid substrate. Each combination of stool sample and substrate was prepared in two sets of triplicates; one set was incubated anaerobically at 37°C for 48 hours while another set was immediately frozen in liquid nitrogen as an unmetabolized control and stored at −80°C. Following incubation for 48 hours, the other set of reactions was also frozen in liquid nitrogen and stored at −80°C.

To mitigate effects from differential evaporation between wells, samples were lyophilized to dryness in 96 well plates and resuspended in three volumes of 2:1:1 MeCN:MeOH:H_2_O. Plates were vortexed and sonicated for 5 minutes to resuspend and then stored at −80°C overnight to precipitate any proteins. Samples were centrifuged at 3,000 *g* for 10 minutes, diluted in 2:1:1 MeCN:MeOH:H_2_O, and then filtered through a 0.45 μm PTFE filter for analysis by LC-MS.

### Human commensal type strain culturing and metabolism

Media was prepared for each strain (**Supplemental Table 2**) and pre-reduced overnight in an anaerobic chamber. Bacterial glycerol stocks stored at −80°C were thawed at room temperature for 15 minutes and then inoculated at 100x dilution into their respective media in a 96 well format. Cultures were grown anaerobically at 37°C for three days. Individual cultures were pelleted by centrifugation at 4700 *g* for 5 minutes, and then resuspended in pre-reduced PBS. Resuspended cultures were used to inoculate modified Mega medium supplemented with 25 μM steroidal alkaloid substrate. Mega medium (Wu et al., 2015) was modified to exclude soluble starch and Tween 80 to maintain compatibility with downstream LC-MS analysis. Cultures were grown anaerobically at 37°C for 2 days. After incubation, cultures removed from the anaerobic chamber, flash-frozen in liquid nitrogen, and lyophilized to dryness to mitigate well-to-well variation in evaporation. Dried reactions were resuspended in three times the original reaction volume with 2:1:1 MeCN:MeOH:H_2_O, vortexed and sonicated to resuspend, and then frozen at - 80°C. Prior to analysis by LC-MS, samples were centrifuged at 4000 *g* for 15 minutes to remove particulates, diluted with 2:1:1 MeCN:MeOH: H_2_O, and then filtered through a 0.45 μm PTFE filter.

### Phylogenetic analysis of human commensal type strains

Taxonomic classifications of the human commensal type strains were determined using the GTDBtk v.1.6.0 classify workflow which uses 120 marker genes from bacterial whole genome sequences to place strains of interest in a GTDBtk reference tree (release202) (Chaumeil et al., 2020). Bacterial genome sequences were obtained from the NIH Human Microbiome Project. Trees were visualized using Interactive Tree of Life (iTOL) version 7.2.2 (Letunic and Bork, 2024).

### Preparation of synthetic communities

#### Small communities for *in vitro* metabolism

*Bacteroides* and *Roseburia* strains were streaked from glycerol stocks onto pre-reduced BHIS plates and incubated anaerobically in an anaerobic chamber at 37°C for one day. Single colonies were inoculated into pre-reduced modified Mega medium and grown anaerobically at 37°C for about one day. Liquid cultures were subcultured once more into modified Mega medium and allowed to grow anaerobically for a day. An equal volume of each culture was combined, pelleted by centrifugation at 6,000 *g* for 10 minutes, and resuspended in one volume of fresh modified Mega medium supplemented with 20 μM tomatine substrate. Cocultures were incubated anaerobically at 37°C for three days. Samples were frozen at −80°C and lyophilized to dryness. Dried samples were then extracted with three volumes of 2:1:1 MeCN:MeOH:H_2_O, vortexed and sonicated to resuspend, and centrifuged at 4000 *g* for 15 minutes and filtered through a 0.45 μm PTFE filter prior to LC-MS analysis.

#### Complex communities for *in vivo* experiments

Media was prepared for each strain (**Supplemental Table 2**) and pre-reduced in aliquots of 40 mL per strain in 50 mL conical tubes. Bacterial glycerol stocks stored at −80°C were thawed at room temperature for 15 minutes and then inoculated with 100x dilution into their respective culture tube. Cultures were grown anaerobically at 37°C for three days. Strains were pooled according to desired community composition and then pelleted by centrifugation at 4700 *g*. Bacterial pellets were resuspended in modified Columbia medium supplemented with 25% glycerol to a total volume of 40 mL. An aliquot was removed and stored at −80°C for metagenomic sequencing of bacterial inoculum. The reminder was aliquoted into 1.2 mL volumes in cryovials and stored at −80°C until used to inoculate germ-free mice.

### Metagenomic sequencing of bacterial communities

Genomic DNA from pooled bacterial inoculum and mouse fecal pellets was extracted using the DNeasy 96 PowerSoil Pro kit (Qiagen) and quantified in 96-well format using the Quant-iT PicoGreen dsDNA Assay kit (Thermo Fisher). A minimum of 15 ng of DNA was taken forward to construct metagenomics sequencing libraries using the Illumina DNA Prep kit, with half-volumes being utilized at each step to minimize cost. Five cycles of PCR were utilized to amplify DNA and introduce unique dual indices. Post PCR, libraries were purified using a 0.8x bead clean and were quantified using the the Quant-iT PicoGreen kit. Equal masses of each metagenomics library were pooled, and a dual-sided AMPure XP (Beckman Coulter) bead clean was performed on the pooled material to achieve buffer removal and proper size-selection for sequencer loading. The final library pool was quality-checked for size distribution using the Tapestation 2200 (Agilent Technologies). Sequencing was performed on the NovaSeq6000 (Illumina) using a 2x150bp read configuration, with 20 million paired-end reads being targeted per sample. Raw reads were processed using NinjaMap (an algorithm introduced by (Cheng et al., 2022)).

Relative strain abundances were calculated by normalizing the raw NinjaMap output of each strain by the sum of strains in a given sample. Relative abundances less than 10^-7^ were considered undetectable (Cheng et al., 2022) and set to a minimum level of 10^-7^ before log10 transformation. To identify any strains enriched or depleted by dietary intervention, unpaired two-tailed T tests were performed between fecal pellets from collected from individuals engrafted with a given community before dietary intervention and two weeks after dietary intervention. False discovery rate (FDR) corrected q values were determined using the R package ‘qvalue’ version 2.42 (Storey et al., 2004).Thresholds for significant differences in relative strain abundances were set to q<0.05 and fold change > 10.

### *In vitro* cell viability assays

The human intestinal epithelial cell line Caco-2 was obtained from the lab of Sarah Heilshorn (Stanford) and used between passages 5 to 15. Cells were grown and maintained in DMEM (ATCC) supplemented with 10% fetal bovine serum, 1% nonessential amino acids, 100 U/mL penicillin, and 100 μg/mL streptomycin at 37°C in 5% CO2. Cells were seeded into 96 well plates at a density of 10,000 cells per well and incubated for 17 hours for cells to adhere. Cells were then treated with varying concentrations of steroidal alkaloid compounds dissolved in solvent vehicle. The total amount of vehicle added to each well was normalized to 0.4% v/v. As a nontreated control, cells were treated with vehicle alone. Cells were incubated with compound at 37°C for 24 hours, after which cell viability was determined using the CellTiter-Glo Luminescent Cell Viability Assay (Promega). All treatments were performed in triplicate and repeated in three separate experiments.

### Caco-2 permeability assay

Transport of steroidal alkaloid compounds *in vitro* were determined using differentiated Caco-2 cells as a model monolayer (Hubatsch et al., 2007). Cells were seeded into 12 well plates fitted with 0.4 μm polycarbonate Transwell inserts (Corning) at a density of 300,000 cells per insert. The apical compartment above the insert was filled with 0.5 mL of medium and the basolateral compartment below the insert was filled with 1.5 mL of medium. Medium was refreshed on both the apical and basolateral sides every 2 to 3 days for 21 days for monolayer formation. After 21 days, cells were rinsed with serum free DMEM supplemented with 1% nonessential amino acids, 100 U/mL penicillin, and 100 μg/mL streptomycin. The apical compartment was filled with serum-free DMEM additionally supplemented with nontoxic concentrations of steroidal alkaloid compounds, as determined by viability assays described above (solanine: 10 μM, chaconine: 3 μM, solanidine: 10 μM). A subset of wells was treated with vehicle only as a control. The basolateral compartment was filled with serum-free DMEM supplemented with 4% bovine serum albumin (BSA) to mimic physiological protein concentrations on either side of the intestinal epithelium (Hubatsch et al., 2007). After 17 hours of incubation with steroidal alkaloid compounds, medium was collected from both the apical and basolateral components. An aliquot of medium was assayed for extracellular LDH using the LDH-Glo Cytotoxicity Assay (Promega) to ensure that cell monolayers were not compromised by compound treatment. All compound-treated wells showed similar LDH levels to vehicle only treatment, indicating that compound treatments at the selected concentrations did not impact cell viability. The remainder of the medium collected was assayed for steroidal alkaloid content using LC-MS. To prepare medium for analysis, samples were diluted with nine volumes of 3:1 MeCN:MeOH, kept at -20°C overnight to precipitate proteins, centrifuged at 10,000 *g* for 10 minutes, and filtered through a 0.45 μm PTFE filter.

### Cholinesterase activity assay

Acetylcholinesterase (AChE) and butyrylcholinesterase (BChE) activities were measured using Ellman’s method, adapted to a 96-well format (Ellman et al., 1961). For acetylcholinesterase activity, 5 ng of recombinant human AChE (Sigma) or 30 μg of tissue lysate was incubated with steroidal alkaloid compound or at room temperature for 30 minutes in 50 mM Tris pH 8.0 in a total volume of 90 μL. To initiate reactions, 5 μL of 10 mM acetylthiocholine iodide and 5 μL of 10 mM 5,5’-dithiobis-(2-nitrobenzoic acid) (DTNB) were added to each well and absorbance at 405 nm was measured over the course of 30 minutes. BChE activity was analogously measured using 5 ng of recombinant human BChE (R&D Systems) and butyrylthiocholine iodide. Enzyme activity was quantified as the linear slope of absorbance over time. Relative inhibition by steroidal alkaloid compounds was quantified by normalization to enzyme activity in the presence of vehicle.

### Colon lysate preparation

Normal human colon tissue from three de-identified human donors was obtained from the Stanford Tissue Bank. About 100 mg of tissue was homogenized in 2 mL of tissue lysis buffer (50 mM Tris pH 8, 150 mM sodium chloride, 0.5% Triton X-100, 10 μg/mL leupeptin, 10 μg/mL aprotinin, 20 μM pepstatin, and 2.5 mM benzamidine hydrochloride) using a gentleMACs Tissue Dissociator (Miltenyi). Homogenized samples were centrifuged at 2,000 *g* for 5 minutes at 4°C to collect lysate, transferred to new centrifuge tubes, and centrifuged at 9,000 *g* for 20 minutes at 4°C to pellet cell debris.

To prepare protein lysates from Caco-2 intestinal epithelial cells, cells were maintained as described above. A confluent 10 cm plate was rinsed three times with cold PBS, lysed by adding tissue lysis buffer as described above, and incubated with rocking at 4°C for 20 minutes. Lysate was collected and centrifuged at 9,000 *g* for 20 minutes at 4°C to remove cell debris.

To the resulting supernatants from human tissue and Caco-2 cells, glycerol was added to reach a final concentration of 20% glycerol. Protein concentrations in lysates were estimated using BSA standards and a detergent-compatible Bradford assay (Abcam). Lysates were aliquoted, flash-frozen in liquid nitrogen, and stored at −80°C until used in Ellman’s assay as described above. To assay the contributions of AChE to acetylthiocholine hydrolysis in these homogenates, BChE-specific inhibitor iso-OMPA was added to homogenates at a concentration of 500 μM and incubated at room temperature for 30 minutes before the assay was initiated with acetylthiocholine and DTNB.

### Acetylcholinesterase inhibition by human stool samples

A subset of eleven of the previously assayed de-identified human stool samples was selected for their differences in solanine metabolism, including samples that did not appear to metabolize solanine (02, 17, 22, 28), samples that deglycosylated solanine (07, 13, 23, 30), and samples that appeared to partially deglycosylate solanine by removing a glucose moiety from the glycone (08, 15, 16). *Ex vivo* incubations with 20 μM solanine or DMSO vehicle were prepared as described above for each of the donor samples. For compatibility with Ellman’s assay, we minimized the presence of free thiols in the assay medium; these *ex vivo* stool incubations were performed in pre-reduced SAAC minimal medium formulated without cysteine. Samples were incubated anaerobically at 37°C for 48 hours, after which an aliquot was removed for LC-MS analysis. This aliquot was diluted in 3:1 MeCN:MeOH to precipitate proteins, centrifuged at 12,000 *g* for 10 minutes, diluted in 2:1:1 MeCN:MeOH:H_2_O, and then filtered through a 0.45 μm PTFE filter prior to analysis by LC-MS. The remainder of the samples was spun down at 12,000 *g* for 20 minutes and the resulting supernatant used to inhibit recombinant human AChE as described above, with a few adaptations. These AChE assays were performed in spent supernatant from human stool cultures (spent SAAC without cysteine), instead of 50 mM Tris pH 8. Incubation with stool samples was found to lower pH of the media; to adjust the assay pH to levels amenable to AChE activity and DTNB-thiol conjugation, equal volumes of sodium hydroxide was added to samples to reach a final concentration of 6 mM. Sodium hydroxide was not added to media-only controls (i.e. sterile).

### LC-MS

#### *Ad libitum* fresh tomato feeding and pure tomatine feeding

Mouse serum and tissues resulting from fresh tomato supplementation and pure tomatine feeding were analyzed on an Agilent 1260 Infinity II HPLC using both reversed-phase liquid chromatography (LC) and normal phase LC. Reversed-phase LC was conducted using a Gemini NX-C18 2x100 mm, 5 μm column (Phenomenex) at 40°C. Mobile phase solvents were water with 0.1% (v/v) formic acid and acetonitrile with 0.1% (v/v) formic acid at a flow rate of 0.4 mL/min. A volume of 3 μL of sample was injected. Chromatographic separation was achieved using the following linear gradient (with percentages indicating the levels of acetonitrile with formic acid): 3% to 97%, 20 minutes; 97%, 5 minutes; 97% to 3%, 1 minute; 3%, 5 minutes. Normal phase liquid chromatography was conducted using a Poroshell 120 HILIC-Z, 2.1x100 mm, 2.7 μm column (Agilent) at 40°C. Mobile phase solvents were 10 mM ammonium formate supplemented with 0.1% formic acid and 90% acetonitrile with 10 mM ammonium formate supplemented with 0.1% formic acid at a flow rate of 0.25 mL/ min. A volume of 2 μL of sample was injected. Chromatographic separation was achieved using the following linear gradient (with percentages indicating the levels of acetonitrile-containing mobile phase): 100% to 60%, 15 minutes; 60%, 2 minutes; 60% to 100%, 1 minute; 100%, 8 minutes. For both types of chromatography, coupled mass spectrometry (MS) data was collected with an Agilent 6545 quadrupole time-of-flight (qTOF) ESI mass spectrometer in positive polarity. Parameters for the 6545 qTOF MS were as follows: mass range: 50-1700 m/z, gas temperature: 250°C, nebulizer: 10 psi, drying gas flow rate: 12 L/min, sheath gas temperature: 300°C, sheath gas flow rate: 12 L/min, fragmentor: 100 V, skimmer: 50 V.

#### Hydrogen-deuterium exchange

Human stool incubated with tomatine was chromatographically separated on an Agilent 1260 Infinity II HPLC using reversed phase (LC) using reversed-phase LC. LC was conducted using a Gemini NX-C18 2.0x100 mm, 5 μm column (Phenomenex) at room temperature. Mobile phase solvents were deuterium oxide (D_2_O) with 0.1% (v/v) deuterated formic acid (FA-D_2_) and acetonitrile with 0.1% (v/v) deuterated formic acid (FA-D_2_) at a flow rate of 0.4 mL/min. A volume of 5 μL of sample was injected. Chromatographic separation was achieved using the following linear gradient (with percentages indicating the levels of acetonitrile with FA-D_2_: 3% to 50%, 20 minutes; 50% to 97%, 1 minute; 97%, 5 minutes; 97% to 3%, 1 minute; 3%, 5 minutes. Coupled mass spectrometry (MS) data was collected with an Agilent 6520 qTOF mass spectrometer in positive polarity. Parameters for the 6520 qTOF MS were as follows: mass range: 50-1700 m/z, gas temperature: 300°C, nebulizer: 35 psi, drying gas flow rate: 10 L/min, fragmentor: 150 V, skimmer: 65 V.

#### Human urine

Urine from human colectomy patients and donors with intact colons were chromatographically separated on an Agilent 1260 Infinity II HPLC using both reversed-phase LC. LC was conducted using a SB-C18 3.0x100 mm, 1.8 μm column (Agilent) at 40°C. Mobile phase solvents were water with 0.1% (v/v) formic acid and acetonitrile with 0.1% (v/v) formic acid at a flow rate of 0.4 mL/min. A volume of 5 μL of undiluted sample was injected. Chromatographic separation was achieved using the following linear gradient (with percentages indicating the levels of acetonitrile with formic acid): 3% to 97%, 20 minutes; 97%, 4 minutes; 97% to 3%, 1 minute; 3%, 4 minutes. Coupled mass spectrometry (MS) data was collected with an Agilent 6545 qTOF ESI mass spectrometer in positive polarity. Parameters for the 6545 qTOF MS were as follows: mass range: 50-1700 m/z, gas temperature: 250°C, nebulizer: 60 psi, drying gas flow rate: 6 L/min, sheath gas temperature: 250°C, sheath gas flow rate: 11 L/min, fragmentor: 140 V, skimmer: 65 V.

#### All other studies

Samples from all other studies were separated on an Agilent 1290 Infinity II HPLC using reversed phase (LC). Chromatography for mouse serum and tissue samples and human liver enzyme incubations was performed using a Zorbax RRHT Eclipse Plus C18, 2.1x100 mm, 1.8 μm column (Agilent) at 40°C. Mobile phase solvents were water with 0.1% (v/v) formic acid and acetonitrile with 0.1% (v/v) formic acid at a flow rate of 0.4 mL/min. Chromatographic separation was achieved using the following linear gradient (with percentages indicating the levels of acetonitrile with formic acid): 3% to 97%, 20 minutes; 97%, 5 minutes; 97% to 3%, 1 minute; 3%, 5 minutes. Chromatography for human stool incubations, commensal type strain cultures, bacterial cocultures, and Caco-2 monolayer transport studies was performed using a Zorbax RRHT Eclipse Plus C18, 2.1x50 mm, 1.8 μm column (Agilent) at 40°C. Mobile phase solvents were water with 0.1% (v/v) formic acid and acetonitrile with 0.1% (v/v) formic acid at a flow rate of 0.6 mL/min. Chromatographic separation was achieved using the following linear gradient (with percentages indicating the levels of acetonitrile with formic acid): 3% to 60%: 6 minutes, 60% to 95%: 0.1 minutes, 95%: 1.5 minutes, 95% to 3%: 0.1 minutes, 3%: 1 minute. For all samples, a volume of 1 μL of sample was injected. MS data was collected using an Agilent 6546 qTOF ESI mass spectrometer in positive polarity. Parameters for the 6546 qTOF MS were as follows: mass range: 50-1700 m/z, gas temperature: 325°C, nebulizer: 35 psi, drying gas flow rate: 10 L/min, sheath gas temperature: 400°C, sheath gas flow rate: 12 L/min, fragmentor: 140 V, skimmer: 45 V.

#### Data analysis: untargeted metabolomics

Untargeted metabolomics was performed using XCMS Online (Tautenhahn et al., 2012).

Mass features were detected using the centWave algorithm, with the following parameters:

Maximal tolerated m/z deviation: 30 ppm
Chromatographic peak width [min, max]: [10 seconds, 60 seconds] Signal to noise threshold: 10
Minimum m/z difference in features with overlapping retention times: 0.01

Retention time correction was performed using the obiwarp method, with a step size of m/z 0.5. Peak alignment was performed with bandwidth of 3 seconds and minimum fraction (minfrac) of samples necessary for a valid group of 0.5.

From the table of mass features generated by XCMS Online, filtering criteria were applied to identify features present or differentially accumulating in certain treatment groups. The first filtering criteria was retention time; all mass features that eluted before 1 minute were assumed to not be retained on the column and therefore discarded. Feature peak areas that were statistically significantly different between treatment groups were then identified by two-tailed Student’s t-test (p < 0.1 for normal phase analysis and p < 0.05 for reverse phase analysis). Features that were higher in the treatment group of interest (e.g. higher in the serum of tomato-fed mice than in the serum of unsupplemented mice) were identified by fold change difference (ratio of average peak areas between groups). Mass features above a threshold signal (peak height > 10^4^) were then retained. Finally, putative parent ions were identified by retention time and peak area clustering to remove adducts, potential fragment ions, dimers, and isotopologues.

#### Data analysis: semi-targeted analysis for steroidal alkaloids in complex samples

To search for compounds in human stool incubations containing steroidal alkaloid substructures, one set of replicates was analyzed with a constant collision energy of 40V (for tomatidine-derived compounds) or 60V (for solanidine- and solasodine-derived compounds). Ions characteristic of tomatidine, solanidine, and solasodine fragmentation at these collision energies were extracted; tomatidine-derived: [M+H]^+^ 161.1325, solanidine-derived: [M+H]^+^ 98.0964, and solasodine-derived: [M+H]^+^ 157.1012. To find the intact mass features that yielded these characteristic fragments, the same sample was run with a collision energy of 0V to search for mass features eluting at the same retention time after stool sample incubation. These intact mass features were then further characterized by MS/MS.

#### Data analysis: targeted analysis

Extracted ion chromatograms (EIC) were extracted from raw data files using MassHunter Qualitative Data Analysis software and integrated using MassHunter Quantitative Data Analysis software. EICs were extracted using a 20-40 ppm window centered on the exact m/z value (**Supplemental Table 1**). For modifications with multiple detected isomers (e.g. aglycone hydroxylations), peak areas for each of the isomers were summed for a representative total EIC peak area. For compounds with available chemical standards, molar quantities were estimated by running a standard curve alongside samples.

### Quantification and Statistical Analysis

For untargeted metabolomics to identify mass features of interest, statistical analyses were performed on the output of XCMS peak quantification in Microsoft Excel for calculation of two-tailed Student’s t tests. Tracked mass features were integrated using MassHunter Quantitative Data Analysis software, with statistical analyses performed on integrated peak areas or estimated molar quantities using GraphPad Prism 10. For metagenomic analyses of complex communities, statistical analyses for relative strain abundances were performed in Microsoft Excel for calculation of two-tailed Student’s t tests. Calculated P values were adjusted for false discovery rate using the R package ‘qvalue’ version 2.42 (Storey et al., 2004).For statistical analysis of whole gut transit times in mice, two-tailed Student’s t tests (for solanine comparison with vehicle) and Dunnett’s multiple comparison test (for solanine and solanidine comparison with vehicle) were performed using GraphPad Prism 10. More details of statistical analyses and number of replicates are described in respective methods sections and figure captions.

## REFERENCES

1. Agarwal, S., Rao, A.V., 1998. Tomato lycopene and low density lipoprotein oxidation: A human dietary intervention study. Lipids 33, 981–984. 10.1007/s11745-998-0295-6

2. Almasieh, M., Zhou, Y., Kelly, M.E., Casanova, C., Di Polo, A., 2010. Structural and functional neuroprotection in glaucoma: role of galantamine-mediated activation of muscarinic acetylcholine receptors. Cell Death Dis. 1, e27–e27. 10.1038/cddis.2009.23

3. Almeida, A.F., Santos, C.N., Ventura, M.R., 2017. Synthesis of New Sulfated and Glucuronated Metabolites of Dietary Phenolic Compounds Identified in Human Biological Samples. J. Agric. Food Chem. 65, 6460–6466. 10.1021/acs.jafc.6b05629

4. Andrew Luckey, Edward Livingston, Yvette Tache, 2003. Mechanisms and Treatment of Postoperative Ileus. Arch. Surg. 138, 206. 10.1001/archsurg.138.2.206

5. Balks, H.J., Holst, J.J., Von Zur Mühlen, A., Brabant, G., 1997. Rapid Oscillations in Plasma Glucagon-Like Peptide-1 (GLP-1) in Humans: Cholinergic Control of GLP-1 Secretion via Muscarinic Receptors1. J. Clin. Endocrinol. Metab. 82, 786–790. 10.1210/jcem.82.3.3816

6. Billipp, T.E., Fung, C., Webeck, L.M., Sargent, D.B., Gologorsky, M.B., McDaniel, M.M., Kasal, D.N., McGinty, J.W., Barrow, K.A., Rich, L.M., Barilli, A., Sabat, M., Debley, J.S., Myers, R., Howitt, M.R., Von Moltke, J., 2023. Tuft cell-derived acetylcholine regulates epithelial fluid secretion (preprint). Immunology. 10.1101/2023.03.17.533208

7. Butelli, E., Titta, L., Giorgio, M., Mock, H.P., Matros, A., Peterek, S., Schijlen, E.G.W.M., Hall, R.D., Bovy, A.G., Luo, J., Martin, C., 2008. Enrichment of tomato fruit with health-promoting anthocyanins by expression of select transcription factors. Nat. Biotechnol. 26, 1301–1308. 10.1038/nbt.1506

8. Cahill, M.G., Caprioli, G., Vittori, S., James, K.J., 2010. Elucidation of the mass fragmentation pathways of potato glycoalkaloids and aglycons using Orbitrap mass spectrometry. J. Mass Spectrom. 45, 1019–1025. 10.1002/jms.1785

9. Caprioli, G., Cahill, M., Logrippo, S., James, K., 2015. Elucidation of the mass fragmentation pathways of tomatidine and β1-hydroxytomatine using orbitrap mass spectrometry. Nat. Prod. Commun. 10, 575–576. 10.1177/1934578x1501000409

10. Carter, M.M., Demis, D., Perelman, D., St. Onge, M., Petlura, C., Cunanan, K., Mathi, K., Maecker, H.T., Chow, J.M., Robinson, J.L., Sabag-Daigle, A., Sonnenburg, E.D., Buck, R.H., Gardner, C.D., Sonnenburg, J.L., 2025. A human milk oligosaccharide alters the microbiome, circulating hormones, and metabolites in a randomized controlled trial of older adults. Cell Rep. Med. 6, 102256. 10.1016/j.xcrm.2025.102256

11. Chagnon, F., Guay, I., Bonin, M.A., Mitchell, G., Bouarab, K., Malouin, F., Marsault, É., 2014. Unraveling the structure-activity relationship of tomatidine, a steroid alkaloid with unique antibiotic properties against persistent forms of Staphylococcus aureus. Eur. J. Med. Chem. 80, 605–620. 10.1016/j.ejmech.2013.11.019

12. Chaudhary, P., Sharma, A., Singh, B., Nagpal, A.K., 2018. Bioactivities of phytochemicals present in tomato. J. Food Sci. Technol. 55, 2833–2849. 10.1007/s13197-018-3221-z

13. Chaumeil, P.A., Mussig, A.J., Hugenholtz, P., Parks, D.H., 2020. GTDB-Tk: A toolkit to classify genomes with the genome taxonomy database. Bioinformatics 36, 1925–1927. 10.1093/bioinformatics/btz848

14. Cheng, A.G., Ho, P.-Y., Aranda-Díaz, A., Jain, S., Yu, F.B., Meng, X., Wang, M., Iakiviak, M., Nagashima, K., Zhao, A., Murugkar, P., Patil, A., Atabakhsh, K., Weakley, A., Yan, J., Brumbaugh, A.R., Higginbottom, S., Dimas, A., Shiver, A.L., Deutschbauer, A., Neff, N., Sonnenburg, J.L., Huang, K.C., Fischbach, M.A., 2022. Design, construction, and in vivo augmentation of a complex gut microbiome. Cell 185, 3617–3636.e19. 10.1016/j.cell.2022.08.003

15. Cheng, J., Venkatesh, S., Ke, K., Barratt, M.J., Gordon, J.I., 2024. A human gut *Faecalibacterium prausnitzii* fatty acid amide hydrolase. Science 386, eado6828. 10.1126/science.ado6828

16. Cichon, M.J., Riedl, K.M., Wan, L., Thomas-Ahner, J.M., Francis, D.M., Clinton, S.K., Schwartz, S.J., 2017. Plasma Metabolomics Reveals Steroidal Alkaloids as Novel Biomarkers of Tomato Intake in Mice. Mol. Nutr. Food Res. 61, 1–9. 10.1002/mnfr.201700241

17. Clark, R.L., Connors, B.M., Stevenson, D.M., Hromada, S.E., Hamilton, J.J., Amador-Noguez, D., Venturelli, O.S., 2021. Design of synthetic human gut microbiome assembly and butyrate production. Nat. Commun. 12, 3254. 10.1038/s41467-021-22938-y

18. Cockburn, D.W., Koropatkin, N.M., 2016. Polysaccharide Degradation by the Intestinal Microbiota and Its Influence on Human Health and Disease. J. Mol. Biol. 428, 3230–3252. 10.1016/j.jmb.2016.06.021

19. Cockcroft, D.W., Gault, H., 1976. Prediction of Creatinine Clearance from Serum Creatinine. Nephron 16, 31–41. 10.1159/000180580

20. Croswell, A., Amir, E., Teggatz, P., Barman, M., Salzman, N.H., 2009. Prolonged impact of antibiotics on intestinal microbial ecology and susceptibility to enteric Salmonella infection. Infect. Immun. 77, 2741–2753. 10.1128/IAI.00006-09

21. Day, A.J., DuPont, M.S., Ridley, S., Rhodes, M., Rhodes, M.J.C., Morgan, M.R.A., Williamson, G., 1998. Deglycosylation of flavonoid and isoflavonoid glycosides by human small intestine and liver β-glucosidase activity. FEBS Lett. 436, 71–75. 10.1016/S0014-5793(98)01101-6

22. Do, D., Casperson, S.L., Cooperstone, J., 2024. Assessing Plasma Accumulation of Steroidal Alkaloids and Their Metabolites in Healthy Adults Following Long-Term Tomato-Based Vegetable Juice Consumption. Curr. Dev. Nutr. 8, 102675. 10.1016/j.cdnut.2024.102675

23. Dodd, D., Spitzer, M.H., Van Treuren, W., Merrill, B.D., Hryckowian, A.J., Higginbottom, S.K., Le, A., Cowan, T.M., Nolan, G.P., Fischbach, M.A., Sonnenburg, J.L., 2017. A gut bacterial pathway metabolizes aromatic amino acids into nine circulating metabolites. Nature 551, 648–652. 10.1038/nature24661

24. Dyle, M.C., Ebert, S.M., Cook, D.P., Kunkel, S.D., Fox, D.K., Bongers, K.S., Bullard, S.A., Dierdorff, J.M., Adams, C.M., 2014. Systems-based discovery of tomatidine as a natural small molecule inhibitor of skeletal muscle atrophy. J. Biol. Chem. 289, 14913–14924. 10.1074/jbc.M114.556241

25. Dzakovich, M.P., Goggans, M.L., Thomas-Ahner, J.M., Moran, N.E., Clinton, S.K., Francis, D.M., Cooperstone, J.L., 2024. Transcriptomics and Metabolomics Reveal Tomato Consumption Alters Hepatic Xenobiotic Metabolism and Induces Steroidal Alkaloid Metabolite Accumulation in Mice. Mol. Nutr. Food Res. 68, 2300239. 10.1002/mnfr.202300239

26. Economic Research Service (ERS), U.S. Department of Agriculture (USDA), n.d. Food Availability (Per Capita) Data System.

27. Ellman, G.L., Courtney, K.D., Andres, V., Featherstone, R.M., 1961. A new and rapid colorimetric determination of acetylcholinesterase activity. Biochem. Pharmacol. 7, 88–95. 10.1016/0006-2952(61)90145-9

28. Eyssen, H.J., Parmentier, G.G., Compernolle, F.C., De Pauw, G., Piessens-Denef, M., 1973. Biohydrogenation of Sterols by *Eubacterium* ATCC 21,408— *Nova* Species. Eur. J. Biochem. 36, 411–421. 10.1111/j.1432-1033.1973.tb02926.x

29. Fahey, J.W., Zhang, Y., Talalay, P., 1997. Broccoli sprouts: An exceptionally rich source of inducers of enzymes that protect against chemical carcinogens. Proc. Natl. Acad. Sci. U. S. A. 94, 10367–10372. 10.1073/pnas.94.19.10367

30. Falany, C.N., Wheeler, J., Tae Sung Oh, Falany, J.L., 1994. Steroid sulfation by expressed human cytosolic sulfotransferases. J. Steroid Biochem. Mol. Biol. 48, 369–375. 10.1016/0960-0760(94)90077-9

31. Freier, T.A., Beitz, D.C., Li, L., Hartman, P.A., 1994. Characterization of Eubacterium coprostanoligenes sp. nov., a Cholesterol-Reducing Anaerobe. Int. J. Syst. Bacteriol. 44, 137–142. 10.1099/00207713-44-1-137

32. Friedman, M., Kozukue, N., Harden, L.A., 1998. Preparation and Characterization of Acid Hydrolysis Products of the Tomato Glycoalkaloid α-Tomatine. J. Agric. Food Chem. 46, 2096–2101. 10.1021/jf970898k

33. Friedman, M., Levin, C.E., Lee, S.U., Kim, H.J., Lee, I.S., Byun, J.O., Kozukue, N., 2009. Tomatine-containing green tomato extracts inhibit growth of human breast, colon, liver, and stomach cancer cells. J. Agric. Food Chem. 57, 5727–5733. 10.1021/jf900364j

34. Fuhrman, B.J., Teter, B.E., Barba, M., Byrne, C., Cavalleri, A., Grant, B.J., Horvath, P.J., Morelli, D., Venturelli, E., Muti, P.C., 2008. Equol Status Modifies the Association of Soy Intake and Mammographic Density in a Sample of Postmenopausal Women. Cancer Epidemiol. Biomarkers Prev. 17, 33–42. 10.1158/1055-9965.EPI-07-0193

35. Furness, J.B., 2012. The enteric nervous system and neurogastroenterology. Nat. Rev. Gastroenterol. Hepatol. 9, 286–294. 10.1038/nrgastro.2012.32

36. Gérard, P., Lepercq, P., Leclerc, M., Gavini, F., Raibaud, P., Juste, C., 2007. *Bacteroides sp.* Strain D8, the First Cholesterol-Reducing Bacterium Isolated from Human Feces. Appl. Environ. Microbiol. 73, 5742–5749. 10.1128/AEM.02806-06

37. Giovannucci, E., 1999. Tomatoes, Tomato-Based Products, Lycopene, and Cancer: Review of the Epidemiologic Literature. JNCI J. Natl. Cancer Inst. 91, 317–331. 10.1093/jnci/91.4.317

38. Goyal, R.K., Hirano, I., 1996. The Enteric Nervous System. N. Engl. J. Med. 334, 1106–1115. 10.1056/NEJM199604253341707

39. Han, B.H., Park, M.H., Han, Y.N., 1982. Degradation of ginseng saponins under mild acidic conditions. Planta Med. 44, 146–149. 10.1055/s-2007-971425

40. Hang, S., Paik, D., Yao, L., Kim, E., Jamma, T., Lu, J., Ha, S., Nelson, B.N., Kelly, S.P., Wu, L., Zheng, Y., Longman, R.S., Rastinejad, F., Devlin, A.S., Krout, M.R., Fischbach, M.A., Littman, D.R., Huh, J.R., 2019. Bile acid metabolites control TH17 and Treg cell differentiation. Nature 576, 143–148. 10.1038/s41586-019-1785-z

41. Hennessy, R.C., Jørgensen, N.O.G., Scavenius, C., Enghild, Jan. J., Greve-Poulsen, M., Sørensen, O.B., Stougaard, P., 2018. A Screening Method for the Isolation of Bacteria Capable of Degrading Toxic Steroidal Glycoalkaloids Present in Potato. Front. Microbiol. 9, 2648. 10.3389/fmicb.2018.02648

42. Hövelmann, Y., Lewin, L., Steinert, K., Hübner, F., Humpf, H., 2020. Mass Spectrometry–Based Analysis of Urinary Biomarkers for Dietary Tomato Intake. Mol. Nutr. Food Res. 64, 2000011. 10.1002/mnfr.202000011

43. Hubatsch, I., Ragnarsson, E.G.E., Artursson, P., 2007. Determination of drug permeability and prediction of drug absorption in Caco-2 monolayers. Nat. Protoc. 2, 2111–2119. 10.1038/nprot.2007.303

44. Ibach, B., Haen, E., 2004. Acetylcholinesterase Inhibition in Alzheimers Disease. Curr. Pharm. Des. 10, 231–251. 10.2174/1381612043386509

45. Johnson, E.L., Heaver, S.L., Waters, J.L., Kim, B.I., Bretin, A., Goodman, A.L., Gewirtz, A.T., Worgall, T.S., Ley, R.E., 2020. Sphingolipids produced by gut bacteria enter host metabolic pathways impacting ceramide levels. Nat. Commun. 11, 1–11. 10.1038/s41467-020-16274-w

46. Kacmaz, H., Alto, A., Knutson, K., Linden, D.R., Gibbons, S.J., Farrugia, G., Beyder, A., 2021. A simple automated approach to measure mouse whole gut transit. Neurogastroenterol. Motil. 33, e13994. 10.1111/nmo.13994

47. Kaup, O., Gräfen, I., Zellermann, E.M., Eichenlaub, R., Gartemann, K.H., 2005. Identification of a tomatinase in the tomato-pathogenic actinomycete Clavibacter michiganensis subsp. michiganensis NCPPB382. Mol. Plant. Microbe Interact. 18, 1090–1098. 10.1094/MPMI-18-1090

48. Kenny, D.J., Plichta, D.R., Shungin, D., Koppel, N., Hall, A.B., Fu, B., Vasan, R.S., Shaw, S.Y., Vlamakis, H., Balskus, E.P., Xavier, R.J., 2020. Cholesterol Metabolism by Uncultured Human Gut Bacteria Influences Host Cholesterol Level. Cell Host Microbe 28, 245–257.e6. 10.1016/j.chom.2020.05.013

49. Kim, D.H., 2018. Gut microbiota-mediated pharmacokinetics of ginseng saponins. J. Ginseng Res. 42, 255–263. 10.1016/j.jgr.2017.04.011

50. King, R.R., McQueen, R.E., 1981. Transformations of Potato Glycoalkaloids by Rumen Microorganisms. J. Agric. Food Chem. 29, 1101–1103. 10.1021/jf00107a056

51. Koeth, R.A., Wang, Z., Levison, B.S., Buffa, J.A., Org, E., Sheehy, B.T., Britt, E.B., Fu, X., Wu, Y., Li, L., Smith, J.D., DiDonato, J.A., Chen, J., Li, H., Wu, G.D., Lewis, J.D., Warrier, M., Brown, J.M., Krauss, R.M., Tang, W.H.W., Bushman, F.D., Lusis, A.J., Hazen, S.L., 2013. Intestinal microbiota metabolism of l-carnitine, a nutrient in red meat, promotes atherosclerosis. Nat. Med. 19, 576–585. 10.1038/nm.3145

52. Kovatcheva-Datchary, P., Shoaie, S., Lee, S., Wahlström, A., Nookaew, I., Hallen, A., Perkins, R., Nielsen, J., Bäckhed, F., 2019. Simplified Intestinal Microbiota to Study Microbe-Diet-Host Interactions in a Mouse Model. Cell Rep. 26, 3772–3783.e6. 10.1016/j.celrep.2019.02.090

53. Kreis, M.E., Kasparek, M., Zittel, T.T., Becker, H.D., Jehle, E.C., 2001. Neostigmine increases postoperative colonic motility in patients undergoing colorectal surgery. Surgery 130, 449–456. 10.1067/msy.2001.116451

54. Krupp, D., Doberstein, N., Shi, L., Remer, T., 2012. Hippuric Acid in 24-Hour Urine Collections Is a Potential Biomarker for Fruit and Vegetable Consumption in Healthy Children and Adolescents,. J. Nutr. 142, 1314–1320. 10.3945/jn.112.159319

55. Kúdelová, J., Seifrtová, M., Suchá, L., Tomšík, P., Havelek, R., Řezáčová, M., 2013. Alpha-tomatine activates cell cycle checkpoints in the absence of DNA damage in human leukemic MOLT-4 cells. J. Appl. Biomed. 11, 93–103. 10.2478/v10136-012-0033-8

56. Kuo, C.Y., Huang, W.C., Liou, C.J., Chen, L.C., Shen, J.J., Kuo, M.L., 2017. Tomatidine Attenuates Airway Hyperresponsiveness and Inflammation by Suppressing Th2 Cytokines in a Mouse Model of Asthma. Mediators Inflamm. 2017. 10.1155/2017/5261803

57. Kuss, S.K., Best, G.T., Etheredge, C.A., Pruijssers, A.J., Frierson, J.M., Hooper, L.V., Dermody, T.S., Pfeiffer, J.K., 2011. Intestinal Microbiota Promote Enteric Virus Replication and Systemic Pathogenesis. Science 334, 249–252. 10.1126/science.1211057

58. Letunic, I., Bork, P., 2024. Interactive Tree of Life (iTOL) v6: recent updates to the phylogenetic tree display and annotation tool. Nucleic Acids Res. 52, W78–W82. 10.1093/nar/gkae268

59. Liou, C.S., Sirk, S.J., Diaz, C.A.C., Klein, A.P., Fischer, C.R., Higginbottom, S.K., Erez, A., Donia, M.S., Sonnenburg, J.L., Sattely, E.S., 2020. A Metabolic Pathway for Activation of Dietary Glucosinolates by a Human Gut Symbiont. Cell 180, 717–728.e19. 10.1016/j.cell.2020.01.023

60. Mair, R.D., Sirich, T.L., Plummer, N.S., Meyer, T.W., 2018. Characteristics of Colon-Derived Uremic Solutes. Clin. J. Am. Soc. Nephrol. 13, 1398–1404. 10.2215/CJN.03150318

61. Marcobal, A., Kashyap, P.C., Nelson, T.A., Aronov, P.A., Donia, M.S., Spormann, A., Fischbach, M.A., Sonnenburg, J.L., 2013. A metabolomic view of how the human gut microbiota impacts the host metabolome using humanized and gnotobiotic mice. ISME J. 7, 1933–1943. 10.1038/ismej.2013.89

62. Martin, C., Zhang, Y., Tonelli, C., Petroni, K., 2013. Plants, diet, and health. Annu. Rev. Plant Biol. 64, 19–46. 10.1146/annurev-arplant-050312-120142

63. McMillan, M., Thompson, J.C., 1979. An outbreak of suspected solanine poisoning in schoolboys: examination of criteria of solanine poisoning. QJM Int. J. Med. 48. 10.1093/oxfordjournals.qjmed.a067573

64. Mensinga, T.T., Sips, A.J.A.M., Rompelberg, C.J.M., Van Twillert, K., Meulenbelt, J., Van Den Top, H.J., Van Egmond, H.P., 2005. Potato glycoalkaloids and adverse effects in humans: An ascending dose study. Regul. Toxicol. Pharmacol. 41, 66–72. 10.1016/j.yrtph.2004.09.004

65. Murphy, M.M., Barraj, L.M., Herman, D., Bi, X., Cheatham, R., Randolph, R.K., 2012. Phytonutrient Intake by Adults in the United States in Relation to Fruit and Vegetable Consumption. J. Acad. Nutr. Diet. 112, 222–229. 10.1016/j.jada.2011.08.044

66. Nagashima, K., Zhao, A., Atabakhsh, K., Bae, M., Blum, J.E., Weakley, A., Jain, S., Meng, X., Cheng, A.G., Wang, M., Higginbottom, S., Dimas, A., Murugkar, P., Sattely, E.S., Moon, J.J., Balskus, E.P., Fischbach, M.A., 2023. Mapping the T cell repertoire to a complex gut bacterial community. Nature 621, 162–170. 10.1038/s41586-023-06431-8

67. Nigg, H.N., Ramos, L.E., Graham, E.M., Sterling, J., Brown, S., Cornell, J.A., 1996. Inhibition of Human Plasma and Serum Butyrylcholinesterase (EC 3.1.1.8) by α -Chaconine and α - Solanine. Toxicol. Sci. 33, 272–281. 10.1093/toxsci/33.2.272

68. Nohara, T., Ono, M., Ikeda, T., Fujiwara, Y., El-Aasr, M., 2010. The tomato saponin, esculeoside A. J. Nat. Prod. 73, 1734–1741. 10.1021/np100311t

69. Norred, W., Nishie, K., Osman, S., 1976. Excretion, distribution and metabolic fate of 3H-alpha-chaconine. Res. Commun. Chem. Pathol. Pharmacol. 13.

70. Osbourn, A., Bowyer, Paul, Lunness, Patricia, Clarke, Belinda, Daniels, Michael, 1995. Fungal Pathogens of Oat Roots and Tomato Leaves Employ Closely Related Enzymes to Detoxify Different Host Plant Saponins. Mol. Plant. Microbe Interact. 8, 971. 10.1094/MPMI-8-0971

71. Panevska, A., Skočaj, M., Križaj, I., Maček, P., Sepčić, K., 2019. Ceramide phosphoethanolamine, an enigmatic cellular membrane sphingolipid. Biochim. Biophys. Acta - Biomembr. 1861, 1284–1292. 10.1016/j.bbamem.2019.05.001

72. Patil, B.C., Sharma, R.P., Salunkhe, D.K., Salunkhe, K., 1972. Evaluation of solanine toxicity. Food Cosmet. Toxicol. 10, 395–398. 10.1016/S0015-6264(72)80258-X

73. Plageman, L.R., Pauletti, G.M., Skau, K.A., 2002. Characterization of acetylcholinesterase in Caco-2 cells. Exp. Biol. Med. 227, 480–486. 10.1177/153537020222700712

74. Rakoff-Nahoum, S., Coyne, M.J., Comstock, L.E., 2014. An Ecological Network of Polysaccharide Utilization among Human Intestinal Symbionts. Curr. Biol. 24, 40–49. 10.1016/j.cub.2013.10.077

75. Rakoff-Nahoum, S., Paglino, J., Eslami-Varzaneh, F., Edberg, S., Medzhitov, R., 2004. Recognition of Commensal Microflora by Toll-Like Receptors Is Required for Intestinal Homeostasis. Cell 118, 229–241. 10.1016/j.cell.2004.07.002

76. Roddick, J.G., 1989. The acetylcholinesterase-inhibitory activity of steroidal glycoalkaloids and their aglycones. Phytochemistry 28, 2631–2634. 10.1016/S0031-9422(00)98055-5

77. Romano, K.A., Vivas, E.I., Amador-Noguez, D., Rey, F.E., 2015. Intestinal Microbiota Composition Modulates Choline Bioavailability from Diet and Accumulation of the Proatherogenic Metabolite Trimethylamine- *N* -Oxide. mBio 6, e02481–14. 10.1128/mBio.02481-14

78. Schieber, A., Saldaña, M., 2009. Potato Peels: A Source of Nutritionally and Pharmacologically Interesting Compounds – A Review. Food 3, 23–29.

79. Schneider, H., Schwiertz, A., Collins, M.D., Blaut, M., 1999. Anaerobic transformation of quercetin-3-glucoside by bacteria from the human intestinal tract. Arch. Microbiol. 171, 81–91. 10.1007/s002030050682

80. Sesso, H.D., Liu, S., Gaziano, J.M., Buring, J.E., 2003. Dietary Lycopene, Tomato-Based Food Products and Cardiovascular Disease in Women. J. Nutr. 133, 2336–2341. 10.1093/jn/133.7.2336

81. Shieh, J.M., Cheng, T.H., Shi, M.D., Wu, P.F., Chen, Y., Ko, S.C., Shih, Y.W., 2011. α-Tomatine Suppresses Invasion and Migration of Human Non-Small Cell Lung Cancer NCI-H460 Cells Through Inactivating FAK/PI3K/Akt Signaling Pathway and Reducing Binding Activity of NF-κB. Cell Biochem. Biophys. 60, 297–310. 10.1007/s12013-011-9152-1

82. Sholola, M.J., Goggans, M.L., Dzakovich, M.P., Francis, D.M., Jacobi, S.K., Cooperstone, J.L., 2025. Discovery of steroidal alkaloid metabolites and their accumulation in pigs after short-term tomato consumption. Food Chem. 463, 141346. 10.1016/j.foodchem.2024.141346

83. Simó, C., García-Cañas, V., 2020. Dietary bioactive ingredients to modulate the gut microbiota-derived metabolite TMAO. New opportunities for functional food development. Food Funct. 11, 6745–6776. 10.1039/D0FO01237H

84. Slanina, P., 1990. Solanine (glycoalkaloids) in potatoes: Toxicological evaluation. Food Chem. Toxicol. 28, 759–761. 10.1016/0278-6915(90)90074-W

85. Storey, J.D., Taylor, J.E., Siegmund, D., 2004. Strong Control, Conservative Point Estimation and Simultaneous Conservative Consistency of False Discovery Rates: A Unified Approach. J. R. Stat. Soc. Ser. B Stat. Methodol. 66, 187–205. 10.1111/j.1467-9868.2004.00439.x

86. Tautenhahn, R., Patti, G.J., Rinehart, D., Siuzdak, G., 2012. XCMS Online: A Web-Based Platform to Process Untargeted Metabolomic Data. Anal. Chem. 84, 5035–5039. 10.1021/ac300698c

87. Tennant, D.R., Davidson, J., Day, A.J., 2014. Phytonutrient intakes in relation to European fruit and vegetable consumption patterns observed in different food surveys. Br. J. Nutr. 112, 1214–1225. 10.1017/S0007114514001950

88. Turnbaugh, P.J., Ley, R.E., Mahowald, M.A., Magrini, V., Mardis, E.R., Gordon, J.I., 2006. An obesity-associated gut microbiome with increased capacity for energy harvest. Nature 444, 1027–1031. 10.1038/nature05414

89. Wang, M., Osborn, L.J., Jain, S., Meng, X., Weakley, A., Yan, J., Massey, W.J., Varadharajan, V., Horak, A., Banerjee, R., Allende, D.S., Chan, E.R., Hajjar, A.M., Wang, Z., Dimas, A., Zhao, A., Nagashima, K., Cheng, A.G., Higginbottom, S., Hazen, S.L., Brown, J.M., Fischbach, M.A., 2023. Strain dropouts reveal interactions that govern the metabolic output of the gut microbiome. Cell 186, 2839–2852.e21. 10.1016/j.cell.2023.05.037

90. Wang, W., Xiao, G., Du, G., Chang, L., Yang, Y., Ye, J., Chen, B., 2022. Glutamicibacter halophytocola-mediated host fitness of potato tuber moth on Solanaceae crops. Pest Manag. Sci. 78, 3920–3930. 10.1002/ps.6955

91. Wu, M., McNulty, N.P., Rodionov, D.A., Khoroshkin, M.S., Griffin, N.W., Cheng, J., Latreille, P., Kerstetter, R.A., Terrapon, N., Henrissat, B., Osterman, A.L., Gordon, J.I., 2015. Genetic determinants of in vivo fitness and diet responsiveness in multiple human gut *Bacteroides*. Science 350, aac5992. 10.1126/science.aac5992

92. Yajima, T., Inoue, R., Matsumoto, M., Yajima, M., 2011. Non-neuronal release of ACh plays a key role in secretory response to luminal propionate in rat colon. J. Physiol. 589, 953–962. 10.1113/jphysiol.2010.199976

93. Yao, L., D’Agostino, G.D., Park, J., Hang, S., Adhikari, A.A., Zhang, Y., Li, W., Avila-Pacheco, J., Bae, S., Clish, C.B., Franzosa, E.A., Huttenhower, C., Huh, J.R., Devlin, A.S., 2022. A biosynthetic pathway for the selective sulfonation of steroidal metabolites by human gut bacteria. Nat. Microbiol. 7, 1404–1418. 10.1038/s41564-022-01176-y

94. Yu, H., Xia, H., Tang, Q., Xu, H., Wei, G., Chen, Y., Dai, X., Gong, Q., Bi, F., 2017. Acetylcholine acts through M3 muscarinic receptor to activate the EGFR signaling and promotes gastric cancer cell proliferation. Sci. Rep. 7, 40802. 10.1038/srep40802

95. Zhao, L., Zhang, F., Ding, X., Wu, G., Lam, Y.Y., Wang, Xuejiao, Fu, H., Xue, X., Lu, C., Ma, J., Yu, L., Xu, C., Ren, Z., Xu, Y., Xu, S., Shen, H., Zhu, X., Shi, Y., Shen, Q., Dong, W., Liu, R., Ling, Y., Zeng, Y., Wang, Xingpeng, Zhang, Q., Wang, J., Wang, L., Wu, Y., Zeng, B., Wei, H., Zhang, M., Peng, Y., Zhang, C., 2018. Gut bacteria selectively promoted by dietary fibers alleviate type 2 diabetes. Science 359, 1151–1156. 10.1126/science.aao5774

96. Zhu, W., Gregory, J.C., Org, E., Buffa, J.A., Gupta, N., Wang, Z., Li, L., Fu, X., Wu, Y., Mehrabian, M., Sartor, R.B., McIntyre, T.M., Silverstein, R.L., Tang, W.H.W., Didonato, J.A., Brown, J.M., Lusis, A.J., Hazen, S.L., 2016. Gut Microbial Metabolite TMAO Enhances Platelet Hyperreactivity and Thrombosis Risk. Cell 165, 111–124. 10.1016/j.cell.2016.02.011

