## supplemental materials for "Gut microbiota gate host exposure to cholinesterase inhibitors from dietary Solanums"

##### SUPPLEMENTAL INFORMATION

### SUPPLEMENTAL FIGURES

A)

| Criteria | Number of Mass Features HILIC | C18 |
|---|---|---|
| Total | 7787 | 9968 |
| Retention > 1 min | 4876 | 9959 |
| p value | p<0.10: 649 | p<0.05: 663 |
| Fold > 3 | 66 | 101 |
| Higher in tomato fed | 49 | 72 |
| Peak height > $10^4$ | 28 | 29 |
| Parent features | 14 | 23 |

B)

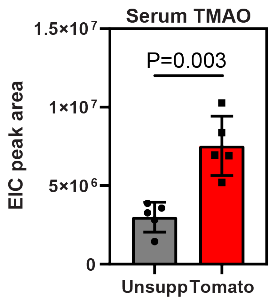


C)

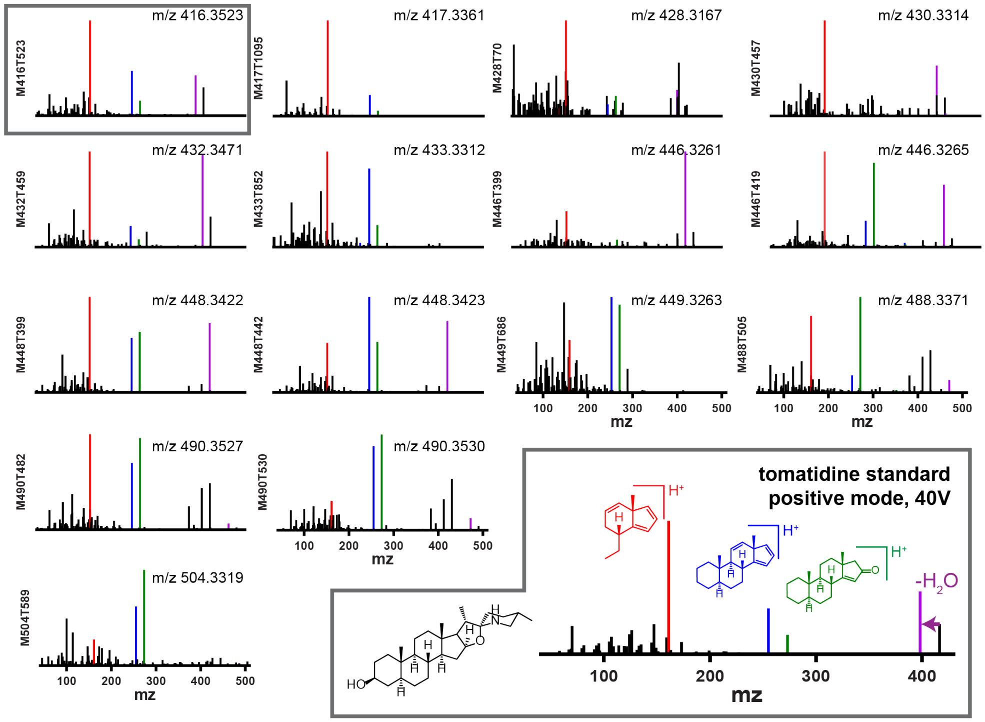


**Supplemental Figure 1. Host relevant metabolites from tomato dietary intervention. Relates to Figure 1.**

A) Filtering criteria for untargeted metabolomics to identify diet-derived mass features present in mouse serum following tomato consumption. Mass features were identified and compiled using xcms for data collected in positive mode ionization with HILIC or C18 chromatography.

B) Abundance of trimethylamine N-oxide (TMAO) in the serum of mice receiving tomatoes (red bars) and unsupplemented diet (grey bars). Values shown are the mean±SD of EIC peak areas for $[M+H]^+$ *m/z* 76.0757 for each treatment group (n=5), with individual replicates overlaid.

C) Tandem MS (MS/MS) spectra collected at 40V in positive mode for SA derived metabolites observed in mouse serum following tomato-supplemented diet. The MS/MS spectrum for a tomatidine standard is shown on the bottom right, with characteristic fragments colored and annotated with putative structures. Conserved fragments (±2 amu) are colored similarly in each spectrum. The serum MS/MS spectrum in the upper left boxed in grey is a match to the spectrum for the tomatidine standard.

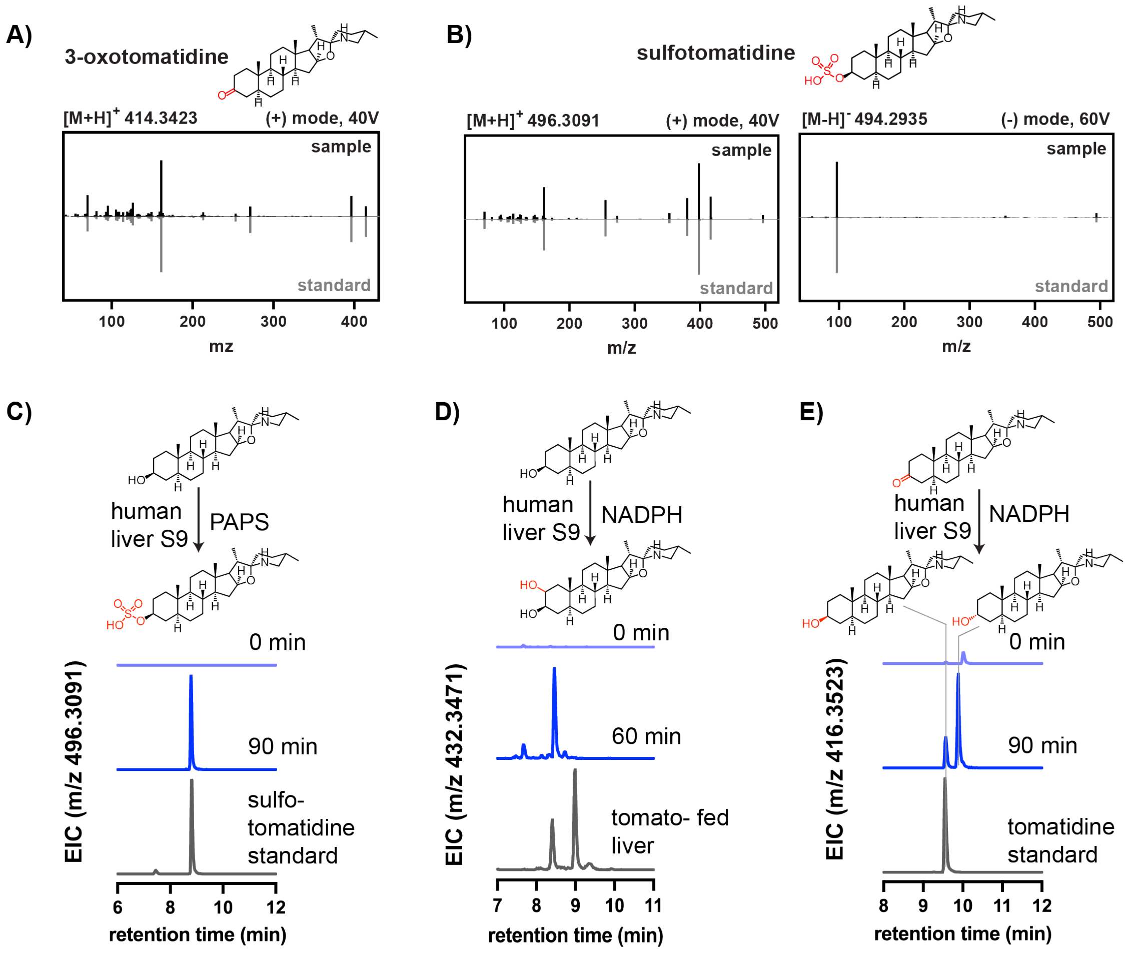


**Supplemental Figure 2. Characterization of host relevant steroidal alkaloids from tomato. Relates to Figure 1.**

A) MS/MS fragmentation of 3-oxotomatidine in mouse liver in ($[M+H]^+$ *m/z* 414.3423, positive mode, 40V) (black) compared to the MS/MS spectrum of a synthesized standard (grey)
B) MS/MS fragmentation of tomatidine sulfate in mouse liver ($[M+H]^+$ *m/z* 496.3091, positive mode, 40V and $[M-H]^-$ *m/z* 494.2935, negative mode, 60V) (black) compared to the MS/MS spectrum of a synthesized standard (grey)
C) Production of tomatidine sulfate by human liver protein extracts before and after incubation with tomatidine and the sulfate donor PAPS for 90 minutes. Data shown are EICs for the mass of tomatidine sulfate in positive mode ($[M+H]^+$ *m/z* 496.3091).
D) Production of hydroxytomatidine by human liver protein extracts before and after incubation with tomatidine and NADPH for 60 minutes. Data shown are EICs for the mass of hydroxytomatidine in positive mode ($[M+H]^+$ *m/z* 432.3471), alongside a chromatograph for the mass in mouse liver following tomato feeding. The structure shown is proposed; the position of hydroxylation is not known.
E) Production of tomatidine and a putative 3αOH tomatidine epimer by human liver protein extracts before and after incubation with 3-oxotomatidine and NADPH for 90 minute. Data shown are EICs for the mass of tomatidine in positive mode ($[M+H]^+$ *m/z* 416.3523), alongside a chromatograph for an authentic tomatidine standard.

A)

| Criteria | Mass Features C18 |
|---|---|
| Total | 29085 |
| Retention > 1 min | 29081 |
| peak width < 5s | 20895 |
| p < 0.05 | 1527 |
| Fold change > 3 | 347 |
| Higher in potato fed | 285 |
| Peak height > $10^4$ | 132 |
| Parent features | 40 |

B)

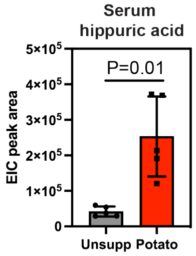


C)

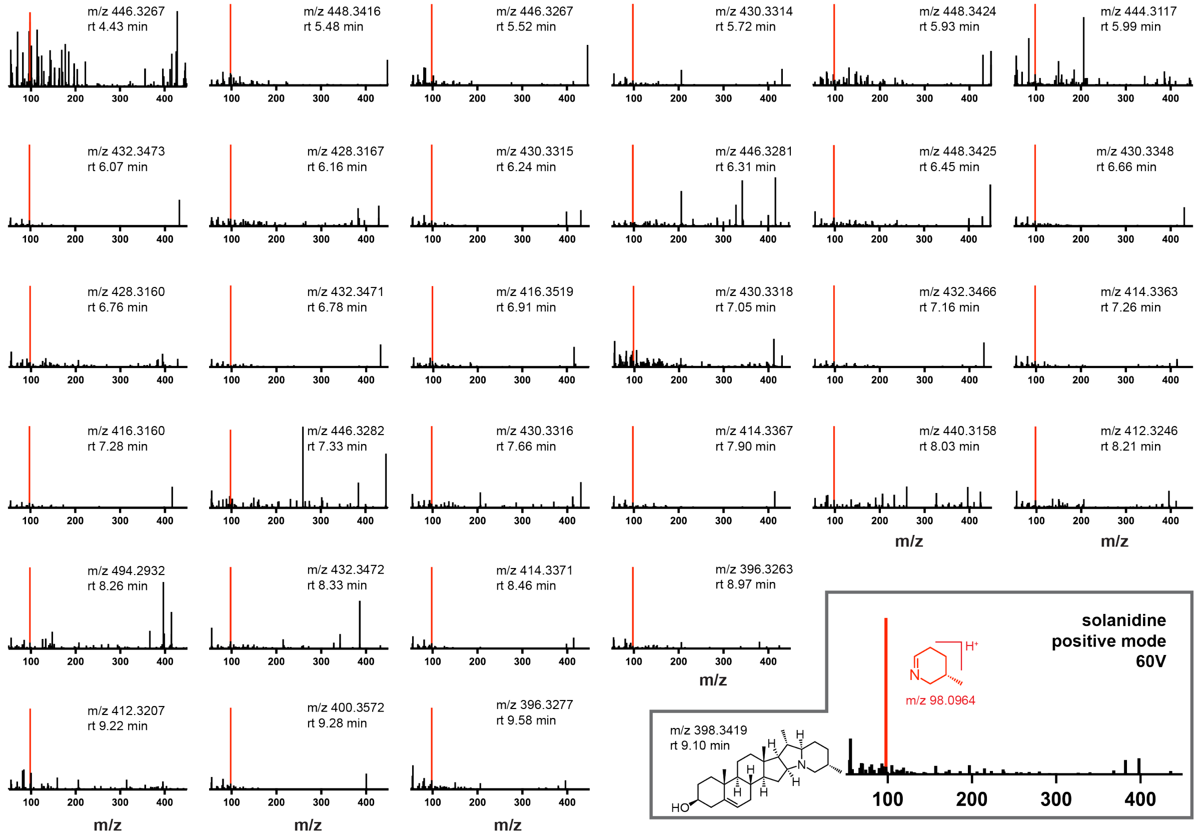


**Supplemental Figure 3. Host relevant metabolites from potato dietary intervention. Relates to Figure 1.**

A) Filtering criteria for untargeted metabolomics to identify diet-derived mass features present in mouse serum following potato consumption. Mass features were identified and compiled using xcms for LC-MS data collected in positive mode ionization with C18 chromatography.

B) Abundance of hippuric acid in the serum of mice receiving potatoes (red bars) and unsupplemented diet (grey bars). Values shown are the mean±SD of EIC peak areas for $[M+H]^+$ *m/z* 180.0655 for each treatment group (n=5), with individual replicates overlaid.

C) MS/MS spectra collected at 60V in positive mode for SA metabolites in mouse serum following potato-supplemented diet. The MS/MS spectrum for a solanidine standard is shown on the bottom right, with the characteristic fragment in red and annotated with putative structures. Conserved fragments are colored similarly for each spectrum.

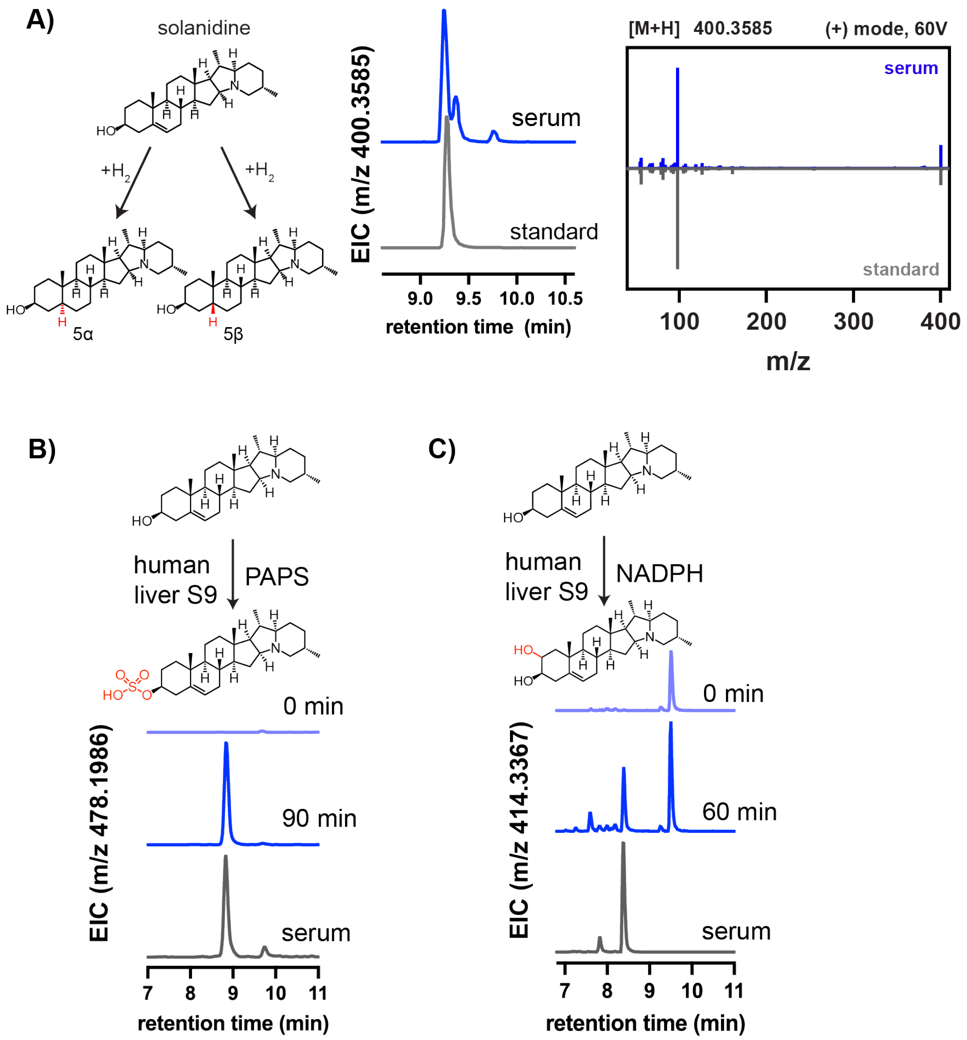


**Supplemental Figure 4. Characterization of host relevant steroidal alkaloids from potato. Relates to Figure 1.**

A) MS/MS fragmentation and EIC of dihydrosolanidine from mouse serum in ($[M+H]^+$ *m/z* 400.3585, positive mode, 60V) (blue) compared with those of a synthesized standard (grey)

B) Production of solanidine sulfate by human liver protein extracts before and after incubation with solanidine and sulfate donor PAPS for 90 minutes. Data shown are EICs for the mass of solanidine sulfate in positive mode ($[M+H]^+$ *m/z* 478.1986), alongside a chromatograph for the mass in mouse serum following potato feeding

C) Production of hydroxysolanidine by human liver protein extracts before and after incubation with tomatidine and NADPH for 60 minutes. Data shown are EICs for the mass of hydroxysolanidine in positive mode ($[M+H]^+$ *m/z* 414.3367), alongside a chromatograph for the mass in mouse serum following potato feeding. The position of hydroxylation is not known.

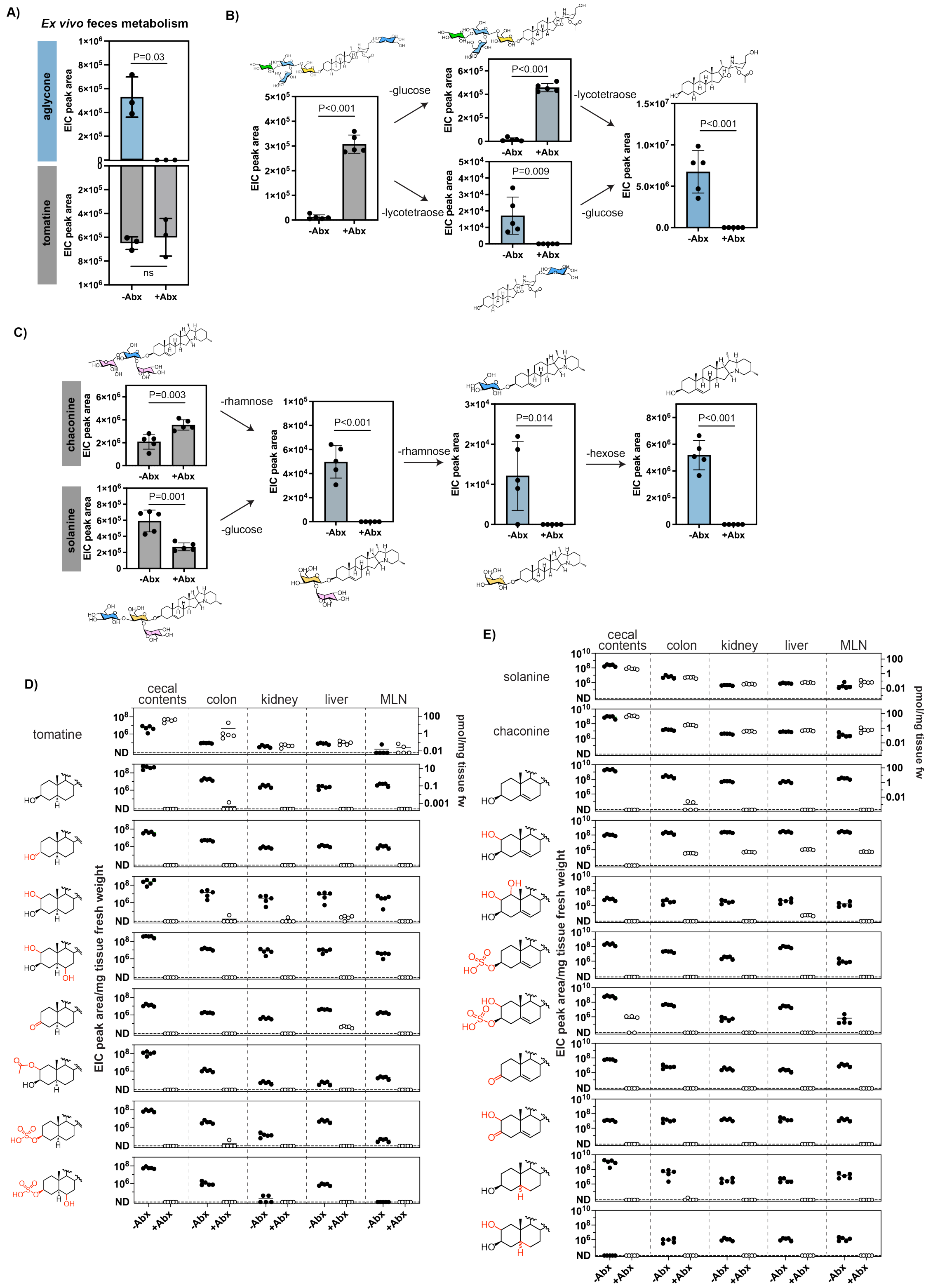


**Supplemental Figure 5. Gut microbiota gate SGA metabolism *in vivo*. Relates to Figure 2**

A) *Ex vivo* tomatine metabolism by feces from conventional Swiss Webster mice receiving water (-Abx) or antibiotics solution (+Abx). Fecal extracts from n=3 mice were incubated with 20 μM tomatine for 36 hours. Values shown are the mean±SD EIC peak areas of the glycoside substrate tomatine and the aglycone product tomatidine with individual biological replicates overlaid. Each individual biological replicate shown is the average of three replicate incubations. P values determined using a two-tailed t test. ns = not significant (P>0.05).

B) Esculeoside A metabolites measured in cecal contents from Swiss Webster mice receiving water (-Abx) or antibiotics (+Abx) alongside 10% (w/w) tomato-supplemented diet. Values shown are mean±SD EIC peak areas with individual replicates overlaid (n=5). Putative structures according to *m/z* values and MS/MS spectra are shown. P values determined using a two-tailed t test.

C) Partially deglycosylated potato SGA metabolites measured in cecal contents from Swiss Webster mice receiving water (-Abx) or antibiotics (+Abx) alongside 5% (w/w) potato-supplemented diet. Values shown are mean±SD EIC peak areas with individual replicates overlaid (n=5). Putative structures according to *m/z* values and MS/MS spectra are shown, with multiple structures drawn for peaks with more than one possible assignment. P values determined using a two-tailed t test.

D) Accumulation of tomatine-derived metabolites in tissues following dietary intervention with 10% (w/w) tomato diet in C57BL/6J mice receiving water (filled circles) or antibiotics (unfilled circles). Metabolite EIC peak areas normalized to tissue fresh weight are represented for individual animals (n=5) with the mean value for each group indicated by a line. Molar quantities for compounds with commercial standards are indicated on the right axes where applicable. For molecules with multiple detected isomers (+OH and +2OH aglycones), the sum of peak areas is shown. ND = not detectable.

E) Accumulation of glycoalkaloid-derived metabolites in tissues following dietary intervention with 5% (w/w) potato diet in C57BL/6J mice receiving water (filled circles) or antibiotics (unfilled circles). Metabolite EIC peak areas normalized to tissue fresh weight are represented for individual animals (n=5) with the mean value for each group indicated by a line. Molar quantities for compounds with commercial standards are indicated on the right axes where applicable. For molecules with multiple detected isomers (+OH and +2OH aglycones), the sum of peak areas is shown. ND = not detectable.

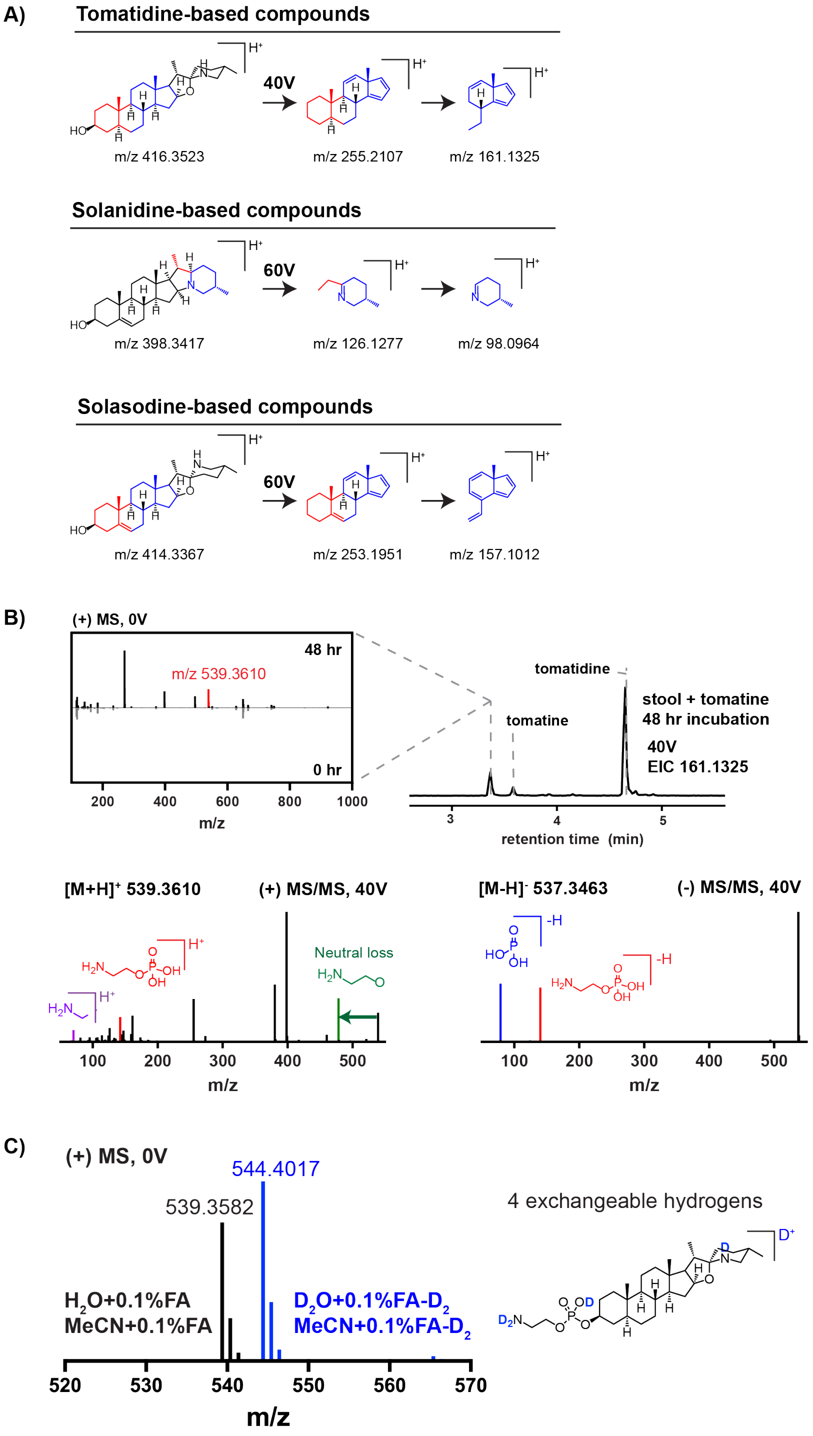


**Supplemental Figure 6. *Ex vivo* metabolism of SGAs with human stool samples. Relates to Figure 3.**

A) The characteristic fragments of tomatidine, solanidine, and solasodine generated by fragmentation at 40V, 60V, and 60V, respectively, used to identify steroidal alkaloid metabolites produced by human stool samples. Structures drawn according to proposed fragments (Cahill et al., 2010; Caprioli et al., 2015).

B) Identification of phosphoethanolamine (PEA) conjugated tomatidine as a product of tomatine metabolism by human stool samples. MS fragmentation of stool samples following incubation with tomatine at 40V revealed a peak that contained characteristic fragments of tomatidine (fragment *m/z* 161.1325). Comparing the MS spectra at this retention time of samples before and after incubation showed a new mass feature with $[M+H]^+$ *m/z* 539.3610. MS/MS fragmentation of this new metabolite in positive ($[M+H]^+$ *m/z* 539.3610) and negative modes ($[M-H]^-$ *m/z* 537.3463) led us to propose a phosphoethanolamine containing structure.

C) Hydrogen-deuterium exchange LC-MS to provide further support for the structure of PEA-conjugated tomatidine. Human stool incubated with tomatidine and containing the $[M+H]^+$ *m/z* 539.3710 mass was re-analyzed using deuterated chromatography solvents deuterium oxide ($D_2O$) and acetonitrile (MeCN) supplemented with 0.1% deuterated formic acid (FA-$D_2$). Positive mode MS analysis using deuterated solvents yielded a feature with $[M+D]^+$ *m/z* 544.4017, suggestive of a structurex with 4 exchangeable hydrogens.

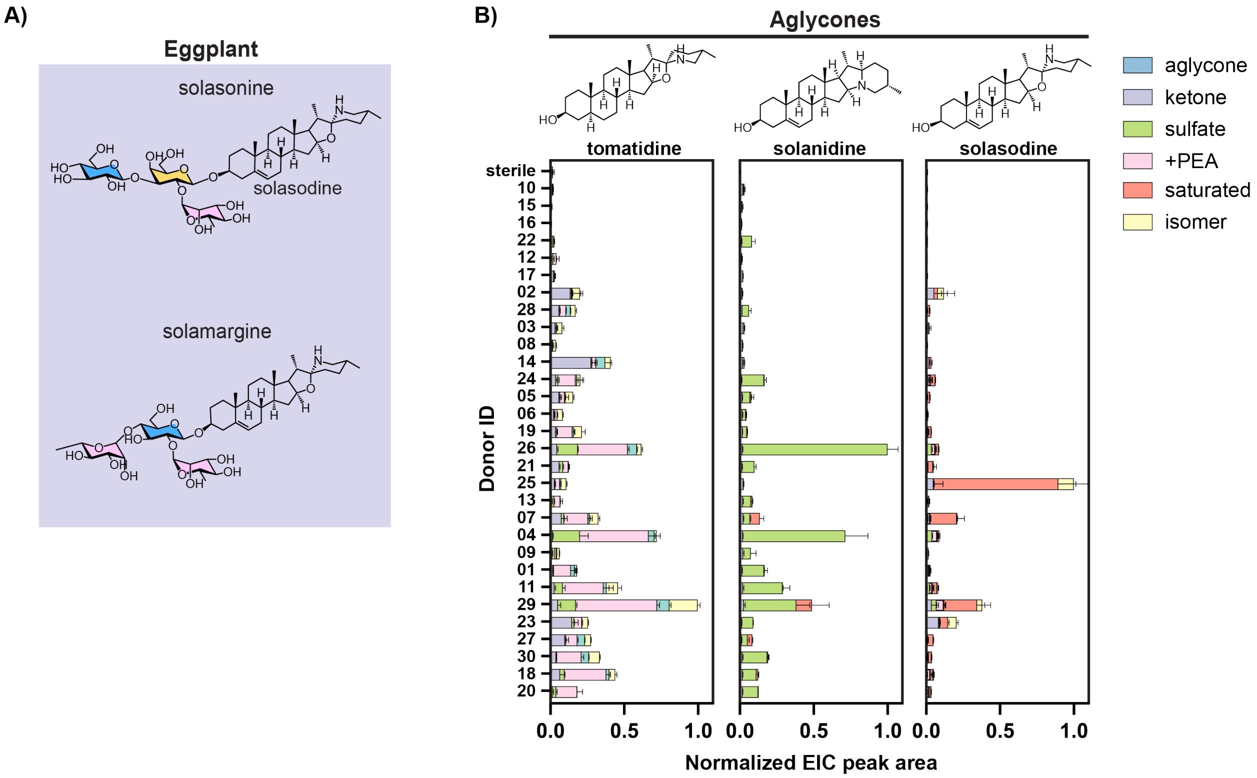


**Supplemental Figure 7. *Ex vivo* metabolism of eggplant SGAs and steroidal alkaloid aglycones by human stool samples. Relates to Figure 3.**

A) Major SGAs accumulating in eggplant.

B) *Ex vivo* metabolism of tomato, potato, and eggplant aglycones (tomatidine, solanidine, and solasodine, respectively) by human stool samples. Stool samples were directly incubated with aglycone (25 µM) in SAAC minimal medium for 48 hours. EIC peak areas of modified products were normalized to the maximum sum of product peak areas registered across all donors. Values shown are the mean±SD of three replicate incubations per donor and substrate combination.

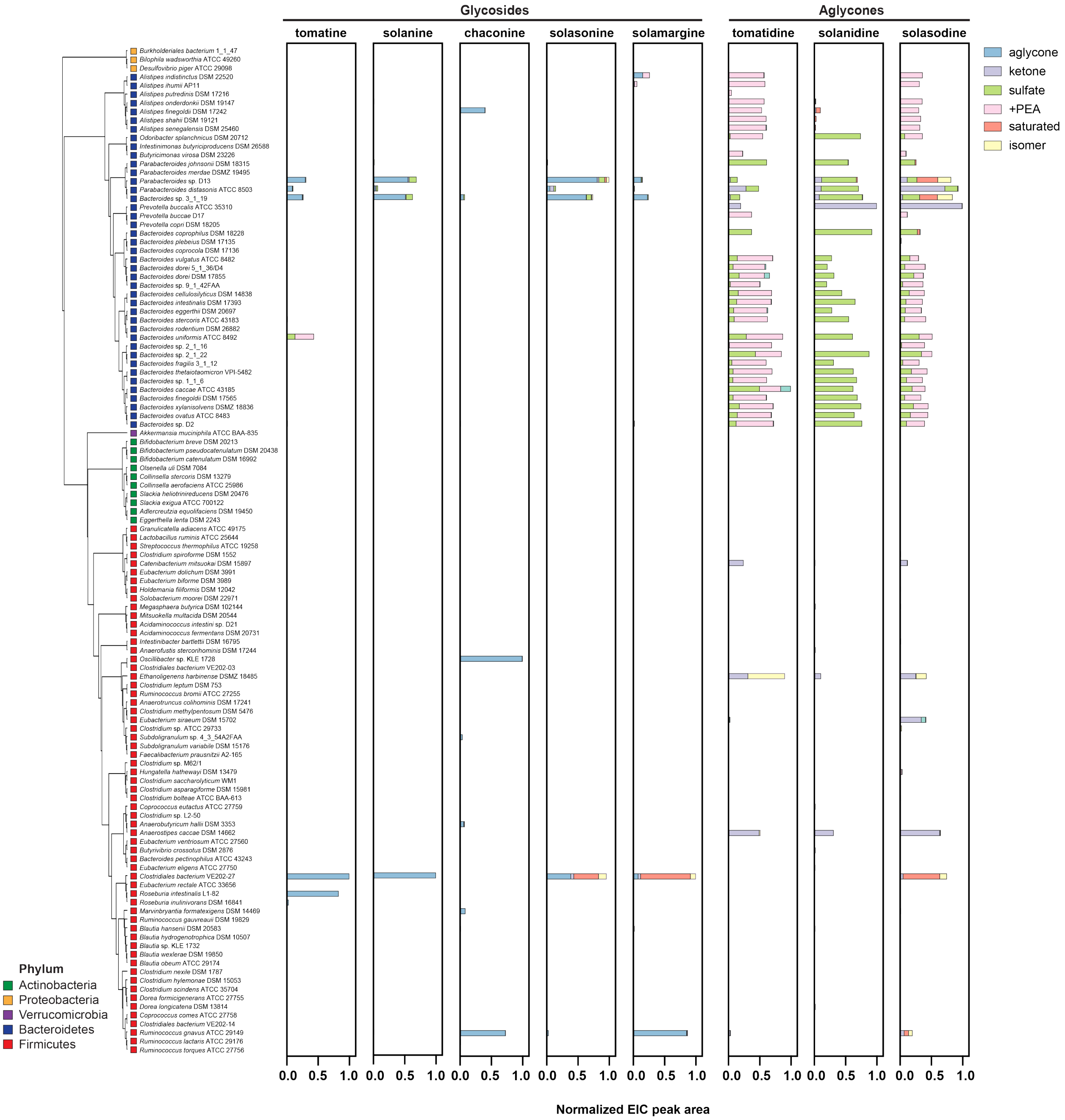


**Supplemental Figure 8. Metabolism of steroidal alkaloid aglycones by human commensal type strains. Relates to Figure 4.**

A complete, uncollapsed phylogenetic tree of the 116 human commensal type strains surveyed for steroidal alkaloid aglycone metabolism. Colored boxes next to strain names indicate phylum. Bars indicate the products of steroidal alkaloid aglycone metabolism by human commensal type strains after 48 hours of growth in Mega medium supplemented with steroidal alkaloid substrate (25 µM). EIC peak areas of the product aglycones are normalized to the maximum sum of product peak areas registered across all strains. Colors indicate different aglycone products. Data shown are representative of two repeat experiments.

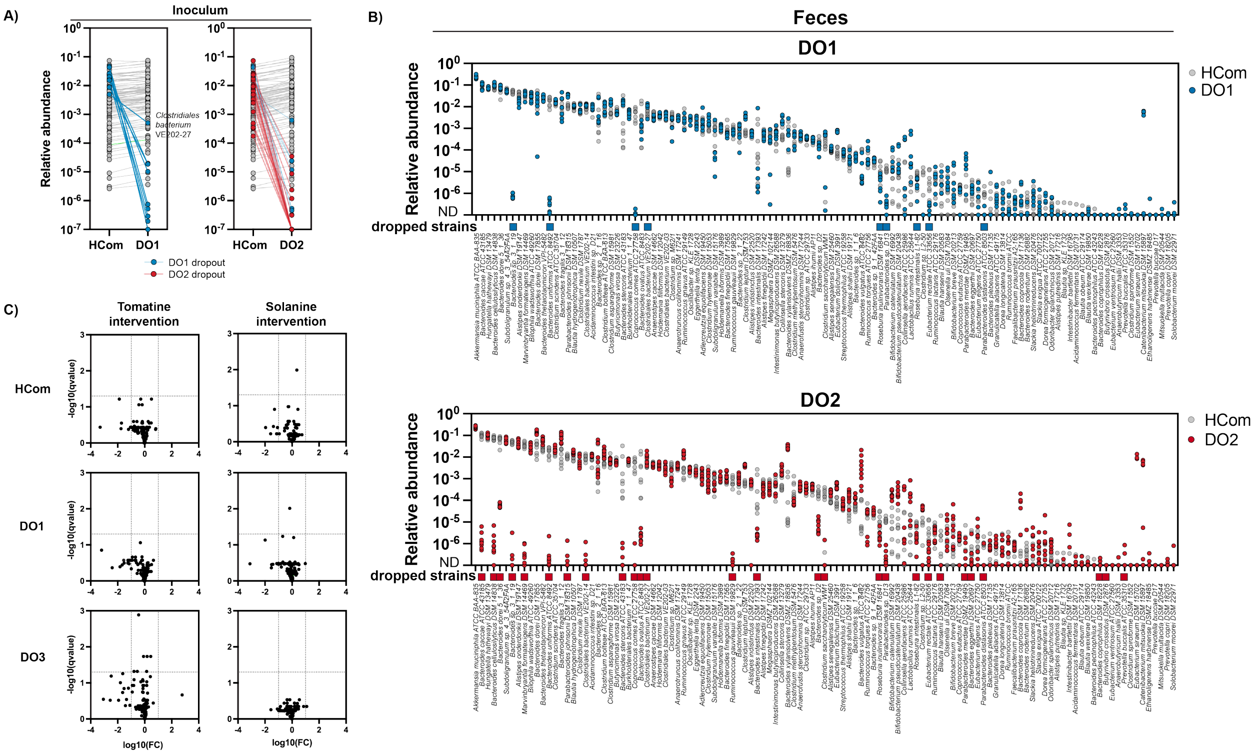


**Supplemental Figure 9. Metagenomic sequencing of gnotobiotic C57BL/6 mice engrafted with type strain communities. Relates to Figure 4.**

A) Relative abundances of strains in the pooled inoculum delivered to germ-free mice. Differences in strain abundance between the full community HCom (containing all 116 type strains surveyed for metabolism) and the SGA metabolism dropout communities DO1 and DO2 (containing 107- and 88- strain subsets of HCom, respectively) are highlighted in blue and red. Undetectable strains were set to a relative abundance of $10^{-7}$. While all dropped strains are lower abundance in their respective dropout communities relative to HCom, some maintain detectable representation in the dropout inoculum (e.g. *Clostridiales bacterium* VE202-27 highlighted).

B) Relative abundances of engrafted strains in feces collected from HCom (grey), DO1 (blue), and DO2 (red) colonized mice two weeks following colonization and prior to any dietary intervention. Values are shown for individual biological replicates (n=11 or 12). Strains are ordered from left to right by descending abundance in HCom-colonized samples. Strains from HCom that were excluded from DO1 and DO2 are indicated using colored boxes.

C) Volcano plots representing changes in type strain relative abundance after two weeks of dietary intervention with tomatine or solanine. No strains in HCom, DO1, or DO2 showed significant differences in relative abundance due to SGA intervention, defined as false discovery rate (FDR) <0.05 and fold change >10.

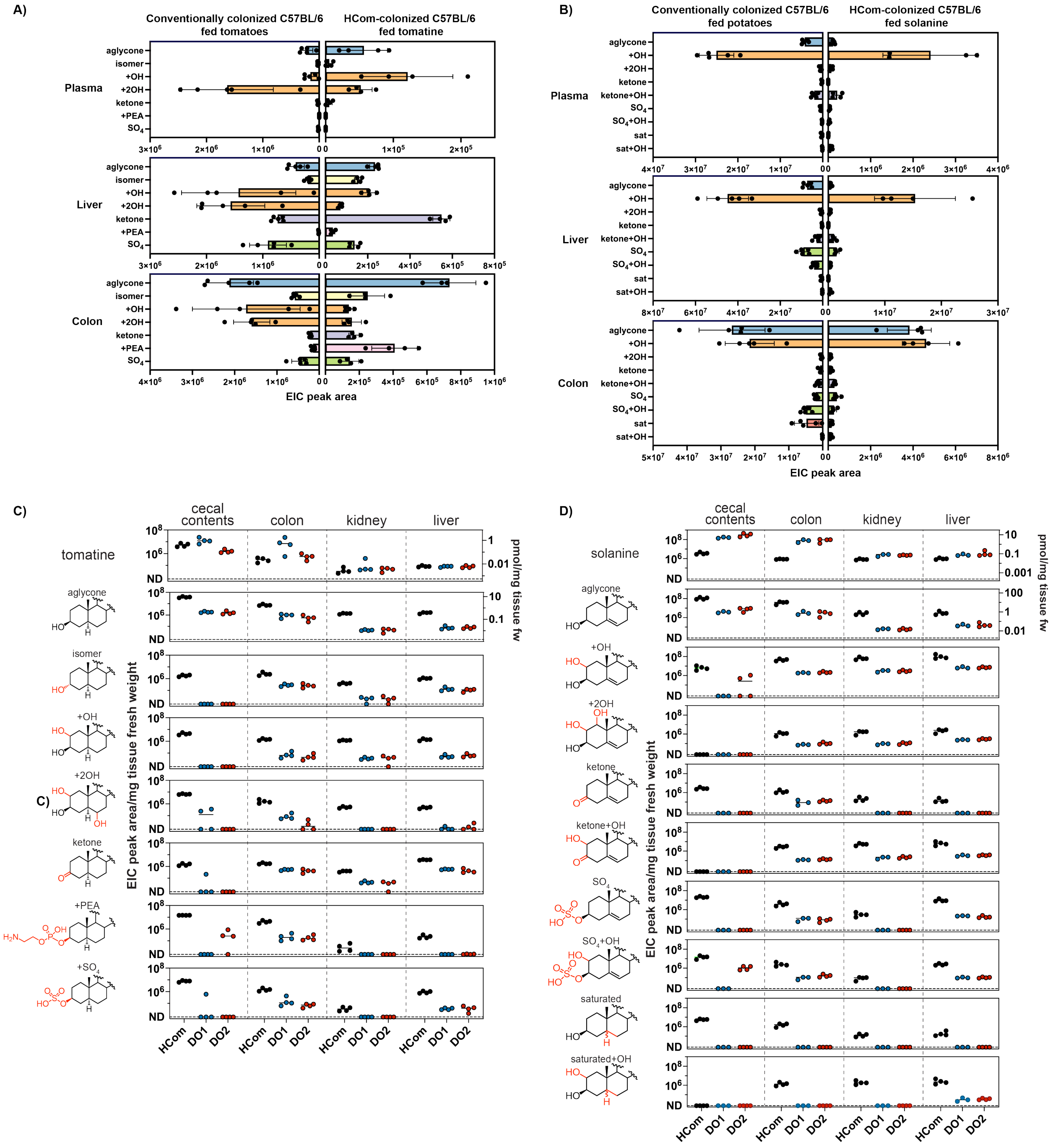


**Supplemental Figure 10. Community composition determines SGA metabolic fate. Relates to Figure 4.**

A) Comparison of the metabolic fate of tomatine in conventional C57BL/6 mice and HCom-colonized C57BL/6 mice. Major metabolites of tomatine were monitored in plasma, liver, and colon of conventional mice receiving tomato (Figure 2) and HCom-colonized mice receiving pure tomatine. Data shown are the mean±SD of n=5 conventional mice and n=4 gnotobiotic mice with individual replicates overlaid.

B) Comparison of the metabolic fate of solanine in conventional C57BL/6 mice and HCom-colonized C57BL/6 mice. Major metabolites of solanine were monitored in plasma, liver, and colon of conventional mice receiving potato (Figure 2) and HCom-colonized mice receiving pure solanine. Data shown are the mean±SD of n=5 conventional mice and n=4 gnotobiotic mice with individual replicates overlaid.

C) Accumulation of tomatine-derived metabolites in tissues of gnotobiotic C57BL/6J mice colonized with HCom (black), DO1 (blue), or DO2 (red). Metabolite EIC peak areas normalized to tissue fresh weight are represented for individual animals (n=4) with the mean value for each group indicated by a line. Molar quantities for compounds with commercial standards are indicated on the right axes where applicable. For molecules with multiple detected isomers (+OH and +2OH aglycones), the sum of peak areas is shown. ND = not detectable.

D) Accumulation of solanine-derived metabolites in tissues of gnotobiotic C57BL/6J mice colonized with HCom, DO1, or DO2. Metabolite EIC peak areas normalized to tissue fresh weight are represented for individual animals (n=3 or 4) with the mean value for each group indicated by a line. Molar quantities for compounds with commercial standards are indicated on the right axes where applicable. For molecules with multiple detected isomers (+OH and +2OH aglycones), the sum of peak areas is shown. ND = not detectable.

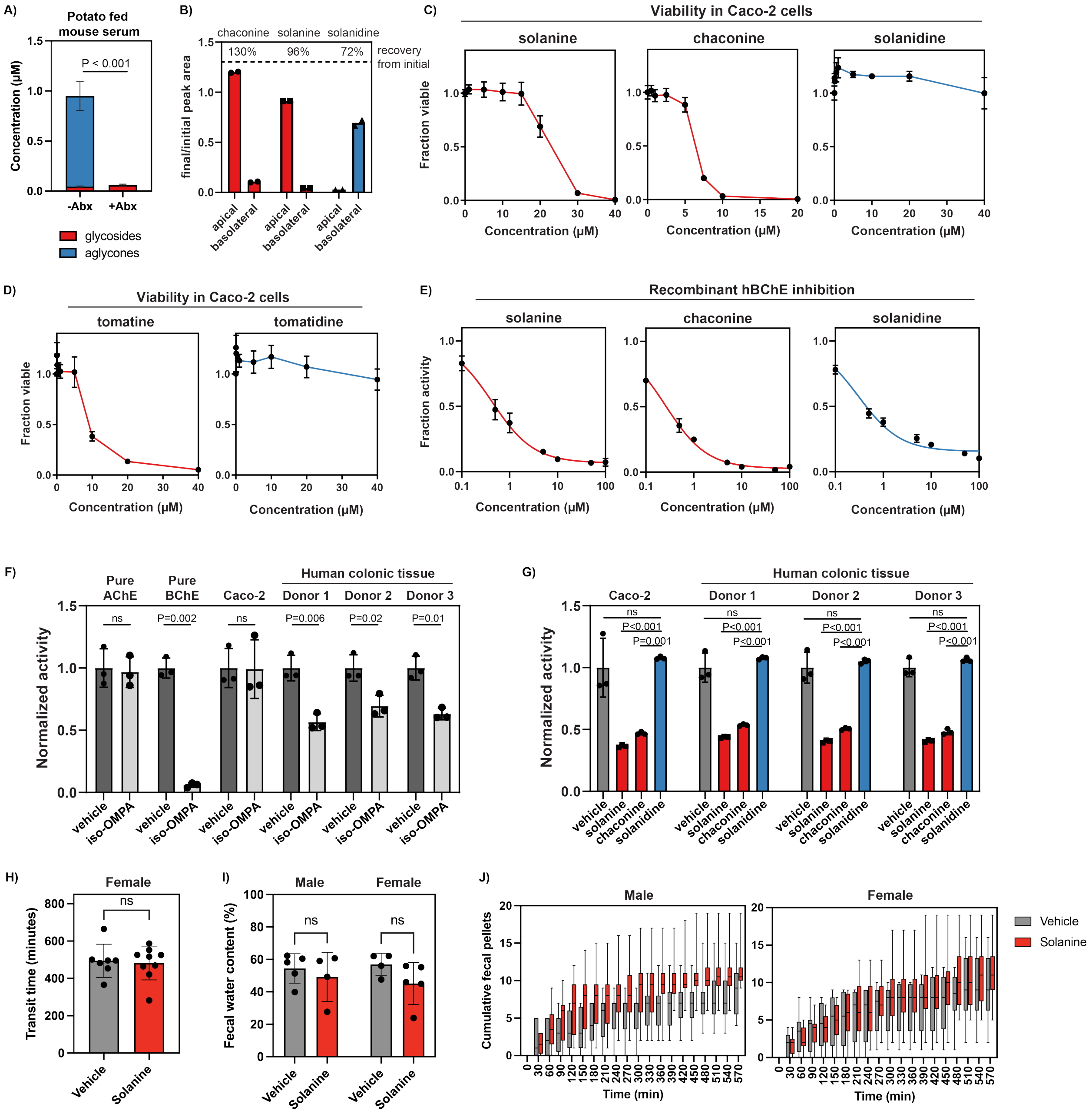


**Supplemental Figure 11. Effects of SGA glycosylation on compound bioavailability and bioactivity. Relates to Figure 4**.

A) Total steroidal alkaloid content in the serum of C57BL/6J mice fed potato-supplemented diet, with (+Abx) or without (-Abx) antibiotics treatment. Total steroidal alkaloid content was estimated as the sum of solanine, chaconine, and aglycones in the serum. Data for one of the steroidal alkaloid aglycones solanidine are shown in **Fig 2D**. Concentrations of modified solanidine compounds were estimated using a solanidine standard curve. Data shown are the mean±SD of five replicates.

B) Chaconine, solanine, and solanidine transport across a differentiated Caco-2 monolayer *in vitro*. Compounds (10 μM solanine, 3 μM chaconine, and 10 μM solanidine) were introduced into the apical side of the monolayer. EIC peak areas in each side of the monolayer following incubation were normalized by compartment volumes and divided by the initial EIC peak area of compound provided in media. The relative amount of initial compound recovered after incubation was calculated as the sum of EIC peak areas in the basolateral and apical sides after incubation, divided by the EIC peak area in the apical side before incubation. Values shown are the mean of two replicate incubations, with individual replicates overlaid.

C) Viability of undifferentiated Caco-2 cells treated with potato SAs. Cells were treated with compound dissolved in vehicle (DMSO for solanine, ethanol for chaconine and solanidine) for 24 hours. The total amount of vehicle included in each treatment was kept constant at 0.4% v/v. Viability readings were normalized to the viability readings of cells treated with vehicle. Data shown is the mean±SD (n=3 replicates).

D) Viability of undifferentiated Caco-2 cells treated with tomatine and tomatidine. Cells were treated with compound dissolved in vehicle (DMSO for tomatine, methanol for tomatidine) for 24 hours.

E) *In vitro* inhibition of recombinant human butyrylcholinesterase activity by potato SAs. Data shown are normalized to vehicle-treated BChE (DMSO for solanine and chaconine, ethanol for solanidine) with a total vehicle concentration of 1% v/v for all treatments. Values shown are the mean±SD of three replicates. 5 ng of purified protein were used in each reaction.

F) Acetylthiocholine hydrolysis by pure AChE, BChE, and colonic protein homogenates from Caco-2 epithelial cells and normal tissue from three donors with 500 µM of BChE selective inhibitor iso-OMPA. Activity was normalized to the activity of the sample treated with vehicle (DMSO) only. Values shown are the mean±SD of three replicates. 5 ng each of recombinant human AChE and BChE were used for pure protein reactions, while 40 µg total protein was used for each homogenate.

G) Inhibition of AChE by 50 µM potato steroidal alkaloids in human protein homogenates from normal colonic tissue (three donors) and Caco-2 colonic epithelial cells in the presence of BChE inhibitor iso-OMPA. Activity was normalized to the activity of the iso-OMPA only treatment for that respective sample. Values shown are the mean±SD of three replicates. 35 µg total protein was used for each homogenate.

H) Whole gut transit time in female Swiss Webster mice receiving 50 mg/kg body weight solanine. Values shown are the mean±SD for each group, with individual replicates overlaid (n=7 or 9). P values determined using a two-sided t test. ns = not significant (P>0.05).

I) Water content of fecal pellets collected following whole gut transit assay from male and female Swiss Webster mice receiving 50 mg/kg body weight solanine. Values shown are the mean±SD for each group, with individual replicates overlaid (n=4 or 5). P values determined using a two-sided unpaired t test. ns = not significant (P>0.05).

J) Cumulative pellet output during the whole gut transit assay by male and female Swiss Webster mice receiving 50 mg/kg body weight solanine. Boxes represent the 25 to 75$^{th}$ percentile of treatment groups of n=8 or 9 mice and whiskers represent the 10 to 90$^{th}$ percentiles.

#### Supplemental Table 1. Mass features tracked in this study

| Putative assignment | Formula | Level* | $[M+H]^+$ m/z | Column | Rt (min) |
|---|---|---|---|---|---|
| **Solanum meal associated compounds** | | | | | |
| Trigonelline | $C_7H_7NO_2$ | 1 | 138.0551 | HILIC | 5.30 |
| Trimethylamine N-oxide | $C_3H_9NO$ | 1 | 76.0757 | HILIC | 3.60 |
| Hippuric acid | $C_9H_9NO_3$ | 1 | 180.0655 | C18(10cm) | 4.40 |
| **Tomato Steroidal Alkaloids** | | | | | |
| Tomatine | $C_{50}H_{83}NO_{21}$ | 1 | 1034.5530 | C18(10cm) | 7.49 |
| Tomatidine | $C_{27}H_{45}NO_2$ | 1 | 416.3523 | C18(10cm) | 9.56 |
| 3a tomatidine | $C_{27}H_{45}NO_2$ | 2 | 416.3523 | C18(10cm) | 9.90 |
| 3-oxotomatidine | $C_{27}H_{43}NO_2$ | 1 | 414.3367 | C18(10cm) | 10.04 |
| Hydroxytomatidine 1 | $C_{27}H_{45}NO_3$ | 2 | 432.3460 | C18(10cm) | 8.44 |
| Hydroxytomatidine 2 | $C_{27}H_{45}NO_3$ | 2 | 432.3460 | C18(10cm) | 9.02 |
| Dihydroxytomatidine 1 | $C_{27}H_{45}NO_4$ | 2 | 448.3418 | C18(10cm) | 6.93 |
| Dihydroxytomatidine 2 | $C_{27}H_{45}NO_4$ | 2 | 448.3418 | C18(10cm) | 7.03 |
| Dihydroxytomatidine 3 | $C_{27}H_{45}NO_4$ | 2 | 448.3418 | C18(10cm) | 7.20 |
| Dihydroxytomatidine 4 | $C_{27}H_{45}NO_4$ | 2 | 448.3418 | C18(10cm) | 8.15 |
| Acetyhydroxytomatidine | $C_{29}H_{47}NO_4$ | 2 | 474.3554 | C18(10cm) | 9.58 |
| Tomatidine sulfate | $C_{27}H_{45}NO_5S$ | 1 | 496.3091 | C18(10cm) | 8.77 |
| Hydroxysulfotomatidine 1 | $C_{27}H_{45}NO_6S$ | 2 | 512.3040 | C18(10cm) | 7.76 |
| Hydroxysulfotomatidine 2 | $C_{27}H_{45}NO_6S$ | 2 | 512.3040 | C18(10cm) | 9.74 |
| Phosphoethanolamine tomatidine | $C_{29}H_{51}N_2O_5P$ | 2 | 539.3608 | C18(5cm) | 3.42 |
| Acetyltomatidine | $C_{27}H_{47}NO_3$ | 2 | 458.3629 | C18(5cm) | 4.85 |
| Tomatidine+ $C_2H_4O$ | $C_{29}H_{49}NO_3$ | 2 | 460.3789 | C18(5cm) | 4.46 |
| Esculeoside A | $C_{58}H_{95}NO_{29}$ | 2 | $[M+Na]^+$ 1292 | C18(10cm) | 6.25 |
| Esculeoside A–glc | $C_{52}H_{85}NO_{24}$ | 2 | 1108.5534 | C18(10cm) | 6.70 |
| Esculeoside A–lyc | $C_{35}H_{57}NO_{10}$ | 2 | 652.4055 | C18(10cm) | 7.12 |
| Esculeogenin A-1 | $C_{29}H_{47}NO_5$ | 2 | 490.3527 | C18(10cm) | 8.68 |
| Esculeogenin A-2 | $C_{29}H_{47}NO_5$ | 2 | 490.3527 | C18(10cm) | 9.75 |
| **Potato Steroidal Alkaloids** | | | | | |
| Solanine | $C_{45}H_{73}NO_{15}$ | 1 | 868.5053 | C18(10cm) | 7.46 |
| Chaconine | $C_{45}H_{73}NO_{14}$ | 1 | 852.5114 | C18(10cm) | 7.51 |
| Solanidine | $C_{27}H_{43}NO$ | 1 | 398.3423 | C18(10cm) | 9.06 |
| Solanine–glc/Chaconine–rham | $C_{39}H_{63}NO_{10}$ | 2 | 706.4523 | C18(10cm) | 7.55 |
| Solanidine plus hexose | $C_{33}H_{53}NO_6$ | 2 | 560.3946 | C18(10cm) | 7.76 |
| Hydroxysolanidine | $C_{27}H_{43}NO_2$ | 2 | 414.3367 | C18(10cm) | 8.43 |
| Dihydroxysolanidine | $C_{27}H_{43}NO_3$ | 2 | 430.3316 | C18(10cm) | 7.25 |
| Dihydrosolanidine 1 | $C_{27}H_{45}NO$ | 1 | 400.3574 | C18(10cm) | 9.26 |
| Dihydrosolanidine 2 | $C_{27}H_{45}NO$ | 2 | 400.3574 | C18(10cm) | 9.37 |
| Sulfosolanidine 1 | $C_{27}H_{43}NO_4S$ | 2 | 478.2986 | C18(10cm) | 8.84 |
| Sulfosolanidine 2 | $C_{27}H_{43}NO_4S$ | 2 | 478.2986 | C18(10cm) | 8.96 |
| Hydroxysulfosolanidine | $C_{27}H_{43}NO_5S$ | 2 | 494.2935 | C18(10cm) | 8.26 |
| 3-oxosolanidine | $C_{27}H_{41}NO$ | 2 | 396.3267 | C18(10cm) | 8.96 |
| Hydroxy-oxosolanidine | $C_{27}H_{41}NO_2$ | 2 | 412.3211 | C18(10cm) | 8.16 |
| Solanidine isomer | $C_{27}H_{43}NO$ | 2 | 398.3423 | C18(10cm) | 9.52 |

*Level of confidence in structural characterization:
1 = retention time, m/z, and MS/MS match to chemical standard
2 = identity proposed based on m/z, MS/MS, and retention time relative to similar structures and transformations confirmed by chemical standard

Glc = glucose, rham = rhamnose, lyc = lycotetraose

**Supplemental Table 2. Human commensal type strains used in this study**

All strains were provided by Microbiome Therapies Initiative, Sarafan Chem-H, Stanford University

| Strain Name | Growth medium |
|---|---|
| *Alistipes putredinis* DSM 17216 | YCFAM |
| *Anaerotruncus colihominis* DSM 17241 | Modified Columbia |
| *Bacteroides caccae* ATCC 43185 | YCFAC |
| *Bacteroides coprophilus* DSM 18228 | YCFAC |
| *Bacteroides dorei* 5_1_36/D4 | YCFAC |
| *Bacteroides eggerthii* DSM 20697 | YCFAC |
| *Bacteroides finegoldii* DSM 17565 | YCFAC |
| *Bacteroides fragilis* 3_1_12 | YCFAC |
| *Bacteroides intestinalis* DSM 17393 | YCFAC |
| *Bacteroides* sp. 1_1_6 | Modified Columbia |
| *Bacteroides* sp. 2_1_22 | Modified Columbia |
| *Bacteroides* sp. 3_1_19 | YCFAC |
| *Bacteroides* sp. 9_1_42FAA | YCFAC |
| *Bacteroides* sp. 2_1_16 | Modified Columbia |
| *Bacteroides* sp. D2 | YCFAC |
| *Bacteroides thetaiotaomicron* VPI-5482 | YCFAC |
| *Bacteroides xylanisolvens* DSMZ 18836 | YCFAC |
| *Bacteroides uniformis* ATCC 8492 | YCFAC |
| *Bacteroides pectinophilus* ATCC 43243 | Modified Columbia+5g/L sodium acetate, fucose, arginine, sodium gluconate, rhamnose, taurine |
| *Bacteroides plebeius* DSM 17135 | Modified PYG |
| *Bacteroides coprocola* DSM 17136 | BHI |
| *Bacteroides stercoris* ATCC 43183 | YCFAC |
| *Coprococcus eutactus* ATCC 27759 | Modified Columbia |
| *Eubacterium dolichum* DSM 3991 | Modified PYG |
| *Ruminococcus gnavus* ATCC 29149 | Modified Columbia |
| *Eubacterium rectale* ATCC 33656 | Modified Columbia |
| *Clostridium methylpentosum* DSM 5476 | Modified Columbia+5g/L sodium acetate, fucose, arginine, sodium gluconate, rhamnose, taurine |
| *Clostridium nexile* DSM 1787 | YCFAC |
| *Clostridium scindens* ATCC 35704 | Modified Columbia |
| *Clostridium* sp. L2-50 | Modified Columbia |
| *Clostridium* sp. M62/1 | YCFAC |
| *Clostridium asparagiforme* DSM 15981 | YCFAC |
| *Clostridium bolteae* ATCC BAA-613 | YCFAC |
| *Hungatella hathewayi* DSM 13479 | YCFAC |
| *Clostridium leptum* DSM 753 | YCFAC |

| | |
|---|---|
| *Dorea formicigenerans* ATCC 27755 | Modified Columbia |
| *Dorea longicatena* DSM 13814 | YCFAC |
| *Coprococcus comes* ATCC 27758 | Modified Columbia |
| *Blautia hansenii* DSM 20583 | Modified PYG |
| *Marvinbryantia formatexigens* DSM 14469 | Modified Columbia |
| *Butyrivibrio crossotus* DSM 2876 | mGAM |
| *Ruminococcus torques* ATCC 27756 | Modified PYG |
| *Parabacteroides merdae* DSMZ 19495 | Modified Columbia |
| *Subdoligranulum variabile* DSM 15176 | Wilkins-Chalgren no glucose |
| *Parabacteroides johnsonii* DSM 18315 | Modified Columbia |
| *Roseburia intestinalis* L1-82 (DSM 14610) | YCFAM |
| *Blautia obeum* ATCC 29174 | Modified PYG |
| *Eubacterium ventriosum* ATCC 27560 | YCFAC |
| *Faecalibacterium prausnitzii* A2-165 | YCFAM |
| *Parabacteroides* sp. D13 | YCFAC |
| *Anaerobutyricum hallii* DSM 3353 | Modified Columbia+5g/L sodium acetate, fucose, arginine, sodium gluconate, rhamnose, taurine |
| *Roseburia inulinivorans* DSM 16841 | Modified Columbia+5g/L sodium acetate, fucose, arginine, sodium gluconate, rhamnose, taurine |
| *Prevotella buccalis* ATCC 35310 | YCFAC |
| *Ruminococcus lactaris* ATCC 29176 | Modified Columbia |
| *Eubacterium eligens* ATCC 27750 | YCFAC |
| *Holdemania filiformis* DSM 12042 | Modified PYG |
| *Bacteroides ovatus* ATCC 8483 | Modified Columbia |
| *Bacteroides vulgatus* ATCC 8482 | YCFAC |
| *Clostridium spiroforme* DSM 1552 | Modified Columbia |
| *Holdemanella biformis* DSM 3989 | Modified PYG |
| *Blautia hydrogenotrophica* DSM 10507 | YCFAC |
| *Clostridium saccharolyticum* WM1 | YCFAC |
| *Parabacteroides distasonis* ATCC 8503 | YCFAC |
| *Eubacterium siraeum* DSM 15702 | YCFAC |
| *Eggerthella lenta* DSM 2243 | Modified Columbia+5g/L sodium acetate, fucose, arginine, sodium gluconate, rhamnose, taurine |
| *Anaerostipes caccae* DSM 14662 | YCFAC |
| *Bacteroides cellulosilyticus* DSM 14838 | Modified Columbia |
| *Clostridium hylemonae* DSM 15053 | YCFAC |
| *Acidaminococcus intestini* sp. D21 | YCFAC |
| *Catenibacterium mitsuokai* DSM 15897 | Modified Columbia |
| *Collinsella aerofaciens* ATCC 25986 | Modified PYG |
| *Acidaminococcus fermentans* DSM 20731 | Modified Columbia |
| *Intestinibacter bartlettii* DSM 16795 | Modified PYG |
| *Ethanoligenens harbinense* DSMZ 18485 | Ethanoligenes medium |

| | |
|---|---|
| *Collinsella stercoris* DSM 13279 | Modified PYG |
| *Prevotella buccae* D17 | YCFAC |
| *Mitsuokella multacida* DSM 20544 | Modified Columbia |
| *Olsenella uli* DSM 7084 | Modified PYG |
| *Slackia heliotrinireducens* DSM 20476 | Modified Columbia+5g/L sodium acetate, fucose, arginine, sodium gluconate, rhamnose, taurine |
| *Prevotella copri* DSM 18205 | YCFAC |
| *Slackia exigua* ATCC 700122 | Modified PYG |
| *Streptococcus thermophilus* ATCC 19258 | YCFAM |
| *Desulfovibrio piger* ATCC 29098 | TSB+5% blood |
| *Lactobacillus ruminis* ATCC 25644 | Modified Columbia |
| *Akkermansia muciniphila* ATCC BAA-835 | YCFAM |
| *Bifidobacterium pseudocatenulatum* DSM 20438 | YCFAC |
| *Solobacterium moorei* DSM 22971 | Modified PYG |
| *Anaerofustis stercorihominis* DSM 17244 | Bifidobacterium medium |
| *Granulicatella adiacens* ATCC 49175 | Modified Columbia |
| *Bacteroides dorei* DSM 17855 | YCFAC |
| *Bifidobacterium catenulatum* DSM 16992 | YCFAC |
| *Ruminococcus bromii* ATCC 27255 | YCFAC |
| *Bifidobacterium breve* DSM 20213 | YCFAC |
| *Megasphaera* DSMZ 102144 | YCFAC |
| *Clostridiales bacterium* VE202-03 | Modified Columbia |
| *Clostridiales bacterium* VE202-14 | YCFAC |
| *Clostridiales bacterium* VE202-27 | YCFAC |
| *Oscillibacter* sp. KLE 1728 | YCFAC |
| *Blautia* sp. KLE 1732 | Modified Columbia |
| *Alistipes shahii* DSM 19121 | Modified Columbia |
| *Alistipes onderdonkii* DSM 19147 | YCFAM |
| *Alistipes indistinctus* DSM 22520 | YCFAC |
| *Alistipes finegoldii* DSM 17242 | YCFAM |
| *Adlercreutzia equolifaciens* DSM 19450 | Wilkens-Chalgren with glucose |
| *Clostridium* sp. ATCC 29733 | mGAM |
| *Bilophila wadsworthia* ATCC 49260 | Modified Columbia+5g/L sodium acetate, fucose, arginine, sodium gluconate, rhamnose, taurine |
| *Alistipes ihumii* AP11 | Modified Columbia |
| *Subdoligranulum* sp. 4_3_54A2FAA | Chopped meat |
| *Alistipes senegalensis* DSM 25460 | YCFAM |
| *Blautia wexlerae* DSM 19850 | Modified Columbia |
| *Butyricimonas virosa* DSM 23226 | YCFAC |
| *Intestinimonas butyriciproducens* DSM 26588 | Modified Columbia |
| *Odoribacter splanchnicus* DSM 20712 | Modified Columbia |

| | |
|---|---|
| *Ruminococcus gauvreauii* DSM 19829 | mGAM |
| *Bacteroides rodentium* DSM 26882 | Modified Columbia |
| *Burkholderiales bacterium* 1_1_47 | Modified PYG + 3g/L sodium formate, sodium fumarate |

#### Supplemental Discussion

##### Metabolism of SGAs by environmental and plant-colonizing microbes

To date, the bacterial strains found to hydrolyze SGAs have been environmental microbes. Tomato pathogens, including the fungus *Septoria lycopersici* (Osbourn et al., 1995) and the bacterial wilt effector *Clavibacter michiganensis* (Kaup et al., 2005), have developed strategies for tomatine deglycosylation to counteract the membrane disrupting capabilities imparted by the lycotetraose glycone. Similarly, soil bacteria from the genus *Arthrobacter* (Hennessy et al., 2018) and *Glutamicibacter halophytocola* S2, a commensal bacteria colonizing the potato tuber moth (Wang et al., 2022), have been shown to metabolize solanine and chaconine from potato. Beyond environmental sources, bovine rumen has been shown to metabolize potato SGAs (King and McQueen, 1981).

##### Hydrogenation of Δ5,6 SA aglycones

A major transformation that we observed in both mouse systemic circulation and in human urine involved hydrogenation of the Δ5,6 double bond of the potato aglycone solanidine. A previous report of such a compound noted its formation in bovine rumen fluid from a solanine input (King and McQueen, 1981). More broadly, however, this reduction is reminiscent of gut microbiota-mediated hydrogenation of cholesterol to coprostanol. Bacteria identified to be coprostanol producers include *Bacteroides* sp. D8 (Gérard et al., 2007) and strains from the genus *Eubacterium* (Eyssen et al., 1973; Freier et al., 1994). In human gut communities, the cholesterol dehydrogenase ismA from uncultured Clostridia has been implicated in coprostanol formation (Kenny et al., 2020). Despite the analogous transformations observed for solanidine and cholesterol, however, we observed solanidine hydrogenation to be performed by Bacteroidetes, including *Alistipes finegoldii* DSM 17242 (**Fig S8**). A BLAST search revealed no close homologs for *ismA* homologs in solanidine reducing type strains. As such, we hypothesize that solanidine hydrogenation proceeds by a route distinct to the one identified for coprostanol formation.

##### Sulfonation of SA aglycones

We found steroidal alkaloid sulfates to significantly accumulate in host liver following dietary intervention with tomatine and potato. Steroids like cholesterol and dehydroepiandrosterone are well-established substrates for mammalian sulfotransferases (Falany et al., 1994). Potentially resulting from promiscuous activity by these sulfotransferases, analogous sulfation of tomatidine and solanidine was performed by human hepatic proteins. Beyond formation by host enzymes, we also noted contributions by gut microbiota towards sulfated steroidal aglycones. Recent work has identified a gene cluster in *Bacteroides* *thetaiotaomicron* VPI-5482 that sulfonates cholesterol and other steroidal compounds with a trans A/B ring fusion, a structural feature shared by tomatidine and solanidine (Yao et al., 2022). In agreement with the prevalence of this gene cluster in Bacteroidetes strains, our survey of human commensal strains revealed primarily Bacteroidetes and not strains from other phyla to be capable of generating sulfonated forms of tomatidine and solanidine. Future work will determine whether the identified bacterial sulfotransferases that act on cholesterol similarly sulfonate steroidal alkaloids from dietary Solanums.

**Phosphoethanolamine conjugation to SA aglycones**

One unexpected transformation of spirosolane type aglycones tomatidine and solasodine that we observed to be performed *ex vivo* by human stool samples was conjugation to a putative phosphoethanolamine moiety. Phosphoethanolamine conjugation to small molecules has been described in the context of sphingolipid biosynthesis, with Bacteroidetes being the only gut commensals known to produce these molecules (Johnson et al., 2020). The major species of sphingolipid made by Bacteroidetes is ceramide phosphoethanolamine and is putatively involved in mitigating environmental stress and interactions with host immunity (Panevska et al., 2019). With only Bacteroidetes being capable of phosphoethanolamine conjugation on both sphingolipids and steroidal alkaloid aglycones (**Fig. S8**), we speculate that PEA conjugation of SA substrates may be a result of promiscuous activity by the phosphoethanolamine transferases that natively act on sphingolipids. From these examples, we hypothesize that these steroidal alkaloids from dietary Solanums may be metabolized through host and bacterial pathways that intersect with those that process endogenous host steroids.

**Bioactivity associated with SA aglycones**

Beyond the depletion of bioactive SGAs by microbiota-mediated deglycosylation, the aglycone products of this SGA may have distinct physiological effects. For example, dietary intervention with the aglycone tomatidine was found to increase mTORC1 signaling in mice, resulting in increased skeletal muscle hypertrophy and decreased adiposity (Dyle et al., 2014). Further modified versions of steroidal aglycones produced by host or bacterial metabolism may also result in differential activity relative to the unmodified aglycone. For example, modification of the bile acid lithocholic acid by oxidation at the C3 alcohol, a transformation that we also observed with tomatidine and solanidine, enabled binding of the compound to host RORγt receptors and resulted in inhibition of $T_H17$ cell differentiation (Hang et al., 2019).

**Toxicity of potato SGAs**

Improper storage of potatoes can result in overaccumulation of SGAs in the tubers and has led to numerous reports of SGA-induced poisonings (McMillan and Thompson, J.C., 1979), with an estimated toxic dose of 2-5 mg/kg body weight in humans (Slanina, 1990). Acute symptoms of SGA toxicity commonly involve nausea, vomiting and diarrhea (Mensinga et al., 2005), indicative of effects local to the gastrointestinal tract. Moreover, treatment with the acetylcholine receptor inhibitor atropine was found to alleviate toxicity following of solanine administration in mice (Patil et al., 1972), suggestive of a role for cholinesterase inhibition in mediating toxicity.
